# MaRNAV-1, an intracellular virus of *Plasmodium vivax*, is associated with increased parasite transmission and altered host immune response

**DOI:** 10.64898/2026.03.22.713337

**Authors:** Dynang Seng, Katie Ko, Agnes Orban, Sokleap Heng, Lionel Brice Feufack-Donfack, Janne Grünebast, Nisa Ya, Franck Dumetz, Kieran Tebben, Tiziano Vignolini, Gregory Dore, Nimol Khim, Jeremy Salvador, Zainab Ouaid, Thierry Lefèvre, Anna Cohuet, Cecile Sommen, Claude Flamand, Anthony A. Ruberto, Abnet Abebe, Meshesha Tsigie, Geremew Tasew, Getachew Tollera, Benoit Malleret, Tassew Tefera Shenkutie, Eugenia Lo, Nicholas M Anstey, Mary E Petrone, Tineke Cantaert, Sebastian Baumgarten, David Serre, Jean Popovici

**Author notes:** Contributed equally (Katie Ko, Agnes Orban, Sokleap Heng).

## Abstract

MaRNAV-1 is an RNA virus recently identified in *Plasmodium vivax*-infected samples, but definitive evidence that it infects the parasite and influences malaria pathogenesis remains unknown.

Here, we demonstrate that MaRNAV-1 is an intracellular virus that is present in *P. vivax* at various stages of its life cycle, including blood, sporozoite, and liver stages. Viral prevalence varied geographically between Cambodian and Ethiopian parasites. MaRNAV-1 presence and load were positively associated with parasite transmission potential, as reflected by increased gametocyte abundance and higher oocyst prevalence and intensity in membrane feeding assays. MaRNAV-1 loads were higher in symptomatic compared to asymptomatic infections, and higher MaRNAV-1 loads were associated with elevated body temperature, independently of parasitemia. MaRNAV-1 infection elicits an antibody response and is associated with dendritic cell activation, a shift from Th2 to a Th1-driven immune response, and an increased frequency of double-negative B cells. Accordingly, MaRNAV-1-infected patients had higher concentrations of circulating cytokines, such as IFN-γ, CXCL10, IL-1RA, and IL-6, independently of parasitemia.

Together, these findings demonstrate that MaRNAV-1 is a genuine parasite-infecting virus associated with increased parasite transmission potential and with modulation of clinical outcomes in, and host immune response to, *P. vivax* infections. Our study broadens the conventional view of host-pathogen interactions in malaria by revealing complex virus-parasite-host relationships.

## Introduction

Viruses are ubiquitous biological entities capable of infecting all forms of life, from bacteria to eukaryotes, including parasitic protozoa. Viruses infecting parasites were first described several decades ago, when virus-like particles (VLPs) were observed in *Plasmodium*, *Entamoeba*, and *Leishmania* by transmission electron microscopy^1–3^. The first well-characterized protozoan-infecting virus was discovered in *Trichomonas vaginalis*, the causative agent of trichomoniasis^4,5^. Despite these early observations, viruses infecting *Plasmodium* species are still poorly characterized. Five *Plasmodium* species primarily cause human malaria: *P. falciparum*, *P. vivax*, *P. ovale*, *P. malariae*, and *P. knowlesi*. Of these, *P. vivax* is the most geographically widespread parasite and represents a major cause of malaria morbidity, particularly in Asia, South America, and the Horn of Africa^6^. Although historically considered responsible for relatively benign disease, increasing evidence indicates that *P. vivax* infections can cause severe complications and mortality^7,8^.

Recently, a bi-segmented narnavirus was identified by meta-transcriptomic analysis in *P. vivax-*infected blood samples^9^. This virus, termed Matryoshka RNA Virus 1 (MaRNAV-1), was detected across multiple RNA-seq datasets from *P. vivax-*infected blood samples. In contrast, it was absent in datasets derived from *P. falciparum, P. malariae, P. ovale,* and *P. knowlesi* infections^9,10^. More recently, an unrelated virus was reported in association with *P. knowlesi* infections^11^, suggesting that different *Plasmodium* species may harbor distinct viruses.

This association of MaRNAV-1 with *P. vivax* infections raises intriguing questions about the virus’s host range and its potential role in parasite biology and pathogenesis. Viruses infecting protozoan parasites can profoundly influence host-parasite interactions^12–18^. For example, double-stranded RNA viruses infecting *Leishmania* parasites have been shown to exacerbate disease severity by triggering inflammatory responses in the human host^19–21^. Additionally, viruses infecting *Leishmania*, *Trichomonas,* and *Giardia* can alter parasite virulence and treatment outcomes^22^. In other systems, viruses may reduce the pathogenicity of their host organism, a phenomenon known as hypovirulence, which has been described for several mitoviruses infecting fungi^23,24^. These examples illustrate how parasite-associated viruses can influence both parasite biology and host disease manifestations. The hypothesis that malaria parasites harbor viruses adds a previously unrecognized and largely unexplored layer of biological complexity to host-parasite interactions.

MaRNAV-1 belongs to the family *Narnaviridae*, one of the simplest groups of RNA viruses. Narnaviruses are non-enveloped viruses with compact single-stranded RNA genomes composed of one or two segments. Most known narnaviruses infect fungal hosts and replicate in the cytoplasm without forming classical virions^25,26^. Their transmission is thought to occur primarily through vertical inheritance during host cell division, allowing long-term persistence within host lineages, although horizontal transmission has also been reported^26,27^.

The genome of MaRNAV-1 comprises two RNA segments (Supp. Fig. 1). The first segment (S1) encodes an RNA-dependent RNA polymerase (RdRp) (GenBank accession No. MN860568.1) that shares conserved motifs with the polymerase of fungal narnaviruses^9^. The second segment (S2) encodes two overlapping open reading frames (ORFs, denoted as S2_ORF1 and S2_ORF2), possibly encoding proteins with no homology with annotated proteins (GenBank accession No. MN860569.1).

**Fig. 1.**
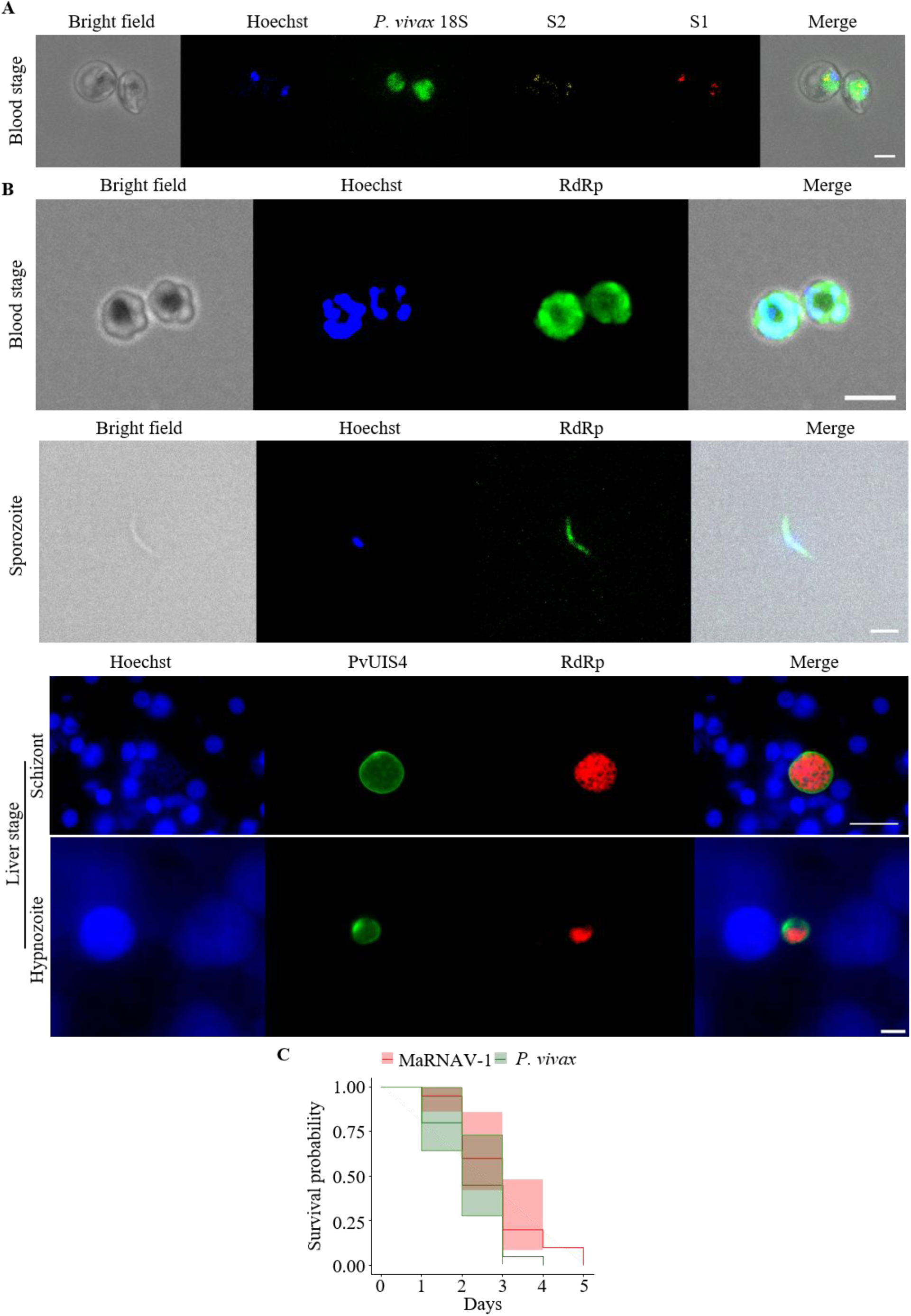
MaRNAV-1 is an intracellular virus of *P. vivax.* **A.** Detection of MaRNAV-1 RNA by smiFISH in blood stage *P. vivax* parasites. From left to right: bright field; DNA stained with Hoechst (blue); *P. vivax* 18S rRNA labeled with FLAP-Y-Alexa Fluor 488 (green); MaRNAV-1 S2 RNA labeled with FLAP-Y-Cy3 (yellow); MaRNAV-1 S1 RNA labeled with FLAP-Y-Cy5 (red); merged image. Viral RNA signals co-localize with parasite 18S rRNA within infected reticulocytes. Scale bar, 5 μm. **B**. Immunofluorescence localization of the MaRNAV-1 RNA-dependent RNA polymerase (RdRp) across *P. vivax* life cycle stages. From top to bottom: blood stage parasites, sporozoite, and liver stage parasites. DNA was stained with Hoechst (blue). In the blood stage and sporozoites, RdRp was detected using rabbit anti-RdRp antibodies followed by goat anti-rabbit Alexa Fluor 488 secondary antibodies (green). In liver stage parasites, the parasitophorous vacuole membrane was labeled using mouse anti-UIS4 monoclonal antibodies detected with goat anti-mouse Alexa Fluor 488 secondary antibodies (green), while RdRp protein was detected with goat anti-rabbit Alexa Fluor Plus 594 secondary antibodies (red). Scale bars: 5 μm (blood stage and sporozoite), 30 μm (schizont), and 5 μm (hypnozoite). **C.** Survival probability of MaRNAV-1 within *P. vivax*-infected patients (n = 20) during 7-day artesunate treatment. The shaded areas represent 95% confidence interval (CI).

Despite the recent discovery of the cooccurrence of MaRNAV-1 and *P. vivax* and the description of its wide geographic distribution^10^, it is unclear whether this virus truly infects the parasite (as opposed to coinfecting patients) and whether it influences parasite biology or host immune responses.

In this study, we characterize MaRNAV-1 in *P. vivax* infections. Using imaging approaches, population surveys of clinical isolates from Cambodia and Ethiopia, and analyses of parasite and host data, we examined the intracellular localization and potential biological consequences of this virus.

## Results

### MaRNAV-1 infects *P. vivax* cells

To determine whether MaRNAV-1 infects *P. vivax* parasites rather than represents a coinfection in patients, we employed three complementary approaches. First, we performed single-molecule inexpensive fluorescence *in situ* hybridization (smiFISH) to visualize viral RNA transcripts from both genomic segments (S1 and S2) in *P. vivax*-infected red blood cells from clinical samples that were positive for MaRNAV-1 by RT-qPCR. Confocal microscopy of blood stage parasites revealed distinct fluorescent signals for both S1 and S2 that co-localized with the parasite 18S rRNA, demonstrating intracellular localization of the virus within *P. vivax* (Fig. 1A). No viral RNA was detected in adjacent uninfected red blood cells (Supp. Fig. 2A).

**Fig. 2.**
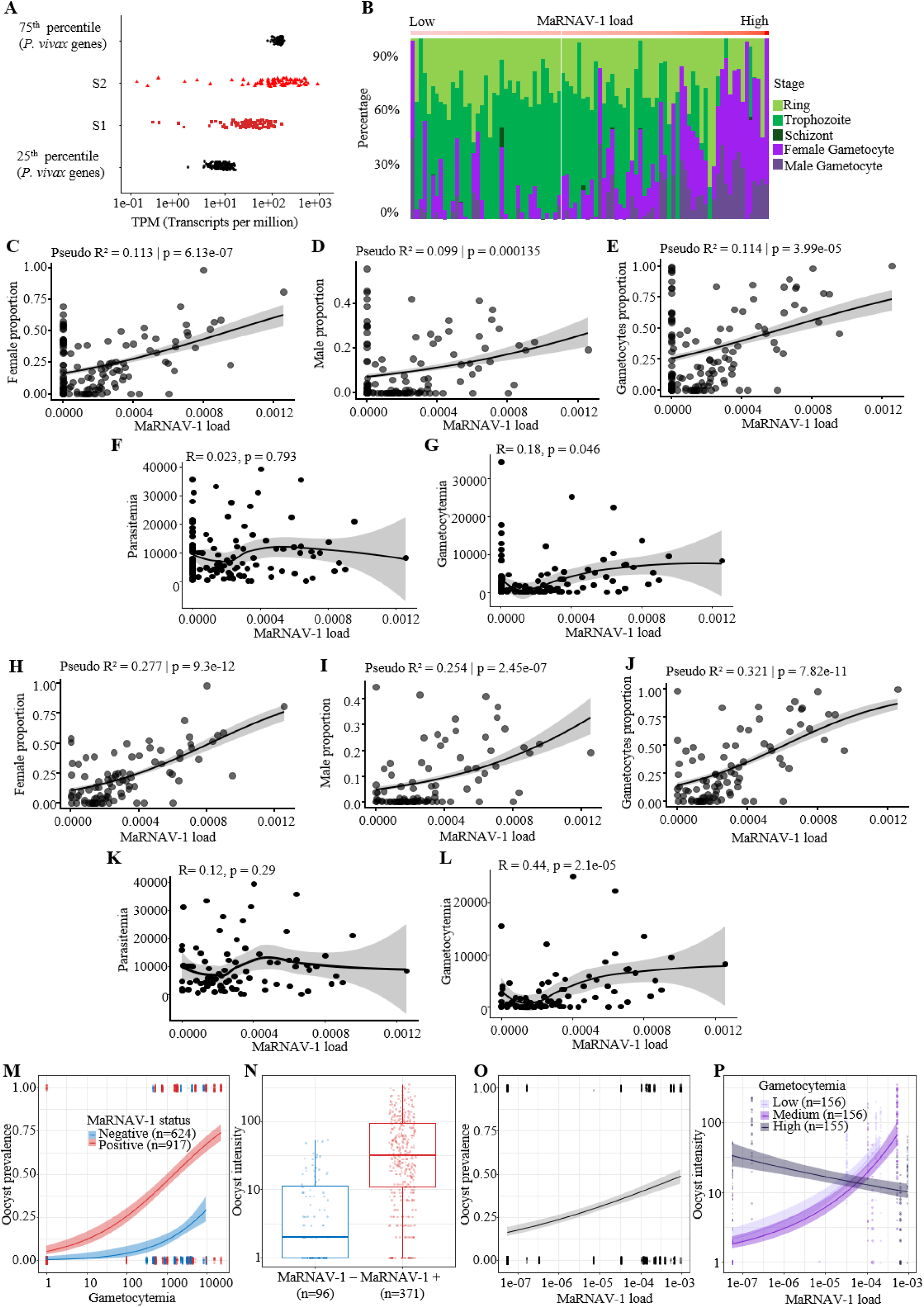
Association of MaRNAV-1 with mosquito transmission. **A.** Expression levels of MaRNAV-1 S1 and S2 (TPM, transcripts per million) are compared to the 25^th^ and 75^th^ percentiles of *P. vivax* gene expression within each sample. Each dot represents an individual patient sample (n = 126). **B**. Distribution of blood stage parasite composition in Cambodian *P. vivax* infections positive for MaRNAV-1, inferred by gene expression deconvolution. Each bar represents one sample, ordered by increasing MaRNAV-1 load (from low to high, n = 88). Colors indicate parasite developmental stages. **C-E.** Beta regression analyses assessing associations between MaRNAV-1 load and blood stage parasite composition, including both MaRNAV-1 negative and positive samples. There were significant positive associations between MaRNAV-1 load and (**C**) female, (**D**) male, and (**E**) total gametocytes proportions (Pseudo R^2^ = 0.113, p < 0.0001; Pseudo R^2^ = 0.099, p = 0.0001; Pseudo R^2^ = 0.114, p < 0.0001, respectively, n = 126). Lines represent fitted values from beta regression models with shaded 95% confidence intervals (CI). **F**. Spearman correlation between gametocytemia and MaRNAV-1 load by RNA-seq, including both MaRNAV-1 negative and positive samples (R = 0.18, p = 0.046, n = 125). **G**. No correlation between MaRNAV-1 load and parasitemia (R = 0.023, p = 0.793, n = 125). (**F-G**) The black line represents a LOESS-smoothed fit, with the shaded area indicating 95% CI. **H**-**J**. Beta regression analyses assessing associations between MaRNAV-1 load and blood stage parasite composition in MaRNAV-1-positive samples only (n = 88). The associations remained significant between MaRNAV-1 load and (**H**) female, (**I**) male, (**J**) total gametocytes proportions (Pseudo R^2^ = 0.277, p < 0.0001; Pseudo R^2^ = 0.254, p < 0.0001; Pseudo R^2^ = 0.321, p < 0.0001, respectively, n= 88). Lines represent fitted values from beta regression models with shaded 95% CI. **K**. Spearman correlation between gametocytemia and MaRNAV-1 load by RNA-seq, in MaRNAV-1-positive samples only (R = 0.44, p < 0.0001, n = 87). **L**. No correlation between parasitemia and MaRNAV-1 load (R = 0.12, p = 0.29, n = 87) among MaRNAV-1 positive samples. (**K-L**) The black line represents a LOESS-smoothed fit, with the shaded area indicating 95% CI. **M**. Relationship between gametocytemia and oocyst prevalence in mosquitoes following membrane feeding, stratified by MaRNAV-1 infection status (number of mosquitoes: n=624 MaRNAV-1-negative, and n=917 MaRNAV-1-positive). Lines represent fitted probabilities with 95% CI. **N.** Oocyst intensity (number of oocysts per infected mosquito) according to MaRNAV-1 infection status (number of infected mosquitoes: n = 96 MaRNAV-1-negative, and n = 371 MaRNAV-1-positive). Boxplots show the median and interquartile range, with individual data points overlaid. **O**. Relationship between virus load and oocyst prevalence. The fitted line shows the predicted probability of infection with 95% CI. **P.** Relationship between oocyst intensity and MaRNAV-1 load, stratified by gametocytemia levels (tertiles: low: 0-1581 parasites/µL; medium: 1581-5133 parasites/µL; high: 5133-22284 parasites/µL, number of infected mosquitoes n=155 or 156 per level). The fitted line represents the predictions with 95% CI. Across panels **M-P**, points represent individual mosquitoes. Continuous predictors are displayed on a log₁₀ scale.

Secondly, we used immunofluorescence assays (IFA) to localize the viral RdRp. The signal was detected in all examined stages of *P. vivax*, including blood stages, sporozoites, liver schizonts, and dormant hypnozoites (Fig. 1B). For both blood and liver stages, a fluorescent signal was detected only in *P. vivax-*infected cells and not in surrounding uninfected ones (Fig. 1B & Supp. Fig. 2B). Negative controls made of *P. falciparum* blood stages or using pre-immune IgGs showed no detectable signal (Supp. Fig. 2C & 2D). Together, these complementary imaging approaches provide direct evidence that MaRNAV-1 infects *P. vivax* cells and that viral proteins are expressed at different stages of the parasite life cycle.

Finally, in 20 patients with *P. vivax* infections positive for MaRNAV-1, we monitored the presence of viral and parasitic RNA by RT-qPCR over the course of antimalarial treatment with artesunate^28^. Over 4-5 days, both viral RNA and parasitic RNA were cleared with similar kinetics (Fig. 1C & Supp. Table 1). Importantly, the concomitant disappearance of viral RNA following parasite clearance further indicates that the virus does not persist independently in peripheral blood outside the parasite.

### MaRNAV-1 presence and load are positively correlated with gametocyte abundance and enhanced transmission to mosquito vectors

To understand the impact of MaRNAV-1 infection on parasite biology, we analyzed bulk RNA-seq data generated from whole-blood samples collected from 126 Cambodian patients seeking treatment for symptomatic *P. vivax* malaria and enrolled in a clinical trial^28^. Among these, 70% (88/126) were positive for MaRNAV-1, and virus transcript abundance spanned a wide range of virus loads (from 10^-7^ to 10^-3^, expressed as the ratio of MaRNAV-1 reads to reads mapping to *P. vivax*) (Supp. Data 1). The MaRNAV-1 transcripts were overall highly expressed. The S2 segment was consistently expressed at a higher level than the S1 segment (p < 0.0001), and often as abundant as the most highly expressed *P. vivax* genes (Fig. 2A & Supp. Data 2). Of note, RNA selection prior to RNA-seq library construction relied on polyA selection. It is unclear whether S1 and S2 are polyadenylated and, if they are not, RNA-seq may have underestimated MaRNAV-1 abundance.

Since *P. vivax* blood stage infections are often asynchronous and contain various proportions of developmental stages, we estimated the proportions of each stage (rings, trophozoites, schizonts, and male and female gametocytes) in every sample using gene expression deconvolution^29^. MaRNAV-1 was detected in samples with variable proportions of the different developmental stages (Fig. 2B). Interestingly, the virus load was positively correlated with the proportion of both female, male, and total gametocytes (Pseudo R^2^ = 0.113, p < 0.0001; Pseudo R^2^ = 0.099, p = 0.0001; Pseudo R^2^ = 0.114, p < 0.0001, respectively, Fig. 2C-E & Supp. Fig. 3). Consistent with this observation, while MaRNAV-1 load was not correlated with parasitemia (R = 0.023, p = 0.793, Supp. Data 1 & Fig. 2F), it was with absolute gametocytemia (R = 0.18, p = 0.046, Fig. 2G). When considering only MaRNAV-1 positive samples, correlations between virus load and gametocyte proportions (female, male and total) were more pronounced (Pseudo R^2^ = 0.277, p < 0.0001; Pseudo R^2^ = 0.254, p < 0.0001; Pseudo R^2^ = 0.321, p < 0.0001, respectively, Fig. 2H-J & Supp. Fig. 3), while virus load remained not correlated to parasitemia (R = 0.12, p = 0.29, Fig. 2K) and correlated to absolute gametocytemia (R = 0.44, p < 0.0001, Fig. 2L)

**Fig. 3.**
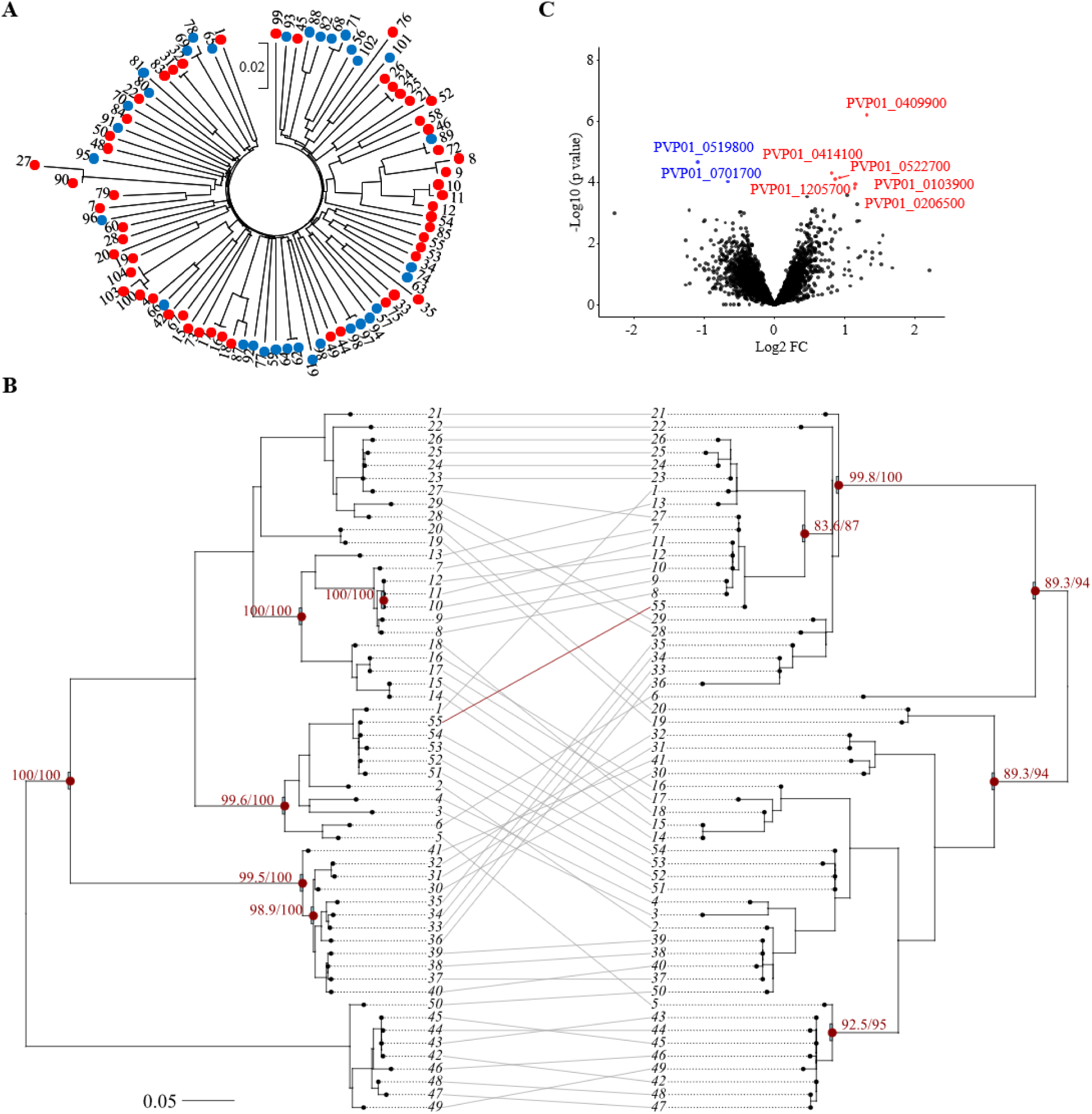
Genomic analyses of *P. vivax* and MaRNAV-1. **A**. Genetic relationships among monoclonal *P. vivax* isolates based on genome-wide SNPs (n=86). Red circles indicate MaRNAV-1-positive samples, while blue circles indicate MaRNAV-1-negative samples. The distribution of virus-positive samples across the tree indicates that MaRNAV-1 infection occurs across genetically diverse parasite backgrounds rather than being restricted to specific parasite lineages. **B.** Tanglegram comparing phylogenetic trees inferred for MaRNAV-1 genomic segments S1 (left) and S2 (right). Maximum likelihood (ML) trees were inferred independently for each segment. Lines connect corresponding viral sequences from the same sample across the two trees. Tip labels represent individual sample IDs. Node support values are displayed as SH-aLRT/UFboot, with supported nodes defined as SH-aLRT ≥ 80% and UFboot ≥ 70% (red dots). Branch lengths represent nucleotide substitutions per site, and both trees were midpoint rooted prior to visualization. A single discordant connection (in red) suggests a putative reassortment event between S1 and S2 segments. **C.** Differential gene expression analysis of *P. vivax* parasites comparing samples with high versus low MaRNAV-1 load (top and bottom quartiles, n = 22 each group). The volcano plot shows log_2_ fold change versus -log_10_ adjusted p-values. Blue dots indicate down-regulated genes in high virus load samples, while red dots indicate up-regulated genes.

Given the association of MaRNAV-1 with gametocytes, we then examined whether MaRNAV-1 was associated with enhanced transmission to mosquito vectors. We analyzed the results of membrane feeding assays using *Anopheles diru*s mosquitoes performed on 33 blood samples analyzed by RNA-seq (13 MaRNAV-1-negative and 20 MaRNAV-1-positive). Virus-positive samples resulted in 12 infectious feeds (i.e., at least one oocyst was found in successfully infected mosquitoes) out of 20 (60%), compared to 5 out of 13 (38%) for virus-negative samples. Both gametocytemia (χ_1_^2^= 10.5, p = 0.001) and MaRNAV-1 presence (χ_1_^2^= 4.9, p = 0.027) increased the likelihood of mosquito infection (Fig. 2M). Infected mosquitoes with virus-positive *P. vivax* exhibited a ∼ 7-fold increase in mean oocyst count compared to virus-negative isolates (mean ± SE oocyst number in mosquitoes with virus-positive *P. vivax*: 60.2 ± 3.54 vs. 8.62 ± 1.26, χ_1_^2^ = 11.8, p < 0.001, Fig. 2N). Gametocytemia was, however, not associated with oocyst intensity (χ_1_^2^ = 2.7, p = 0.096). When considering MaRNAV-1 as a continuous variable, higher virus load was also associated with increased oocyst prevalence in mosquitoes (χ_1_^2^= 5.9, p = 0.01, Fig. 2O), regardless of gametocyte density (virus load by gametocytemia interaction: χ_1_^2^= 0.01, p = 0.9). Gametocytemia was also associated with oocyst prevalence (χ_1_^2^= 10.7, p = 0.001). Consistently, higher virus load was associated with increased oocyst intensity (χ_1_^2^= 7.8, p = 0.005, Fig. 2P), while oocyst intensity was not significantly associated with gametocytemia (χ_1_^2^= 2.8, p = 0.09). However, there was a significant negative interaction between gametocytemia and virus load on oocyst intensity (χ_1_^2^= 5.4, p = 0.02, Fig. 2P). These results indicate that MaRNAV-1 load was associated with higher *P. vivax* transmission, in particular at low gametocytemia.

To validate these findings, we first determined whether a similar correlation between gametocytemia and virus load was observed in a larger dataset of samples. We quantified Pvs25 and Pvs47 transcripts, markers of female and male gametocytes, respectively, by RT-qPCR (relative to human RPS18) in 152 samples with MaRNAV-1 load quantified by RT-qPCR (expressed as S2 abundance relative to *P. vivax* 18S rRNA). Both Pvs25 and Pvs47 relative abundance were correlated with MaRNAV-1 loads (R = 0.32, p < 0.0001 and R = 0.32, p < 0.0001, n = 152, respectively, Supp. Fig. 4).

**Fig. 4.**
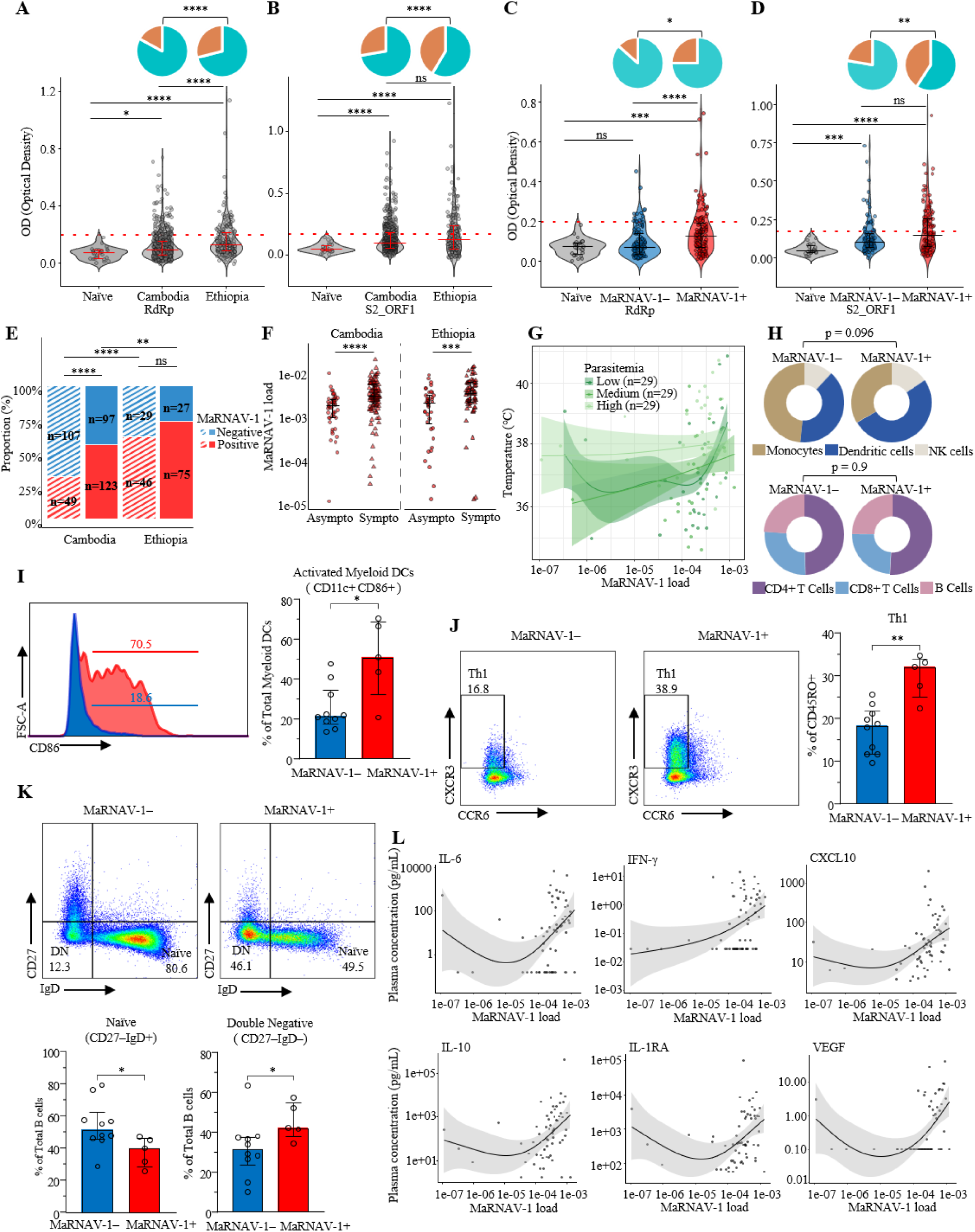
Association of MaRNAV-1 with host immune response and clinical outcomes in *P. vivax* infection. **A-B.** IgG responses measured by ELISA (optical density, OD) against MaRNAV-1 RdRp (**A**) and S2_ORF1 (**B**) antigens in malaria-naïve individuals (n = 23) and *P. vivax*-infected patients from Cambodia (n = 552) and Ethiopia (n = 176). Differences between groups were assessed using chi-square tests. **C-D.** IgG responses against MaRNAV-1 RdRp (**C**) and S2_ORF1 (**D**) stratified by MaRNAV-1 infection status (n = 112 virus-negative and n=120 virus-positive) in Cambodian *P. vivax*-infected and naïve (n = 23) individuals. Differences in seroprevalence between groups were assessed using Fisher’s exact tests. **(A-D)** The pie charts indicate the proportion of seropositive (orange) and seronegative (cyan) individuals in each group. In all panels, violin plots show the distribution of OD values with individual data points overlaid. The red dashed lines indicate the seropositivity thresholds defined as the mean + 3× standard deviations of malaria-naïve controls. **E**. Proportion of MaRNAV-1 positive and negative *P. vivax* infections in asymptomatic (dashed bar) and symptomatic (solid bar) individuals from Cambodia and Ethiopia. The prevalence of MaRNAV-1 among *P. vivax* infections was higher in Ethiopia than in Cambodia, in both asymptomatic and symptomatic groups (p < 0.0001). **F.** Comparison of MaRNAV-1 load, expressed as the ratio of viral S2 RNA to *P. vivax* 18S rRNA, between asymptomatic and symptomatic infections in Cambodia and Ethiopia. MaRNAV-1 load was significantly higher in symptomatic infections in both countries (p < 0.0001 and p = 0.0004, respectively). **G**. Body temperature as a function of MaRNAV-1 load and parasitemia (MaRNAV-1-positive samples only, n = 87). Generalized additive models were used to model nonlinear relationships. Curves represent temperature across MaRNAV-1 load, stratified by parasitemia levels (low: 107-4190 parasites/µL; medium: 4227-10119 parasites/µL; high: 10147-39353 parasites/µL). Shaded areas represent 95% CI. **H**. Relative proportions of innate (Monocytes, Dendritic cells, and NK cells) and adaptive immune cells (CD4+ T cells, CD8+ T cells, and B cells) in PBMCs according to MaRNAV-1 infection status in *P. vivax* patients (n = 5 MaRNAV-1-positive and n = 10 MaRNAV-1-negative). **I.** Proportion of activated myeloid dendritic cells out of total myeloid dendritic cells (DCs). There is a significant increase in the proportion of activated myeloid DCs in MaRNAV-1-positive samples (p = 0.0280). **J**. Proportion of Th1 cells (CD4+CCR4-CCR6-CXCR3+) out of CD4+CD45RO+ T cells. Th1 cells are significantly higher in MaRNAV-1-positive individuals (p = 0.0027). **K**. Distribution of B cell subsets (naïve and double negative B cells). MaRNAV-1-positive samples show reduced naïve B cells (CD19+IgD+CD27-) and increased double-negative B cells (CD19+IgD-CD27-) (p = 0.021, p = 0.04, respectively). **L**. Nonlinear associations between circulating cytokine levels (including IL-6, IFN-γ, CXCL10, IL-10, IL-1RA, and VEGF), as a function of MaRNAV-1 load (MaRNAV-1-positive samples only, n = 60). Curves represent GAM fits with shaded areas indicating 95% CI. Axes are shown on a log scale, and each point represents an individual sample. *: p < 0.05, **: p < 0.01, ***: p < 0.001, ***: p < 0.0001, and ns: non-significant

Then, we examined additional mosquito feeding experiments using blood samples from 87 patients for which MaRNAV-1 load was calculated by RT-qPCR (including the 33 patients’ samples already analyzed by RNA-seq). Of those, 48 samples were negative for MaRNAV-1 while 39 were positive. The results obtained confirmed that virus-positive samples had (i) higher infectiousness (virus-negative samples resulted in 13 infectious feeds out of 48 (27%), compared to 20 out of 39 (51%) for virus-positive samples), (ii) higher oocyst prevalence, and (iii) higher oocyst intensity, whether the gametocytemia covariate was calculated based on Pvs25 (Supp. Fig. 5), or Pvs47 transcripts (Supp. Fig. 6). Gametocytemia was also associated with both oocyst prevalence and intensity (Supp. Fig. 5 & 6).

**Fig. 5.**
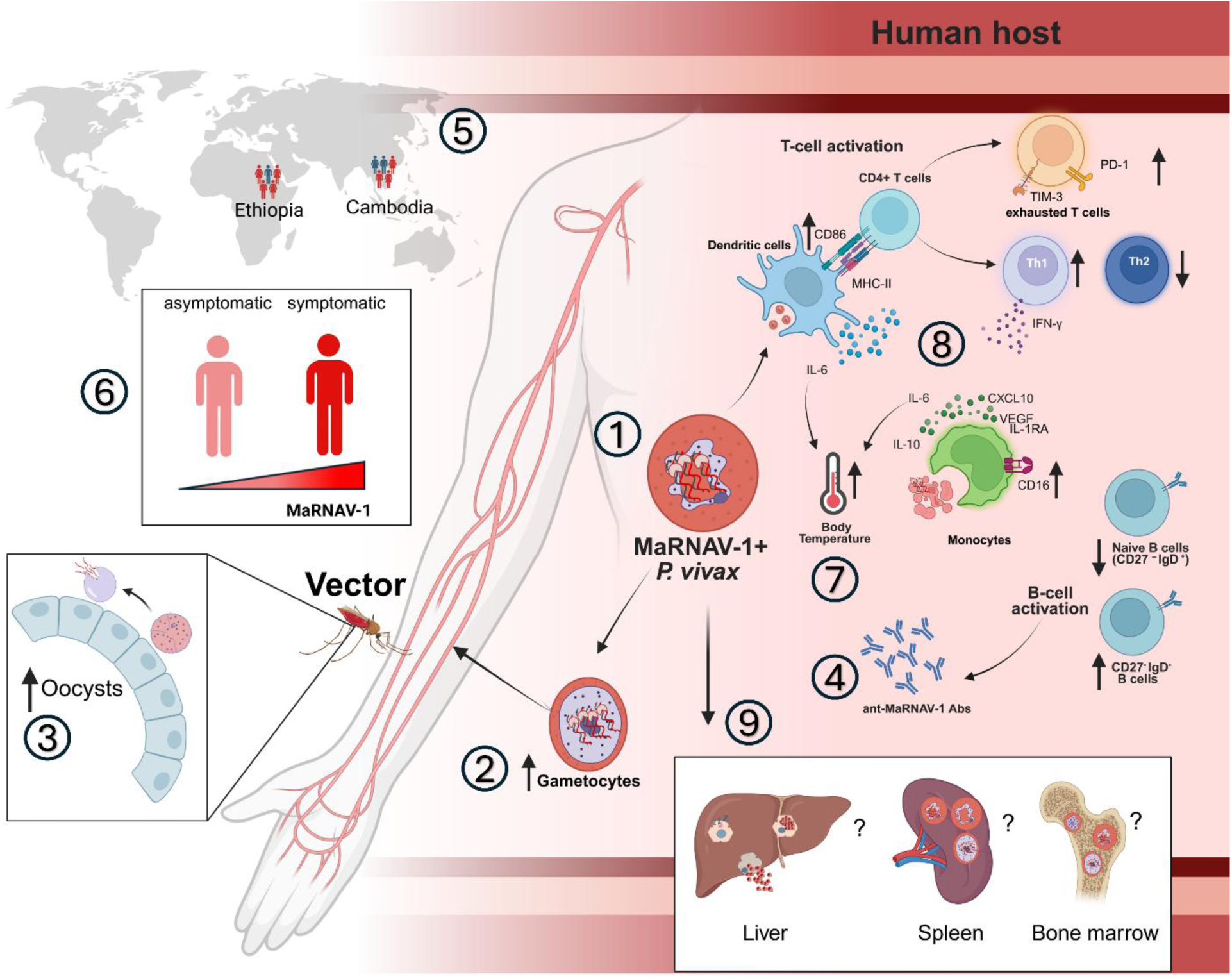
Proposed model of MaRNAV-1-associated modulation of *P. vivax* infection, transmission, and host responses. MaRNAV-1 infects *P. vivax* parasites and is detected across multiple parasite stages (1). MaRNAV-1-positive infections are associated with increased gametocytemia (2) and increased mosquito transmission, including higher oocyst prevalence and intensity (3), potentially by acting as a physiological stress signal that promotes gametocytogenesis. MaRNAV-1 elicits antibody responses in *P. vivax*-infected individuals (4), with geographic variation in seropositivity and prevalence between Cambodia and Ethiopia (5), which may relate to differences in transmission intensity. MaRNAV-1 load is higher in symptomatic than in asymptomatic *P. vivax* infections (6) and is associated with increased body temperature (7), innate immune cell activation, elevated cytokine production, and a shift toward a Th1-driven immune response during *P. vivax* malaria (8). Together, these observations support a model in which MaRNAV-1 reshapes *P. vivax* biology, host immunity, and parasite transmission. While this study focuses on peripheral-blood infections, important questions remain regarding the role of MaRNAV-1 during liver stages, including hypnozoite biology, as well as its potential presence and effects within cryptic parasite reservoirs in the spleen and bone marrow (9).

Overall, these analyses show a positive association between MaRNAV-1 and oocyst prevalence and intensity, independently of gametocytemia.

### MaRNAV-1-*P. vivax* association presents hallmarks of vertical transmission

To investigate whether MaRNAV-1 is vertically transmitted, we analyzed genomic and transcriptomic data from paired parasites and viral sequences. First, we sequenced the whole genome of *P. vivax* isolates that had been analyzed by RNA-seq and for which we had determined the MaRNAV-1 infection status (53 MaRNAV-1-positive and 33 MaRNAV-1-negative samples). To evaluate whether MaRNAV-1 infection was associated with specific parasite lineages, we reconstructed a neighbor-joining tree using whole-genome single-nucleotide polymorphism data from the *P. vivax* isolates. MaRNAV-1-positive samples were distributed across the parasite population and did not cluster within particular lineages, indicating that virus infection occurs across genetically diverse *P. vivax* backgrounds in this population, rather than being restricted to specific parasite genotypes (Fig. 3A).

Next, we examined the sequence diversity of the MaRNAV-1 S1 and S2 segments after *de novo* MaRNAV-1 transcript assembly from RNA-seq data. S1 exhibited substantial nucleotide variability (91.3%-100% identity) but was markedly more conserved at the amino acid level (96.5%-100% amino acid identity). S2 showed higher nucleotide conservation (94.1%-100% identity) and high amino acid conservation (ORF1: 96.6%-100%; ORF2: 96.3%-100%) (Supp. Data 3 to 7).

We then assessed within-patient virus diversity among the 63 samples for which we were able to reconstruct entire viral genomes. Among these, 27 isolates were polyclonal, containing multiple *P. vivax* genotypes. Interestingly, 12 of these 27 polyclonal *P. vivax* infections showed evidence of infection with multiple viral strains, whereas all monoclonal *P. vivax* infections (36/36, p < 0.0001) were associated with a single MaRNAV-1 strain, suggesting that within-patient viral diversity reflects parasite polyclonality.

Given the presence of multiple viruses in the same infection, we then tested whether the two segments were systematically co-inherited or whether reassortment could occur between MaRNAV-1 S1 and S2 segments. Using the subset of monoclonal *P. vivax* infection containing a single viral strain, we estimated a low reassortment rate of 0.004-0.005 events per year. Consistent with this, comparison of rooted phylogenetic trees for S1 and S2 using a tanglegram approach identified only a single putative reassortment event (highlighted in red, Fig. 3B). These results suggest limited segment exchange and vertical transmission of the virus, although this observation will require confirmation with larger, more geographically and temporally diverse genomic datasets.

Finally, we tested whether MaRNAV-1 infection affected *P. vivax* gene expression. After accounting for differences in parasite developmental stages, we compared the transcriptional profiles of the 38 samples with no detectable MaRNAV-1 reads to those of the 38 samples with the highest virus loads. No genes were differentially expressed between these groups (FDR > 0.1, Supp. Data 8). When restricting the analysis to MaRNAV-1 positive infections and considering samples with the most extreme virus loads (below the 25^th^ percentile versus above the 75^th^ percentile, n = 22 per group), only 8 genes were differentially expressed, indicating a very limited impact of MaRNAV-1 on the parasite transcriptome (Fig. 3C & Supp. Data 9). Among these, two genes of unknown function (PVP01_0519800; PVP01_0701700) were downregulated, while six were upregulated, including a PHIST-domain-containing exported protein (PVP01_0522700), a putative ELM2 domain-containing protein potentially involved in transcriptional regulation or chromatin remodeling (PVP01_0206500), a putative acyl-CoA synthetase (PVP01_0409900), a putative nucleoside diphosphate hydrolase (PVP01_1205700), and two additional genes of unknown function (PVP01_0103900; PVP01_0414100). Altogether, these results indicate that MaRNAV-1 has little, if any, impact on *P. vivax* gene expression profiles.

### MaRNAV-1 elicits antibody responses in *P. vivax-*infected individuals

Next, we examined the effect of MaRNAV-1 on the host immune response. We first assessed whether MaRNAV-1 elicits humoral immune responses in infected individuals. IgG responses against viral proteins (RdRP and S2_ORF1) were measured by ELISA in plasma samples from *P. vivax*-infected individuals in Cambodia and Ethiopia and compared with those of malaria-naïve controls. Detectable antibody responses against MaRNAV-1 RdRp were observed in 17% (94/552) and 29% (51/176) of *P. vivax-*infected individuals from Cambodia and Ethiopia, respectively, and against S2_ORF1 in 28% (153/552) and 41% (73/176) of infected individuals from Cambodia and Ethiopia, respectively (Fig. 4A & 4B). Seropositivity rates were higher in Ethiopian individuals compared to Cambodians for both RdRp (p = 0.0008) and S2_ORF1 (p = 0.0008) (Fig. 4A & 4B). Overall, seropositivity rates were significantly higher for the S2_ORF1 protein (31%, 226/728) than for RdRp (20%, 145/728, p < 0.0001). Malaria-naïve individuals showed homogenously low ELISA signal consistent with the lack of seroreactivity to MaRNAV-1.

We next analyzed samples for which both viral detection and ELISA data were available (n = 232), all from Cambodia. Seropositivity rates were significantly higher in individuals with MaRNAV-1 positive *P. vivax* infections (RdRp: 25%, 30/120, S2_ORF1: 41%, 49/120, Fig. 4C & 4D) compared with those with virus-negative infections (RdRp: 13%, 15/112, S2_ORF1: 22%, 25/112, p = 0.0307 and p = 0.0030, respectively, Fig. 4C & 4D). These results suggest that either some infections were misclassified as MaRNAV-1-negative and/or that some individuals were previously infected with a MaRNAV-1-positive infection. Altogether, these analyses show that MaRNAV-1 proteins, especially S2_ORF1, are immunogenic during *P. vivax* infection and elicit measurable antibody responses.

### MaRNAV-1 prevalence in *P. vivax* clinical isolates is associated with clinical outcomes and geographic origin

We next evaluated the prevalence of MaRNAV-1 among *P. vivax* clinical isolates of various origins. RT-qPCR screening of RNA extracted from blood samples of 220 symptomatic *P. vivax*-infected patients seeking treatment in Cambodia detected MaRNAV-1 in 56% of samples (123/220) (Fig. 4E). In contrast, among asymptomatic *P. vivax*-infected individuals identified through active case detection in malaria-endemic communities in Cambodia, MaRNAV-1 detection rate was significantly lower (31%, 49/156, p < 0.0001, Fig. 4E). There was no significant difference in *P. vivax* parasitemia (determined by microscopy examination of thick smears) between MaRNAV-1-positive and negative samples among symptomatic infections (median, IQR: 5685 parasites/µL, 3645–11305 and 5755 parasites/µL, 3232–12180, respectively; p = 0.8; Supp. Fig. 7A), while parasitemia (determined by RT-qPCR) was significantly lower in MaRNAV-1 negative asymptomatic infections compared to MaRNAV-1 positive ones (2.2 arbitrary units, AU, IQR: 0.77-4.62 and 3.97 AU, IQR: 1.53-7.16, respectively; p=0.02, Supp Fig. 7B). This suggests that some asymptomatic infections could have been misclassified as MaRNAV-1 negative due to low parasite density. To account for differences in parasitemia, we expressed MaRNAV-1 load as the ratio of viral S2 RNA to *P. vivax* 18S rRNA. Using this normalized measure, MaRNAV-1 load was significantly lower in asymptomatic compared to symptomatic infections (p < 0.0001; Fig. 4F).

To confirm and extend our findings in a distinct malaria-endemic setting, we evaluated the prevalence of MaRNAV-1 among *P. vivax* infections in Ethiopia. Among symptomatic patients, MaRNAV-1 was detected in 74% of samples (75/102), a prevalence significantly higher than that observed in symptomatic infections from Cambodia (p = 0.003, Fig. 4E). Similarly, prevalence among asymptomatic infections was significantly higher in Ethiopia (61%, 46/75) than in Cambodia (p < 0.0001, Fig. 4E). These results are consistent with the serological data, which showed higher MaRNAV-1 seropositivity in Ethiopia than in Cambodia (Fig. 4A & 4B). Although the same trend toward higher prevalence in symptomatic compared with asymptomatic infections was observed in Ethiopia, this difference did not reach statistical significance (p = 0.1, Fig. 4E). As in Cambodian samples, MaRNAV-1 load was significantly lower in asymptomatic infections compared to symptomatic ones (p = 0.0004, Fig. 4F).

### MaRNAV-1 is associated with increased body temperature and a shift towards a Th1-driven immune response

We next investigated whether MaRNAV-1 was associated with clinical parameters (hemoglobin, white blood cells and platelet counts, body temperature, and C-reactive protein levels) in *P. vivax*-infected patients, while adjusting for parasitemia. Clinical data were available for the 126 Cambodian patients enrolled in a clinical trial and for whom RNA-seq data were generated and presented above.

None of these clinical parameters were significantly associated with infection status (MaRNAV-1 negative/positive) (Supp. Table 2). However, among MaRNAV-1-positive individuals, body temperature was significantly associated with MaRNAV-1 load (FDR-adjusted p = 0.01), parasitemia (FDR-adjusted p = 0.04), and their interaction (FDR-adjusted p = 0.036) (Supp. Table 3). These findings indicate that, at low parasitemia, MaRNAV-1 load is associated with increased body temperature, whereas the effect of parasite burden on body temperature dominates at higher parasitemia (Fig. 4G, Supp. Fig. 8).

Next, to understand the impact of MaRNAV-1 infection on the host immune response, we performed gene expression analysis of whole blood from patients with low and high virus load (below the 25^th^ vs. above the 75^th^ percentile). Here, we identified only 16 genes differentially expressed (14 down-regulated and 2 up-regulated in the high MaRNAV-1 load group), including downregulation of genes involved in lipid metabolism and biosynthesis and upregulation of HLA-A (Supp. Fig. 9 & Supp. Data 10). Using gene expression deconvolution, differences in overall immune cell composition were associated with MaRNAV-1 infection status (p = 0.008), parasitemia (p < 0.001), and their interaction (p = 0.001) (Supp. Table 4 & Supp. Data 11).

Then, we performed in-depth immunophenotyping of peripheral blood mononuclear cells (PBMCs) isolated from 5 MaRNAV-1-positive and 10 MaRNAV-1-negative symptomatic *P. vivax*-infected patients (Supp. Fig. 10A, 10B & Supp. Table 5). Parasitemia was not different between MaRNAV-1-positive and negative infections (p > 0.999, Supp. Fig. 10C). No differences were observed in total cell composition, even though patients with a MaRNAV-1-positive infection tended to have increased dendritic cell numbers (Fig. 4H). We observed an increase in expression of CD86 on myeloid dendritic cells (p = 0.028, Fig. 4I) and a trend toward higher CD16 expression on monocytes (Supp. Fig. 10D), indicating activation and differentiation of the innate immune compartment in MaRNAV-1-positive *P. vivax*-infected patients. In addition, we observed a shift in Th balance with increased Th1 frequency (p = 0.0027, Fig. 4J) and a trend to decreased Th2 frequency (p = 0.0753, Supp. Fig. 10E) in MaRNAV-1-positive patients. Infection also tended to be associated with CD4+ T cell exhaustion, as suggested by increased frequency of TIM-3+PD-1+ CD4+ T cells (p = 0.0992, Supp. Fig. 10F), and alterations in the B cell compartment, with a decreased frequency of naïve B cells and increased frequencies of double negative IgD-CD27-B cells (p = 0.021 and p = 0.04, respectively, Fig. 4K).

Finally, circulating plasma cytokines were analyzed using a panel of canonical inflammatory cytokines. Comparing cytokine concentrations in samples with or without detectable MaRNAV-1 showed no significant difference for any of the cytokines (Supp. Table 6). Next, we evaluated the associations between MaRNAV-1 load and cytokine concentrations in MaRNAV-1-positive individuals. In accordance with a shift towards Th1-driven responses in MaRNAV-1-positive infected patients, IFN-γ was associated with MaRNAV-1 load (FDR-adjusted p = 0.047, Fig. 4L), but not with parasitemia (FDR-adjusted p = 0.103). Additionally, CXCL10, IL-10, IL-1RA, IL-6, and VEGF were significantly associated with both parasitemia and MaRNAV-1 load independently (Fig. 4L & Supp. Table 7). An interaction between virus load and parasitemia was detected (p = 0.011) only for IL-6, suggesting that virus load predominantly drives IL-6 secretion at low parasitemia, whereas parasite burden dominates at higher parasitemia (Supp. Table 7 & Supp. Fig. 11).

Together, these results indicate that MaRNAV-1 infection is associated with innate immune cell activation, increased cytokine production, and a shift towards a Th1-driven immune response during *P. vivax* malaria, thereby modulating clinical outcome during infection.

## Discussion and Conclusion

This study provides the first comprehensive characterization of MaRNAV-1, a recently described narnavirus associated with *P. vivax* infections^9^. Using complementary approaches, we demonstrate that MaRNAV-1 is localized within *P. vivax* parasites across multiple life stages. These observations provide direct evidence that MaRNAV-1 is an intracellular parasite-associated virus rather than a coincidental co-infection. Viral RNA clearance closely paralleled parasite clearance following antimalarial treatment, further indicating that MaRNAV-1 depends on parasite persistence within the human host.

Genomic and transcriptomic analyses suggest that MaRNAV-1 maintains a stable intracellular association with *P. vivax*. Although the viruses in our study populations differed by up to 8.7% in nucleotide sequence, viral proteins were still highly conserved, consistent with strong functional constraints acting on both the polymerase and S2-encoded proteins. We additionally observed limited evidence of reassortment between viral segments and found that MaRNAV-1-positive infections were distributed across genetically diverse *P. vivax* lineages rather than clustering within specific parasite genotypes. Despite strong phenotypic associations with transmission and host immunity, MaRNAV-1 had only minimal detectable effects on the parasite gene expression. Together, these observations support a long-term evolutionary association between MaRNAV-1 and *P. vivax*^10^, similar to patterns described for vertically transmitted endosymbiotic viruses infecting microbial eukaryotes^18,30^.

An intriguing feature of MaRNAV-1 is that S2 transcripts were more abundant than S1-derived polymerase reads, and the protein encoded by S2_ORF1 elicited a higher seroconversion rate in infected individuals than RdRp. These observations suggest that the S2 proteins (at least ORF1) may play important roles during viral replication or host-parasite interactions. Although narnaviruses are generally considered capsid-less viruses^26^, the high expression of S2 and immunogenicity of S2_ORF1 protein raise the possibility that they may contribute to viral particle stability, parasite cell interaction, or immune recognition. Further functional studies will be required to determine the precise role of these proteins in the MaRNAV-1 life cycle.

A major finding of this study is the association between viral presence and load and parasite transmission potential (Fig. 5). Virus-positive infections were associated with increased gametocyte proportions and enhanced mosquito infection success, including higher oocyst prevalence and intensity. These effects were observed independently of gametocytemia, suggesting that MaRNAV-1 may influence transmission biology beyond simply increasing gametocyte density.

The mechanisms underlying these transmission-associated phenotypes remain unclear. One possibility is that viral infection acts as a physiological stress signal within the parasite and promotes gametocytogenesis, similar to stress-induced sexual commitment observed in other *Plasmodium* systems^31–33^. Additionally, MaRNAV-1 may influence mosquito infection rates, indirectly through modulation of host immune or serum factors, or directly through altered parasite-mosquito interactions (i.e. modulation of mosquito immunity or alteration of midgut microbiome)^34,35^. MaRNAV-1-positive isolates may also be intrinsically more infectious. These findings suggest a potential convergence of selection forces between viruses and parasites: by enhancing mosquito infection, the virus may simultaneously favor both its own persistence and the transmission of *P. vivax*. Although the molecular mechanisms underlying this association are still unclear, these observations raise the possibility that parasite-associated viruses may contribute to variability in malaria transmission dynamics.

Our data further indicate that MaRNAV-1 is recognized by the human immune system during *P. vivax* infection. Antibody responses against viral proteins were detected in infected individuals from both Cambodia and Ethiopia. Interestingly, both viral prevalence measured by RT-qPCR and seropositivity rates were higher in Ethiopia than in Cambodia (Fig. 5). These geographic differences may reflect variation in malaria transmission intensity, although the factors underlying these differences have not yet been elucidated. The higher seropositivity observed among individuals infected with MaRNAV-1-positive parasites further supports that host antibody responses reflect exposure to virus-associated *P. vivax* infections.

In both Cambodia and Ethiopia, MaRNAV-1 loads were lower in asymptomatic than in symptomatic *P. vivax* infections (Fig. 5). This is consistent with our observation that higher virus loads are associated with elevated body temperature and increased concentrations of the pyrogenic cytokine IL-6. Interestingly, these relationships were strongest at low parasitemia levels, whereas parasite density became the dominant driver of body temperature and cytokine production at higher parasitemia. This pattern suggests that MaRNAV-1 may contribute to inflammatory signaling, particularly during early or low-density infections. It has been described for many years that the pyrogenic threshold is lower in *P. vivax* than in *P. falciparum* and that, at similar parasite densities, *P. vivax* induces a stronger inflammatory response in infected hosts compared to *P. falciparum*^8^. MaRNAV-1 may contribute to these differences in malaria physiopathology, potentially acting as an additional pathogen-associated molecular pattern activating the inflammatory response^36,37^. One possible mechanistic explanation is that viral RNAs or proteins may be released during schizont rupture or during phagocytosis of infected reticulocytes and sensed by host pattern recognition receptors, triggering dendritic cell maturation and monocyte activation, as observed in MaRNAV-1-positive patients. The concurrent induction of anti-inflammatory mediators, such as IL-10 and IL-1RA, suggests that the host immune response attempts to balance virus-induced inflammation with anti-inflammatory feedback to limit tissue damage. The association between virus load and VEGF production further raises the possibility that MaRNAV-1 may modulate vascular or inflammatory pathways implicated in malaria pathogenesis^38–40^.

Beyond innate inflammatory responses, MaRNAV-1 infection is associated with a shift from a Th2 to a more Th1-driven response, as shown by the increased frequency of CCR4-CCR6-CXCR3+ Th1 cells in MaRNAV-1-infected patients and the correlation between IFN-γ and CXCL10 concentrations and MaRNAV-1 load. Even though the impact of Th balance on clinical outcome after *P. vivax* infection remains to be fully understood, the increased IFN-γ production might further activate macrophages and contribute to the observed systemic inflammation and CD4+ T cell exhaustion. Viral presence is also associated with increased frequencies of CD27-IgD-B cells, a B cell subset suggested to be associated with chronic exposure and altered effector function in *P. falciparum* infection^41–43^. During *P. vivax* infection, this subset is expanded, and these cells are able to produce neutralizing antibodies^44–46^. Whether these alterations in the adaptive immune response are mainly driven by MaRNAV-1 or *P. vivax*-specific B and T cells has yet to be determined (Fig. 5). Although causality cannot be established from observational data, these results suggest that parasite-associated viruses may modulate the balance between asymptomatic carriage and clinical disease. Together, these findings suggest that MaRNAV-1 skews the host response towards a more anti-viral and pro-inflammatory response, possibly contributing to parasite immune evasion and altering clinical outcome.

Several important questions are currently unresolved. While this study focused primarily on peripheral blood infections, *P. vivax* also persists within cryptic tissue reservoirs, including the liver, spleen, and bone marrow^47^. We detected MaRNAV-1 in both liver schizonts and dormant hypnozoites, demonstrating that the virus is present during liver stages of the parasite life cycle. However, whether MaRNAV-1 influences liver stage biology, including hypnozoite dormancy, activation, or persistence, remains entirely unknown. Likewise, the presence and potential role of MaRNAV-1 within parasite populations in the spleen and bone marrow are still unexplored. Although our observational analyses identified robust associations between MaRNAV-1, transmission phenotypes, and host immune responses, causality cannot yet be established experimentally. The lack of continuous *P. vivax* culture systems and genetic manipulation tools is still a major limitation, although related experimentally tractable *Plasmodium* species may provide useful surrogate systems. In particular, the recent identification of a virus infecting *P. knowlesi*^11^ suggests that comparative experimental studies in related *Plasmodium* species could provide valuable insights into the biological consequences of parasite-associated virus infections.

Together, our findings identify MaRNAV-1 as a previously unrecognized component of *P. vivax* infection that may influence parasite transmission, host immunity, and clinical outcome. More broadly, these results reveal an unexpected layer of complexity in malaria biology and suggest that parasite-associated viruses may contribute to the pathogenicity of human malaria parasites.

## Methods

### Ethical statements

The use of samples and data for this study was approved by the National Ethics Committee of the Cambodian Ministry of Health (158-NECHR, 100-NECHR, 0364-NECHR, and 047-NECHR) and by the Ethiopian Public Health Institute Institutional Review Board (IRB/531/2023), Addis Ababa, Ethiopia. All participants or their guardians provided written informed consent prior to blood collection.

### Symptomatic patient enrolment, sample collection, and processing

*P. vivax*-infected blood samples used in this study were collected from treatment-seeking patients presenting with uncomplicated malaria and diagnosed with *P. vivax* infection by rapid diagnostic test or microscopy at local health facilities. *P. vivax* mono-infection was subsequently confirmed for all samples using a species-specific cytB PCR^48^. In Cambodia, patients were enrolled between 2018 and 2025 in Mondulkiri and Kampong Speu provinces, while in Ethiopia, patients were enrolled in the Arba Minch district between 2023 and 2025.

Following informed consent, venous blood samples were collected prior to treatment by healthcare providers in accordance with national treatment guidelines in each country. An aliquot of whole blood was immediately stored at −80 °C in TRIzol until RNA extraction. Blood was also used to prepare Giemsa-stained thick and thin smears for microscopy and was subsequently centrifuged to separate plasma from cell pellets, which were stored at −80°C until further use. For a subset of Cambodian patients, blood samples were additionally used for imaging analyses, experimental mosquito infections, and PBMC phenotyping (see below).

In addition, this study included samples and associated data from patients enrolled in a clinical trial in Cambodia evaluating the therapeutic efficacy of primaquine (NCT04706130), as described elsewhere^28^. At enrolment, complete blood counts, C-reactive protein levels, and body temperature were recorded. Venous blood collected prior to artesunate treatment was used for whole-genome sequencing, RNA-seq, and, in a subset of patients, mosquito infections. Clearance of both MaRNAV-1 and parasites following artesunate treatment was assessed in 20 of these patients by RT-qPCR of RNA extracted from capillary blood samples collected every 24 hours, from Day 0 (prior to treatment initiation) to Day 7.

### Asymptomatic participant enrolment, sample collection, and processing

*P. vivax*-infected blood samples from asymptomatic carriers used in this study were collected between 2023 and 2025 from participants enrolled in two longitudinal cohorts: one conducted in Mondulkiri, Cambodia, and the other in Arba Minch, Ethiopia. Blood samples were processed for TRIzol preservation for subsequent RNA extraction, as well as for DNA extraction and plasma separation. Genomic DNA was extracted using DNeasy kits (QIAGEN) and used for *P. vivax* detection by species-specific cytB PCR^48^.

### Experimental mosquito infections and liver stage assays

Membrane feeding assays (MFAs) were conducted using laboratory-reared *Anopheles dirus*, a primary vector in Southeast Asia. Blood samples collected from symptomatic treatment-seeking patients in Cambodia were used directly to feed 5 to 7-day-old female mosquitoes for 1h via an artificial membrane attached to a water-jacketed mini-feeders maintained at 37°C^49^. Unfed and partially fed females were discarded, while fed *A. dirus* mosquitoes were maintained at 26°C and 80% humidity and fed a 10% sucrose solution containing 0.05% PABA in distilled water. On the sixth day post-blood meal (dpbm), 50 mosquitoes were dissected, and the oocyst prevalence (the proportion of blood-fed mosquitoes with at least one oocyst in the midgut at 6 dpbm) and oocyst intensity (oocyst count per infected mosquito at 6 dpbm) were recorded. On days 15 to 18, individual mosquitoes were dissected to isolate sporozoites. These sporozoites were either fixed with 4% paraformaldehyde (PFA) for imaging or used immediately for liver stage assays. *P. vivax* liver stage assays were performed as previously described^50,51^. Primary human hepatocytes (PHH, purchased from BioIVT) were seeded in a 384-well plate (18,000 cells per well), and 15,000 sporozoites were added 48h later to each well, allowing them to infect the PHH. Culture medium was changed 24h after infection and then every second day until day 15 post-infection, when cells were fixed with 4% PFA.

### SmiFISH detection of S1 and S2 MaRNAV-1 RNA

The primary probes and FLAPs (secondary probes with fluorescent tags) were synthesized and purchased from Integrated DNA Technologies (IDT). The primary probes were designed using the Stellaris FISH Probe Designer (biosearchtech.com/stellarisdesigner). Twenty-one primary probes were synthesized for MaRNAV-1 S1 and S2, and 27 were synthesized for *P. vivax* 18S rRNA. All probe sequences are available in Supp Data. 12. The secondary probes were conjugated to Cy3, Cy5, and Alexa Fluor 488. The FLAP-Y sequences were used as secondary probes as previously described^52^. 18S rRNA was labeled with FLAP-Y-Alexa Fluor 488, S1 with FLAP-Y-Cy5, and S2 with FLAP-Y-Cy3. *P. vivax*-infected red blood cells were enriched using a KCl-Percoll density gradient^53^, fixed at room temperature in 4% PFA and 0.0075% glutaraldehyde. The cells were washed twice with 1X PBS and hybridized according to Stellaris’ instructions for suspension cells. Image stacks were captured using a confocal microscope (Leica SP8) with a 63X/1.4 NA oil-immersion objective and a sCMOS PCO camera. Subsequent image analysis was performed using ImageJ software (1.8.0).

### Immunofluorescence detection of MaRNAV-1 RdRp

Recombinant RdRp and rabbit polyclonal antibodies were purchased from Genscript. Briefly, the full-length RdRp (NCBI sequence) was expressed in pET30a in *E. coli*, purified from inclusion bodies, and used to immunize New Zealand rabbits. Antibodies were purified by antigen affinity purification, titrated by ELISA, and shipped to Cambodia for immunofluorescence assays (IFA). Fixed *P. vivax*-infected red blood cells or sporozoites were placed on poly-L-lysine-coated slides. Cells were permeabilized using 0.3% Triton X-100, then washed with PBS, and incubated overnight with polyclonal anti-RdRp (1:1000). After washing, samples were incubated with goat anti-rabbit Alexa Fluor 488 (1:1000, Thermo Fisher). Samples were washed, and DNA was stained with Hoechst 33342 (1:1000 dilution). Slides were mounted with Vectashield^®^ (Vector Laboratories Cat #H-1000) prior to imaging. IFA on infected hepatocytes followed a similar procedure with some adaptations. Fixed cells were stained overnight with recombinant mouse-anti *P. vivax* Upregulated in Infectious Sporozoites 4 (rPvUIS4, 1:10,000)^54^ and rabbit anti-RdRp antibody (1:1000). After washing, samples were then stained with goat anti-mouse Alexa Fluor™ 488 and anti-rabbit Alexa Fluor™ Plus 594 IgG (H+L) secondary antibodies (1:1000, Thermo Fisher). After washing, the cell DNA was stained with Hoechst 33342. Stained red blood cells, sporozoites, and hepatocytes were visualized using a Lionheart FX Automated Microscope (Biotek^®^). Subsequent image analysis was performed using ImageJ (1.8.0).

### RNA extraction and RT-qPCR detection and quantification of MaRNAV-1, gametocytemia, and parasitemia

Total RNA was extracted from TRIzol-preserved samples using chloroform. After centrifugation at 12,000 g for 15 minutes, the aqueous phase was collected and mixed with 100 % ethanol. RNA purification was performed at 4 °C using the RNeasy Mini Kit (QIAGEN), according to the manufacturer’s instructions. Residual genomic DNA was digested with DNase I (Promega). Reverse RNA transcription to complementary DNA (cDNA) was carried out using Promega GoScript Reverse Transcriptase Kit. All qPCRs were conducted in a 20 µL volume with 0.25µM of primers, 1X EvaGreen SYBR mix, and 1 µL of cDNA. All reactions were performed under the following conditions: 95 °C for 15 minutes, followed by 45 cycles of 95 °C for 15 seconds, 60°C for 20 seconds for human RPS18 and S2 or 61°C for 20 seconds for Pvs47 or 64°C for 30 seconds for *P. vivax* 18S rRNA and Pvs25, and 72 °C for 20 seconds, with a subsequent melting curve analysis. All primers used are listed in Supp. Table 8. MaRNAV-1 load was determined by relative quantification of S1 or S2 normalized to *P. vivax* 18S rRNA. As S2 transcripts were consistently more abundant than S1, S2 relative quantification was preferred for increased sensitivity. Gametocytemia was determined by relative quantification of either Pvs25 or Pvs47 normalized to human RPS18. Parasitemia in asymptomatic infections was determined by relative quantification of *P. vivax* 18S rRNA, normalized to human RPS18. For symptomatic infections, parasitemia was determined by microscopy, counting all parasites on Giemsa-stained thick blood films, and normalized to the number of white blood cells (WBCs) counted, assuming 8000 WBCs/µL.

### RNA extraction, library preparation, and RNA sequencing analysis

RNA was extracted from TRIzol-preserved samples using phenol-chloroform. After rRNA depletion and polyA selection (NEB), RNA-seq libraries were prepared using the NEBNext Ultra II Directional RNA Library Prep Kit (NEB). We sequenced all libraries on an Illumina NovaSeq 6000 to generate ∼30-414 million paired-end reads of 75 bp per sample. We first aligned all reads from each sample using Hisat2 (v2.1.0)^55^ to a FASTA file containing the *P. vivax* P01 and human hg38 genomes, using default parameters and a shorter maximal intron length (--max-intronlen 5000)^56^. We retained only unmapped reads and then de novo assembled them into contigs using Trinity^57^. Minimap was used to map the contigs to the reference sequences of MaRNAV-1 (S1: accession No. MN860568.1, S2: accession No. MN860569.1). The number of different de novo assembled MaRNAV-1 sequences was used to determine whether a single or multiple viral strains were present in one infection. Finally, we realigned all reads to a file containing all NCBI- and de novo-assembled virus sequences using Hisat2, and calculated the number of MaRNAV-1 reads. Separately, we calculated read counts per gene using gene annotations downloaded from PlasmoDB (*P. vivax* genes) and NCBI (human genes), and then used *subread featureCounts* (v1.6.4)^58^. Using these data, MaRNAV-1 load was calculated for these RNA-seq samples as the ratio of reads mapping to MaRNAV-1 to those mapping to *P. vivax*.

Read counts per gene were normalized to transcripts per million (TPM) for *P. vivax* genes and the virus, and to counts per million (cpm) for humans and *P. vivax* genes separately for differential gene analysis. To filter out lowly expressed genes, only genes expressed at least 10 cpm were retained for further analysis (9493 human and 4934 *P. vivax* genes, respectively). edgeR^59^ was used for statistical assessment of differential gene expression analysis with a quasi-likelihood negative binomial generalized model. All results were corrected for multiple testing using FDR^60^, and FDR < 0.1 were considered significant.

We also estimated the proportion at each developmental stage and the proportion of human immune cell types in individual blood samples from infected individuals. We performed gene expression deconvolution using CIBERSORTx^61^, and a custom signature matrix derived from orthologous *Plasmodium berghei* genes was used for *P. vivax* stage deconvolution. We used the total proportion of male and female gametocytes obtained from this deconvolution, multiplied by total parasitemia (obtained from thick smear microscopy), to quantify absolute gametocytemia in these RNA-seq samples. To deconvolute human gene expression profiles, we used a validated leukocyte gene signature matrix, which uses 547 genes to differentiate 22 immune subtypes^62^.

### DNA extraction, whole genome sequencing, and complexity of infection

We extracted parasite DNA from leukocyte-depleted blood samples using the DNeasy blood and tissue kit (Qiagen) and prepared whole genome sequencing libraries using the NEBNext ^R^ Ultra^TM^ II FS DNA Library Prep Kit for Illumina NovaSeq 6000 to generate 25-50 million paired-end reads of 100 bp per sample. We used Hisat2^55^ with default parameters to map the reads to the P01 reference genome^63^ (version 67). Samples with an average coverage greater than 50X were further analyzed. We then estimated whether each sample was monoclonal or polyclonal, using GATK^64^ to call nucleotide variants, excluding telomeric regions and multigene families. We considered only positions with at least 20X coverage in at least 80% of the samples and only polymorphic positions with a maximum of 2 alleles. We then used moimix^65^ to estimate polyclonality, with a F_ws_ ≥ 0.95 being considered as monoclonal and those F_ws_ < 0.95 being considered as polyclonal. We then analyzed the relatedness of monoclonal samples by calculating pairwise distances. We used the proportion of those positions without shared nucleotides to generate a distance matrix. We used this matrix to generate a Neighbor-Joining tree in MEGA11^66^.

### Genetic analysis of MaRNAV-1 and sequence rearrangement determination

Multiple nucleotide sequence alignments for the MaRNAV-1 S1 and S2 were generated using the MUSCLE algorithm in MEGA11^66^. The phylogenetic trees of the S1 and S2 of the MaRNAV-1 were inferred using a maximum likelihood approach. To improve alignment quality, we filtered out poorly aligned regions and gaps using trimAl with an automated method^67^. All trees were estimated using IQ-TREE2 (v3.0.1). To assess the robustness of the resulting topologies, we calculated branch support using 1,000 bootstrap replicates with the UFBoot2 algorithm and an implementation of the SH-like approximate likelihood ratio test within IQ-TREE2^68,69^. The best-fit model of nucleotide substitution was evaluated using the Akaike information criterion (AIC), the corrected AIC, and the Bayesian information criterion (BIC) implemented in the ModelFinder function in IQ-TREE2^70,71^, while TPM2u+I+G4 was selected according to BIC for downstream phylogenetic analyses. The tanglegram was annotated using the R package phytools (v2.5.2)^72^.

### Detection of antibodies against MaRNAV-1 proteins by ELISA

We determined if antibodies against RdRP and S2_ORF1 could be detected in human plasma samples by ELISA. Recombinant RdRp (see above) and recombinant S2_ORF1 were purchased from Genscript (expressed in *E. coli*). Note that attempts to express and purify S2_ORF2 in high enough yield and purity failed. Maxisorp plates (Nunc Immunoplate, Thermo Scientific) were coated overnight with RdRp protein at 2 μg/ml and with S2_ORF1 at 1 μg/ml. Wells were blocked at 37 °C for 1 hour with blocking buffer (5% non-fat milk in PBS-0.05% Tween-20). Human plasmas were added at a 1:50 dilution to the wells for 1h at 37°C. After washing, goat anti-human IgG-HRP was added at a dilution of 1:10,000 for RdRp protein and 1:50,000 for S2_ORF1 for 1 hour at 37 °C. The assay was developed using a TMB substrate solution (BioLegend) and stopped with 0.18 M H_2_SO_4_. The absorbance at 450 nm was measured using the iMark Microplate Absorbance Reader. Human plasmas used were collected from confirmed *P. vivax*-infected individuals from both Cambodia and Ethiopia. Negative controls were obtained from plasma collected from malaria-naïve European volunteers residing in Phnom Penh.

### Flow Cytometry Phenotyping of Innate and Adaptive Immune Cells

Phenotyping of innate and adaptive immune cells was performed by flow cytometric extracellular staining in *P. vivax*-infected symptomatic individuals seeking treatment, stratified into virus-positive (n = 5) and virus-negative (n = 10) groups. Cryopreserved PBMCs were thawed, rested, and assessed for viability before staining. Two antibody panel designs were employed to characterize innate and adaptive immune cell populations (Supp. Table 5). Cells for both panels underwent the same staining procedure with a minimum of 200,000 cells per well. Cells were first incubated with an antibody cocktail targeting chemokine markers for 30 min at 37 °C in 5% CO₂, followed by washing with 1× PBS (Fisher Scientific). Fc receptors were subsequently blocked using Human TruStain FcX Blocking Reagent (BioLegend) for 10 min at 4 °C. Without an intermediate wash step, cells were stained with surface marker antibody cocktails specific to each panel for 30 min at 4 °C. After washing with 1× PBS, cells were incubated with Zombie Aqua Fixable Viability Dye (BioLegend) for 20 min at 4 °C in the dark. Finally, cells were washed and resuspended in FACS buffer (PBS supplemented with BSA and EDTA) prior to acquisition on a FACSAria Fusion flow cytometer (BD Biosciences). Data were analyzed using FlowJo software (version 10.10.0).

### Multiplex assay determination of plasma cytokines

Cytokine concentrations in human plasma samples were determined using a magnetic bead-based multiplex Luminex multi-analyte profiling (MAP) assay (MAGPIX™ systems) with the Cytokine 30-Plex Human Panel (Thermo Fisher Scientific), following the manufacturer’s instructions. Results were plotted as picograms per milliliter. Values below the lower limit of detection (LLOD) were replaced with the lowest detectable measure divided by 2 for data analysis.

## Statistical Analyses

Descriptive statistics and nonparametric tests were performed using GraphPad Prism (version 10). We used R (version 4.5.1) and RStudio (version 2025.05.1) for Spearman correlations, all regressions, and modeling analyses. Mann-Whitney U tests were used for two-group comparisons and Kruskal-Wallis tests for comparisons involving three or more groups, for nonparametric comparisons. Fisher’s exact test was used for pairwise comparisons of proportions, and the chi-square test was used for comparisons involving larger samples. *Mosquito transmission analyses*. To examine the impact of the MaRNAV-1 on transmission potential, we analyzed oocyst counts from the mosquito membrane feeding assay. We used negative binomial hurdle models fitted with the glmmTMB package^73^. This approach models separately the infection prevalence (binomial component) and the infection intensity among infected mosquitoes (zero-truncated negative binomial component). Both components included log-transformed parasitemia, log-transformed virus load or virus infection status, and their interaction as fixed effects, and patient identity as a random intercept. The significance of fixed effects was assessed using likelihood ratio tests comparing nested models. Model assumptions were evaluated using simulated residual diagnostics (DHARMa package). *Clinical and cytokine analyses*. To investigate the associations between MaRNAV-1 and clinical correlates (hemoglobin, white blood cell and platelet counts, body temperature, and C-reactive protein levels) or host immune response (cytokines, chemokines, and growth factors), two complementary modeling approaches were used. When MaRNAV-1 infection status was treated as a binary variable (positive/negative), linear regression models were fitted with virus infection status, log-transformed parasitemia, and their interaction, if necessary, as explanatory variables. When MaRNAV-1 load was analyzed as a continuous variable, generalized additive models (GAM) were fitted using default settings (generalized cross-validation with n = 87 for clinical correlates, n = 60 for cytokines, and thin plate regression splines) in the mgcv package, with smooth terms for log-transformed virus load and log-transformed parasitemia, and tensor product interaction terms to model nonlinear interactions between these two variables when necessary. An additional set of analyses was conducted to evaluate dose-response relationships among MaRNAV-1-positive individuals. *Immune cell composition analysis*. To determine whether the proportions of immune cell types were affected by MaRNAV-1 and parasitemia, a Dirichlet regression^74^ model was applied to jointly analyze the composition of 19 cell types. We fitted a model with MaRNAV-1 infection status, log-transformed parasitemia, and their interaction as predictors. The significance of each term was assessed by likelihood ratio tests comparing the full model to a reduced model with the term removed. These p-values correspond to global tests across all 19 cell-type components jointly; cell-type–specific associations were interpreted from individual regression coefficients. To account for multiple testing^60^, p-values were adjusted using the Benjamini-Hochberg method, applied separately for each predictor across all outcome models. An FDR-adjusted p < 0.05 was considered statistically significant.

## Acknowledgments

We are grateful to all the patients and healthcare workers involved in this study, as well as to the staff at the Malaria Research Unit at the Pasteur Institute in Cambodia. We also thank Julien Fernandes from the Photonic Bioimaging Platform at the Institut Pasteur, Paris, France, for his assistance with microscopy. Part of this study was supported by NIH awards R01AI146590 and R01AI153083 (to D. Ser) and R01AI175134 and R01AI173171 (to J.P.). J.P. is further supported by the NIH/NIAID (R61AI187100 and ICEMR Asia-Pacific U19AI129392) and the Pasteur International Unit PvESMEE. Work in the laboratory of S.B. is supported by an ERC Starting Grant (# 947819) and baseline funding of the Institut Pasteur. J.G. is supported by a DFG Walter Benjamin Postdoc Fellowship.D. Sen is supported by a scholarship from the Institut Pasteur du Cambodge and by a Calmette and Yersin internship grant from the Institut Pasteur.

## Author contributions

Conceptualization: J.P., D. Ser.

Methodology and Investigation: D. Sen., K.K., A.O., S.H., L.B.F.D., N.Y., J.G., F.D., K.T., T.V., G.D., N.K., J.S., Z.O., T.L., A.C., C.S., C.F., A.A.R., A.A., M.T., G.Ta., G.To., B.M., T.T.S., E.L., S.B., D.Ser., J.P.

Validation: D. Sen., A.O., T.L., B.M., T.C., N.M.A., M.E.P, S.B., D. Ser., J.P.

Formal analysis: D.Sen., T.L., D.Ser., J.P.

Writing (original draft): D.Sen., J.P

Writing (review and editing): all authors

Supervision: A.O., E.L., A.A, T.C., S.B., D.Ser., J.P.

## List of Supplementary figures, tables, and data

### Supp. Figures

Supp. Fig 1. Genomic structure of MaRNAV-1

Supp. Fig. 2. smiFISH and immunofluorescence assay (IFA) to detect MaRNAV-1 RNA and RdRp

Supp. Fig. 3. Associations between MaRNAV-1 load and blood stage parasite composition and gametocytemia by RNA-seq

Supp. Fig. 4. Spearman correlation between MaRNAV-1 load and gametocytemia by RT-qPCR

Supp. Fig. 5. Association between MaRNAV-1 infection status or virus load and transmission of *P. vivax* to mosquito vectors

Supp. Fig. 6. Association between MaRNAV-1 infection status or viral load and transmission of *P. vivax* to mosquito vectors using Pvs47 as gametocytemia measure

Supp. Fig. 7. Comparison of parasitemia levels between MaRNAV-1 negative and positive samples in symptomatic and asymptomatic Cambodian *P. vivax* infected patients

Supp. Fig. 8. Interaction between MaRNAV-1 load and parasitemia on predicted body temperature

Supp. Fig. 9. Host gene expression analysis between high and low MaRNAV-1 load in *P. vivax*-infected patients

Supp. Fig. 10. Flow cytometry gating strategy and immune cell subset proportions in *P. vivax*-infected patients stratified by MaRNAV-1 infection status

Supp. Fig. 11. Interaction between MaRNAV-1 load and parasitemia on IL-6 levels.

## Supp. Tables

Supp. Table 1. Clearance kinetics of *P. vivax* parasites and MaRNAV-1 following artesunate treatment.

Supp. Table 2. Association between MaRNAV-1 infection status and clinical correlates in *P. vivax*-infected patients.

Supp. Table 3. Non-linear associations between MaRNAV-1 load and clinical correlates in MaRNAV-1-positive *P. vivax*-infected patients

Supp. Table 4. Global effects of MaRNAV-1 infection, parasitemia, and their interaction on immune cell composition

Supp. Table 5. Monoclonal antibody panel for flow cytometry (innate and adaptive immune panels)

Supp. Table 6. Association between MaRNAV-1 infection status and plasma cytokine levels in *P. vivax*-infected patients

Supp. Table 7. Non-linear associations between MaRNAV-1 load and cytokine levels in MaRNAV-1-positive *P. vivax*-infected patients.

Supp. Table 8. Primers used in this study

## Supp. Data

Supp. Data 1. RNA-seq reads mapping to MaRNAV-1 and to *P. vivax*.

Supp. Data 2. Summary data of *P. vivax* genes expressed as TPM (transcripts per million) for calculating the threshold of the 25^th^ and 75^th^ percentiles of *P. vivax* genes

Supp. Data 3. MaRNAV-1 S1 nucleotide alignment

Supp. Data 4. MaRNAV-1 S2 nucleotide alignment

Supp. Data 5. RdRP protein alignment

Supp. Data 6. S2_ORF1 protein alignment

Supp. Data 7. S2_ORF2 protein alignment

Supp. Data 8. *P. vivax* DGE analysis of presence and absence of MaRNAV-1.

Supp. Data 9. *P. vivax* DGE analysis of low and high of MaRNAV-1 load (below the 25^th^ percentile versus above the 75^th^ percentile)

Supp. Data 10. Results of human DGE analysis comparing samples with MaRNAV-1 load below the 25^th^ percentile to those above the 75^th^ percentile

Supp. Data 11. Significant component-level associations identified by Dirichlet regression analysis of immune cell composition

Supp. Data 12. Primary probes sequences used in smiFISH, including *P. vivax* 18 rRNA, S1, and S2 segments of the MaRNAV-1.

