## Supplementary figures and images for "MaRNAV-1, an intracellular virus of *Plasmodium vivax*, is associated with increased parasite transmission and altered host immune response"

### Supp data 3

NCBI Multiple Sequence Alignment Viewer, Version 1.26.0

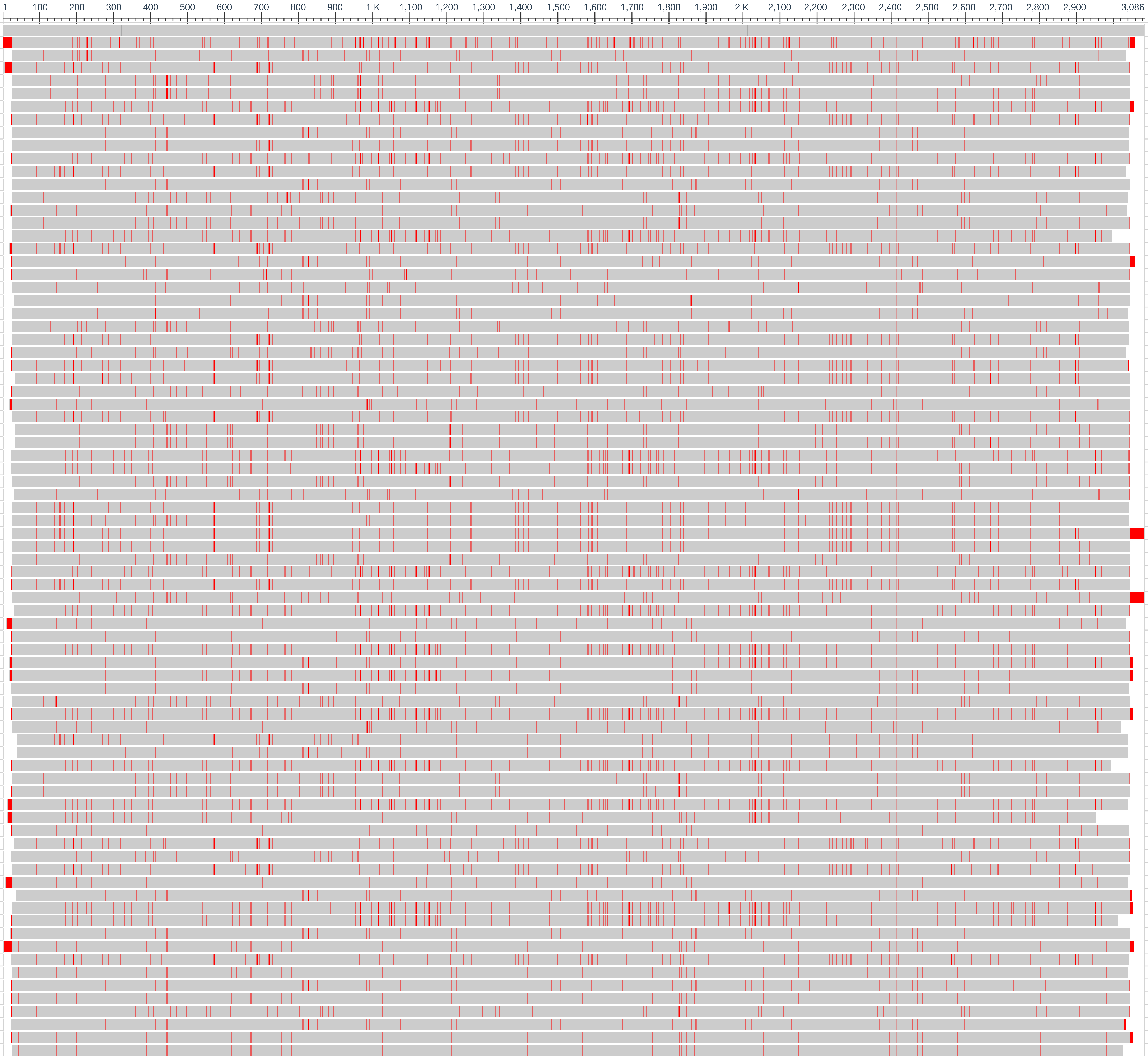


| Sequence ID | | Start | End |
|---|---|---|---|
| consensus | (+) | 1 | 3,086 |
| MN860568.1 | (+) | 1 | 3,057 |
| 29 | (+) | 1 | 3,006 |
| 30 | (+) | 1 | 3,044 |
| 58 | (+) | 1 | 3,018 |
| 58-1 | (+) | 1 | 3,018 |
| 42 | (+) | 1 | 3,032 |
| 33 | (+) | 1 | 3,027 |
| 105 | (+) | 1 | 3,017 |
| 105-1 | (+) | 1 | 3,017 |
| 49 | (+) | 1 | 3,025 |
| 37 | (+) | 1 | 3,008 |
| 23 | (+) | 1 | 3,020 |
| 51 | (+) | 1 | 3,014 |
| 7 | (+) | 1 | 3,017 |
| 52 | (+) | 1 | 3,021 |
| 43 | (+) | 1 | 2,970 |
| 36 | (+) | 1 | 3,029 |
| 22 | (+) | 1 | 3,036 |
| 13 | (+) | 1 | 3,025 |
| 19 | (+) | 1 | 3,018 |
| 21 | (+) | 1 | 3,012 |
| 28 | (+) | 1 | 3,016 |
| 67 | (+) | 1 | 3,018 |
| 67-1 | (+) | 1 | 3,018 |
| 5 | (+) | 1 | 3,012 |
| 34 | (+) | 1 | 3,021 |
| 40 | (+) | 1 | 3,011 |
| 2 | (+) | 1 | 3,024 |
| 14 | (+) | 1 | 3,027 |
| 41 | (+) | 1 | 3,022 |
| 110 | (+) | 1 | 3,012 |
| 110-1 | (+) | 1 | 3,012 |
| 111 | (+) | 1 | 3,025 |
| 111-1 | (+) | 1 | 3,026 |
| 3 | (+) | 1 | 3,024 |
| 20 | (+) | 1 | 3,015 |
| 112 | (+) | 1 | 3,015 |
| 112-1 | (+) | 1 | 3,015 |
| 38 | (+) | 1 | 3,060 |
| 113 | (+) | 1 | 3,018 |
| 113-1 | (+) | 1 | 3,018 |
| 50 | (+) | 1 | 3,025 |
| 39 | (+) | 1 | 3,023 |
| 4 | (+) | 1 | 3,059 |
| 47 | (+) | 1 | 3,015 |
| 18 | (+) | 1 | 3,024 |
| 116 | (+) | 1 | 3,026 |
| 116-1 | (+) | 1 | 3,026 |
| 117 | (+) | 1 | 3,035 |
| 117-1 | (+) | 1 | 3,035 |
| 79 | (+) | 1 | 3,019 |
| 53 | (+) | 1 | 3,018 |
| 45 | (+) | 1 | 3,031 |
| 15 | (+) | 1 | 2,991 |
| 118 | (+) | 1 | 3,001 |
| 118-1 | (+) | 1 | 3,001 |
| 48 | (+) | 1 | 2,971 |
| 54 | (+) | 1 | 3,023 |
| 55 | (+) | 1 | 3,024 |
| 120 | (+) | 1 | 3,028 |
| 120-1 | (+) | 1 | 2,940 |
| 16 | (+) | 1 | 3,020 |
| 35 | (+) | 1 | 3,015 |
| 6 | (+) | 1 | 3,026 |
| 31 | (+) | 1 | 3,021 |
| 17 | (+) | 1 | 3,032 |
| 27 | (+) | 1 | 3,014 |
| 46 | (+) | 1 | 3,029 |
| 44 | (+) | 1 | 2,990 |
| 24 | (+) | 1 | 3,024 |
| 8 | (+) | 1 | 3,052 |
| 32 | (+) | 1 | 3,019 |
| 9 | (+) | 1 | 3,016 |
| 26 | (+) | 1 | 3,026 |
| 10 | (+) | 1 | 3,025 |
| 1 | (+) | 1 | 3,026 |
| 25 | (+) | 1 | 3,012 |
| 12 | (+) | 1 | 3,031 |
| 11 | (+) | 1 | 3,001 |

### Supp data 4

NCBI Multiple Sequence Alignment Viewer, Version 1.26.0

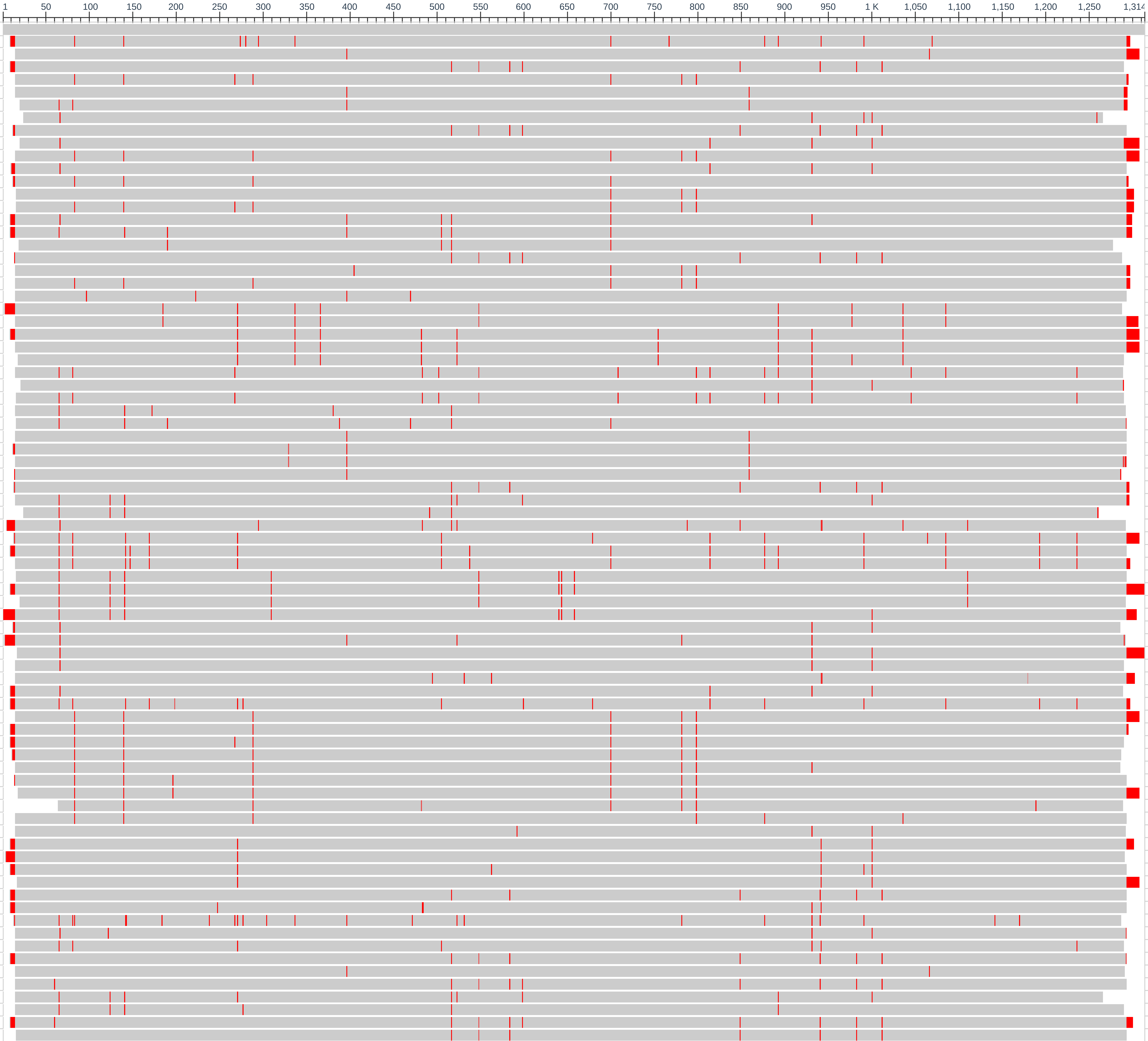


| Sequence ID | | Start | End |
|---|---|---|---|
| consensus | (+) | 1 | 1,314 |
| MN860569.1 | (+) | 1 | 1,289 |
| 1 | (+) | 1 | 1,293 |
| 10 | (+) | 1 | 1,281 |
| 100 | (+) | 1 | 1,281 |
| 105 | (+) | 1 | 1,280 |
| 105-1 | (+) | 1 | 1,275 |
| 108 | (+) | 1 | 1,241 |
| 11 | (+) | 1 | 1,281 |
| 110 | (+) | 1 | 1,288 |
| 111 | (+) | 1 | 1,293 |
| 113 | (+) | 1 | 1,283 |
| 116 | (+) | 1 | 1,284 |
| 117 | (+) | 1 | 1,286 |
| 117-1 | (+) | 1 | 1,286 |
| 118 | (+) | 1 | 1,291 |
| 118-1 | (+) | 1 | 1,291 |
| 119 | (+) | 1 | 1,259 |
| 12 | (+) | 1 | 1,274 |
| 120 | (+) | 1 | 1,283 |
| 120-1 | (+) | 1 | 1,283 |
| 13 | (+) | 1 | 1,278 |
| 14 | (+) | 1 | 1,285 |
| 15 | (+) | 1 | 1,292 |
| 16 | (+) | 1 | 1,299 |
| 17 | (+) | 1 | 1,293 |
| 18 | (+) | 1 | 1,272 |
| 19 | (+) | 1 | 1,274 |
| 2 | (+) | 1 | 1,270 |
| 20 | (+) | 1 | 1,274 |
| 21 | (+) | 1 | 1,277 |
| 22 | (+) | 1 | 1,279 |
| 23 | (+) | 1 | 1,278 |
| 24 | (+) | 1 | 1,282 |
| 25 | (+) | 1 | 1,280 |
| 26 | (+) | 1 | 1,274 |
| 27 | (+) | 1 | 1,284 |
| 28 | (+) | 1 | 1,282 |
| 29 | (+) | 1 | 1,237 |
| 3 | (+) | 1 | 1,287 |
| 30 | (+) | 1 | 1,295 |
| 31 | (+) | 1 | 1,284 |
| 32 | (+) | 1 | 1,283 |
| 33 | (+) | 1 | 1,278 |
| 34 | (+) | 1 | 1,305 |
| 35 | (+) | 1 | 1,272 |
| 36 | (+) | 1 | 1,304 |
| 37 | (+) | 1 | 1,274 |
| 38 | (+) | 1 | 1,290 |
| 38-1 | (+) | 1 | 1,298 |
| 39 | (+) | 1 | 1,275 |
| 4 | (+) | 1 | 1,287 |
| 40 | (+) | 1 | 1,280 |
| 41 | (+) | 1 | 1,289 |
| 42 | (+) | 1 | 1,293 |
| 43 | (+) | 1 | 1,287 |
| 44 | (+) | 1 | 1,281 |
| 45 | (+) | 1 | 1,276 |
| 46 | (+) | 1 | 1,271 |
| 47 | (+) | 1 | 1,279 |
| 48 | (+) | 1 | 1,290 |
| 49 | (+) | 1 | 1,225 |
| 5 | (+) | 1 | 1,278 |
| 50 | (+) | 1 | 1,277 |
| 51 | (+) | 1 | 1,293 |
| 52 | (+) | 1 | 1,287 |
| 53 | (+) | 1 | 1,284 |
| 54 | (+) | 1 | 1,291 |
| 55 | (+) | 1 | 1,284 |
| 58 | (+) | 1 | 1,284 |
| 6 | (+) | 1 | 1,274 |
| 60 | (+) | 1 | 1,280 |
| 67 | (+) | 1 | 1,275 |
| 7 | (+) | 1 | 1,286 |
| 76 | (+) | 1 | 1,276 |
| 8 | (+) | 1 | 1,279 |
| 83 | (+) | 1 | 1,251 |
| 84 | (+) | 1 | 1,275 |
| 9 | (+) | 1 | 1,292 |
| 90 | (+) | 1 | 1,277 |

### Supp data 5

NCBI Multiple Sequence Alignment Viewer, Version 1.26.0

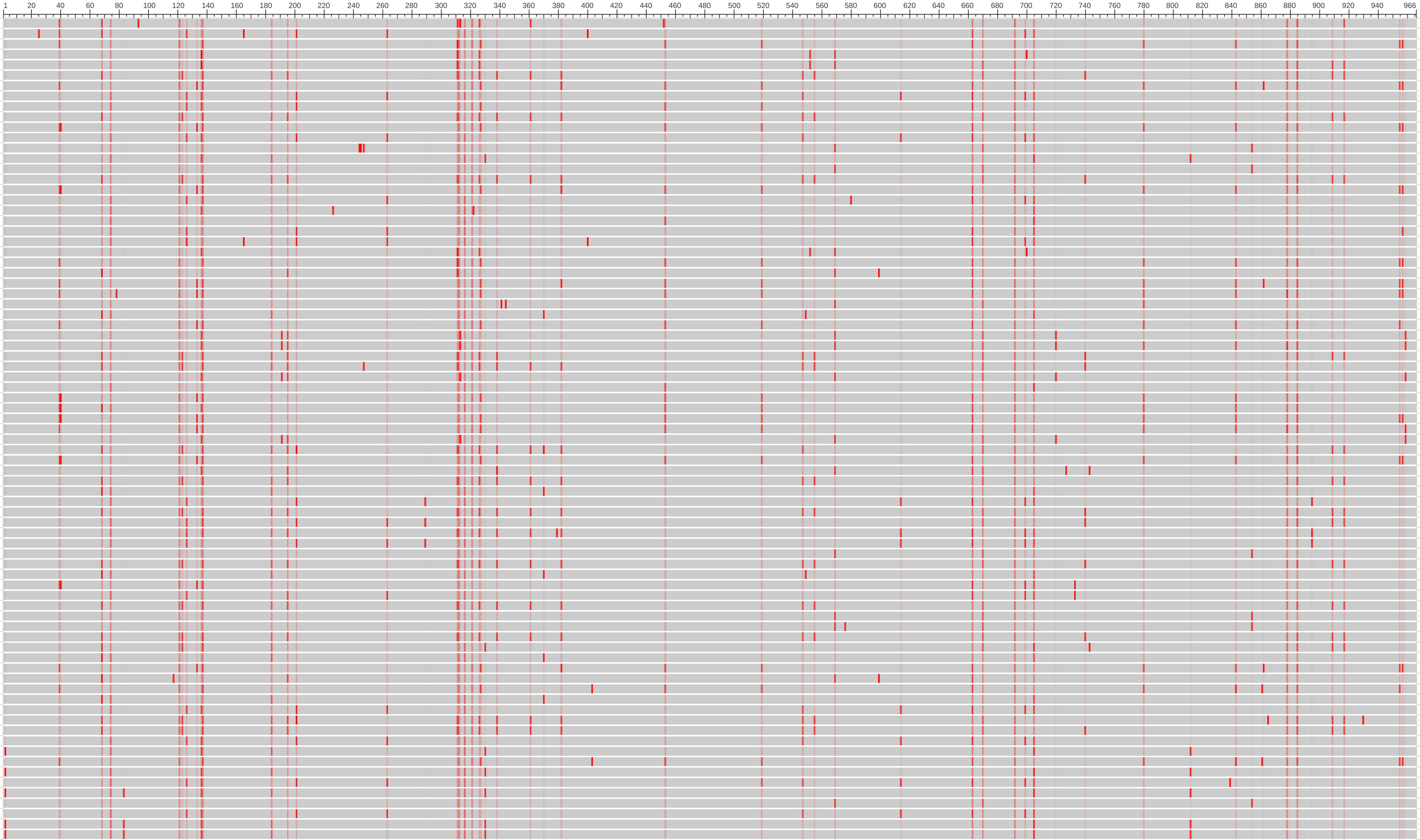


| Sequence ID | | Start | End |
|---|---|---|---|
| QHQ74301.1 | (+) | 1 | 966 |
| 29 | (+) | 1 | 966 |
| 30 | (+) | 1 | 966 |
| 58 | (+) | 1 | 966 |
| 58-1 | (+) | 1 | 966 |
| 42 | (+) | 1 | 966 |
| 33 | (+) | 1 | 966 |
| 105 | (+) | 1 | 966 |
| 105-1 | (+) | 1 | 966 |
| 49 | (+) | 1 | 966 |
| 37 | (+) | 1 | 966 |
| 23 | (+) | 1 | 966 |
| 51 | (+) | 1 | 966 |
| 7 | (+) | 1 | 966 |
| 52 | (+) | 1 | 966 |
| 43 | (+) | 1 | 966 |
| 36 | (+) | 1 | 966 |
| 22 | (+) | 1 | 966 |
| 13 | (+) | 1 | 966 |
| 19 | (+) | 1 | 966 |
| 21 | (+) | 1 | 966 |
| 28 | (+) | 1 | 966 |
| 67 | (+) | 1 | 966 |
| 67-1 | (+) | 1 | 966 |
| 5 | (+) | 1 | 966 |
| 34 | (+) | 1 | 966 |
| 40 | (+) | 1 | 966 |
| 2 | (+) | 1 | 966 |
| 14 | (+) | 1 | 966 |
| 41 | (+) | 1 | 966 |
| 110 | (+) | 1 | 966 |
| 110-1 | (+) | 1 | 966 |
| 111 | (+) | 1 | 966 |
| 111-1 | (+) | 1 | 966 |
| 3 | (+) | 1 | 966 |
| 20 | (+) | 1 | 966 |
| 112 | (+) | 1 | 966 |
| 112-1 | (+) | 1 | 966 |
| 38 | (+) | 1 | 966 |
| 113 | (+) | 1 | 966 |
| 113-1 | (+) | 1 | 966 |
| 50 | (+) | 1 | 966 |
| 39 | (+) | 1 | 966 |
| 4 | (+) | 1 | 966 |
| 47 | (+) | 1 | 966 |
| 18 | (+) | 1 | 966 |
| 116 | (+) | 1 | 966 |
| 116-1 | (+) | 1 | 966 |
| 117 | (+) | 1 | 966 |
| 117-1 | (+) | 1 | 966 |
| 79 | (+) | 1 | 966 |
| 53 | (+) | 1 | 966 |
| 45 | (+) | 1 | 966 |
| 15 | (+) | 1 | 966 |
| 118 | (+) | 1 | 966 |
| 118-1 | (+) | 1 | 966 |
| 48 | (+) | 1 | 966 |
| 54 | (+) | 1 | 966 |
| 55 | (+) | 1 | 966 |
| 120 | (+) | 1 | 966 |
| 120-1 | (+) | 1 | 966 |
| 16 | (+) | 1 | 966 |
| 35 | (+) | 1 | 966 |
| 6 | (+) | 1 | 966 |
| 31 | (+) | 1 | 966 |
| 17 | (+) | 1 | 966 |
| 27 | (+) | 1 | 966 |
| 46 | (+) | 1 | 966 |
| 44 | (+) | 1 | 966 |
| 24 | (+) | 1 | 966 |
| 8 | (+) | 1 | 966 |
| 32 | (+) | 1 | 966 |
| 9 | (+) | 1 | 966 |
| 26 | (+) | 1 | 966 |
| 10 | (+) | 1 | 966 |
| 1 | (+) | 1 | 966 |
| 25 | (+) | 1 | 966 |
| 12 | (+) | 1 | 966 |
| 11 | (+) | 1 | 966 |

### Supp data 6

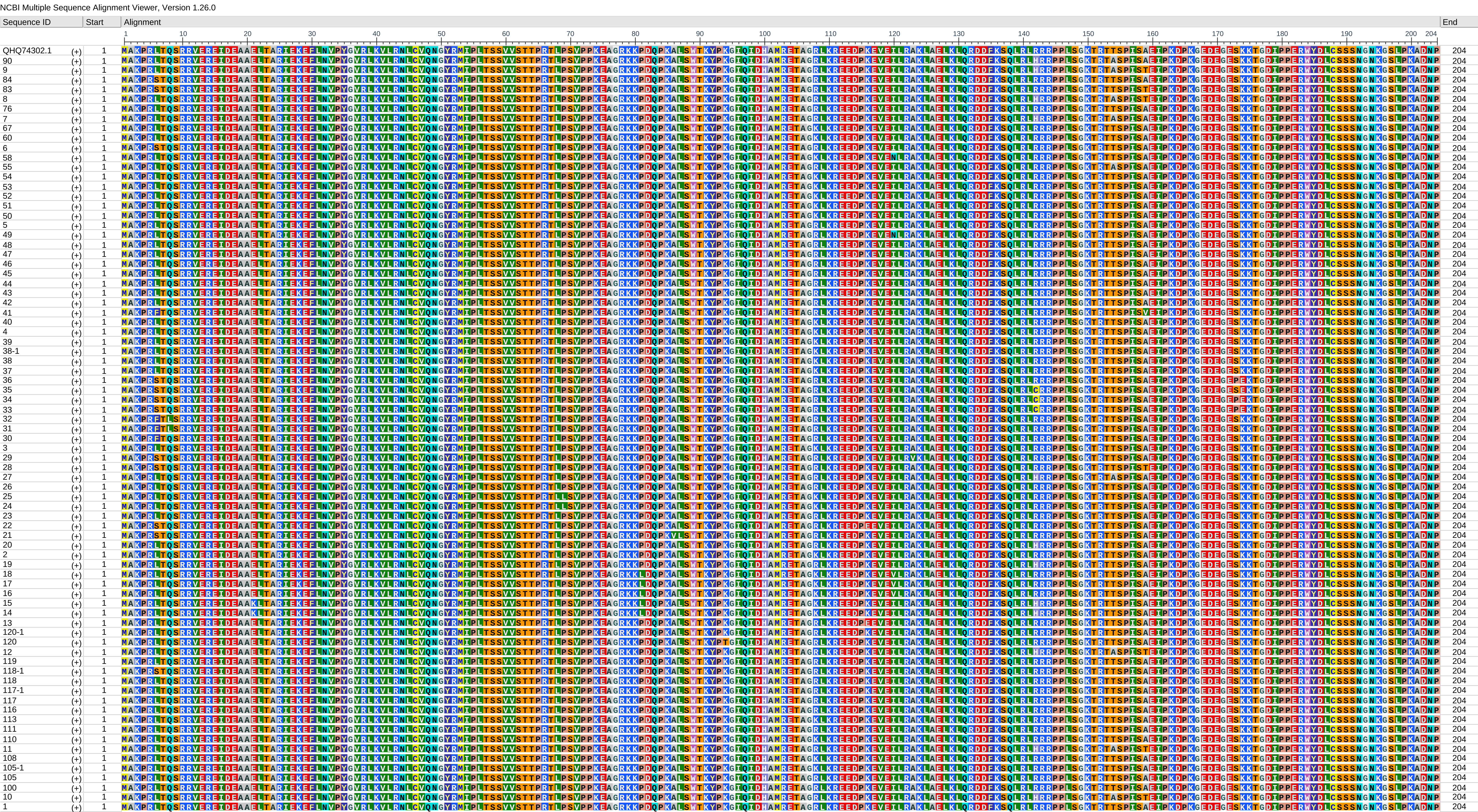
