## Supplementary material for "MaRNAV-1, an intracellular virus of *Plasmodium vivax*, is associated with increased parasite transmission and altered host immune response": Supp data 7

NCBI Multiple Sequence Alignment Viewer, Version 1.26.0

| Sequence ID | | Start | Alignment | End |
|---|---|---|---|---|
| QHQ74303.1 | (+) | 1 | MKGNRKRLGTFHLKGGTTSAALPTETKDLCRRQITLSCPLKFRLSVFEVKKEIDNLKLEESLHLDRRRRRNRKDFFRRPRDVAPDLRYRRINRKEIKALGFLSRLKLEEFSFREYKLVRYIYLKILPRVDLVLYRYVHQLVTFGVLQLLETYLSNRELCSKLN | 163 |
| 90 | (+) | 1 | MKGNRKRLGTFHLKGGTTSAALPMETKDLCRRQITLSCPLKFRLSVFEVKKEIDNLKLEESLHLDRRRRRNRKNFFRRPRDVVPDLRYRRINRKEIKALGFLSRLKLEEFSFREYKLVRYIYLKILPRVDLVLYRYVHQLVTFSVLQLLETYLSNRELCSKLN | 163 |
| 9 | (+) | 1 | MKGNRKRLGTFHLKGGTTSAALPMETKDLCRRQITLSCPLKFRLSVFEVKKEIDNLKLEESLHLDRRRRRNRKNFFRRPRDVVPDLRYRRINRKEIKALGFLSRLKLEEFSFREYKLVRYIYLKILPRVDLVLYRYVHQLVTFSVLQLLETYLSNRELCSKLN | 163 |
| 84 | (+) | 1 | MKGNRKRLGTFHLKGGTTSAALPMETKDLCRRQITLSCPLKFRLSVFEVKKEIDNLKLEESLHLDRRRRRNRKDFFRRPRDVVPDLRYRRINRKEIKALGFLSRLKLEEFSFREYKLVRYIYLKILPRVDLVLYRYVHQLVTFSVLQLLETYLSNRELCSKLN | 163 |
| 83 | (+) | 1 | MKGNRKRLGTFHLKGGTTSAALPMETKDLCRRQITLSCPLKFRLSVFEVKKEIDNLKLEESLHLDRRRRRNRKDFFRRPRDVVPDLRYRRINRKEIKALGFLSRLKLEEFSFREYKLVRYIYLKILPRVDLVLYRYVHQLVTFSVLQLLETYLSNRELCSKLN | 163 |
| 8 | (+) | 1 | MKGNRKRLGTFHLKGGTTSAALPMETKDLCRRQITLSCPLKFRLSVFEVKKEIDNLKLEESLHLDRRRRRNRKNFFRRPRDVVPDLRYRRINRKEIKALGFLSRLKLEEFSFREYKLVRYIYLKILPRVDLVLYRYVHQLVTFSVLQLLETYLSNRELCSKLN | 163 |
| 76 | (+) | 1 | MKGNRKRLGTFHLKGGTTSAALPMETKDLCRRQITLSCPLKFRLSVFEVKKEIDNLKLEESLHLDRRRRRNRKDFFRRPRDVVPDLRYRRINRKEIKALGFLSRLKLEEFSFREYKLVRYIYLKILPRVDLVLYRYVHQLVTFSVLQLLETYLSNRELCSKLN | 163 |
| 7 | (+) | 1 | MKGNRKRLGTFHLKGGTTSAALPMETKDLCRRQITLSCPLKFRLSVFEVKKEIDNLKLEESLHLDRRRRRNRKNFFRRPRDVVPDLRYRRINRKEIKALGFLSRLKLEEFSFREYKLVRYIYLKILPRVDLVLYRYVHQLVTFSVLQLLETYLSNRELCSKLN | 163 |
| 67 | (+) | 1 | MKGNRKRLGTFHLKGGTTSAALPMETKDLCRRQITLSCPLKFRLSVFEVKKEIDNLKLEESLHLDRRRRRNRKDFFRRPRDVVPDLRYRRINRKEIKALGFLSRLKLEEFSFREYKLVRYIYLKILPRVDLVLYRYVHQLVTFGVLQLLETYLSNRELCSKLN | 163 |
| 60 | (+) | 1 | MKGNRKRLGTFHLKGGTTSAALPMETKDLCRRQITLSCPLKFRLSVFEVKKEIDNLKLEESLHLDRRRRRNRKDFFRRPRDVVPDLRYRRINRKEIKALGFLSRLKLEEFSFREYKLVRYIYLKILPRVDLVLYRYVHQLVTFGVLQLLETYLSNRELCSKLN | 163 |
| 6 | (+) | 1 | MKGNRKRLGTFHLKGGTTSAALPMETKDLCRRQITLSCPLKFRLSVFEVKKEIDNLKLEESLHLDRRRRRNRKDFFRRPRDVAPDLRYRRINRKEIKALGFLSRLKLEEFSFREYKLVRYIYLKILPRVDLVLYRYVHQLVTFSVLQLLETYLSNRELCSKLN | 163 |
| 58 | (+) | 1 | MKGNRKRLGTFHLKGGTTSAALPMETKDLCRRQITLSCPLKFRLSVFEVKKEIDNLKLEESLHLDRRRRRNRKDFFRRPRDVVPDLRYRRINRKEIKALGFLSRLKLEEFSFREYKLVRYIYLKILPRVDLVLYRYVHQLVTFGVLQLLETYLSNRELCSKLN | 163 |
| 55 | (+) | 1 | MKGNRKRLGTFHLKGGTTSAALPMETKDLCRRQITLSCPLKFRLSVFEVKKEIDNLKLEESLHLDRRRRRNRKNFFRRPRDVVPDLRYRRINRKEIKALGFLSRLKLEEFSFREYKLVRYIYLKILPRVDLVLYRYVHQLVTFSVLQLLETYLSNRELCSKLN | 163 |
| 54 | (+) | 1 | MKGNRKRLGTFHLKGGTTSAALPMETKDLCRRQITLSCPLKFRLSVFEVKKEIDNLKLEESLHLDRRRRRNRKDFFRRPRDVVPDLRYRRINRKEIKALGFLSRLKLEEFSFREYKLVRYIYLKILPRVDLVLYRYVHQLVTFGVLQLLETYLSNRELCSKLN | 163 |
| 53 | (+) | 1 | MKGNRKRLGTFHLKGGTTSAALPMETKDLCRRQITLSCPLKFRLSVFEVKKEIDNLKLEESLHLDRRRRRNRKDFFRRPRDVVPDLRYRRINRKEIKALGFLSRLKLEEFSFREYKLVRYIYLKILPRVDLVLYRYVHQLVTFGVLQLLETYLSNRELCSKLN | 163 |
| 52 | (+) | 1 | MKGNRKRLGTFHLKGGTTSAALPMETKDLCRRQITLSCPLKFRLSVFEVKKEIDNLKLEESLHLDRRRRRNRKDFFRRPRDVVPDLRYRRINRKEIKALGFLSRLKLEEFSFREYKLVRYIYLKILPRVDLVLYRYVHQLVTFGVLQLLETYLSNRELCSKLN | 163 |
| 51 | (+) | 1 | MKGNRKRLGTFHLKGGTTSAALPMETKDLCRRQITLSCPLKFRLSVFEVKKEIDNLKLEESLHLDRRRRRNRKDFFRRPRDVVPDLRYRRINRKEIKALGFLSRLKLEEFSFREYKLVRYIYLKILPRVDLVLYRYVHQLVTFGVLQLLETYLSNRELCSKLN | 163 |
| 50 | (+) | 1 | MKGNRKRLGTFHLKGGTTSAALPMETKDLCRRQITLSCPLKFRLSVFEVKKEIDNLKLEESLHLDRRRRRNRKDFFRRPRDVVPDLRYRRINRKEIKALGFLSRLKLEEFSFREYKLVRYIYLKILPRVDLVLYRYVHQLVTFGVLQLLETYLSNRELCSKLN | 163 |
| 5 | (+) | 1 | MKGNRKRLGTFHLKGGTTSAALPMETKDLCRRQITLSCPLKFRLSVFEVKKEIDNLRLEESLHLDRRRRRNRKDFFRRPRDVAPDLRYRRINRKEIKALGFLSRLKLEEFSFREYKLVRYIYLKILPRVDLVLYRYVHQLVTFGVLQLLETYLSNRELCSKLN | 163 |
| 49 | (+) | 1 | MKGNRKRLGTFHLKGGTTSAALPTETKDLCRRQITLSCPLKFRLSVFEVKKEIDNLRLEESLHLDRRRRRNRKDFFRRPRDVVPDLRYRRINRKEIKALGFLSRLKLEEFSFREYKLVRYIYLKILPRVDLVLYRYVHQLVTFGVLQLLETYLSNRELCSKLN | 163 |
| 48 | (+) | 1 | MKGNRKRLGTFHLKGGTTSAALPTETKDLCRRQITLSCPLKFRLSVFEVKKEIDNLRLEESLHLDRRRRRNRKDFFRRPRDVVPDLRYRRINRKEIKALGFLSRLKLEEFSFREYKLVRYIYLKILPRVDLVLYRYVHQLVTFGVLQLLETYLSNRELCSKLN | 163 |
| 47 | (+) | 1 | MKGNRKRLGTFHLKGGTTSAALPTETKDLCRRQITLSCPLKFRLSVFEVKKEIDNLRLEESLHLDRRRRRNRKDFFRRPRDVVPDLRYRRINRKEIKALGFLSRLKLEEFSFREYKLVRYIYLKILPRVDLVLYRYVHQLVTFGVLQLLETYLSNRELCSKLN | 163 |
| 46 | (+) | 1 | MKGNRKRLGTFHLKGGTTSAALPTETKDLCRRQITLSCPLKFRLSVFEVKKEIDNLRLEESLHLDRRRRRNRKDFFRRPRDVVPDLRYRRINRKEIKALGFLSRLKLEEFSFREYKLVRYIYLKILPRVDLVLYRYVHQLVTFGVLQLLETYLSNRELCSKLN | 163 |
| 45 | (+) | 1 | MKGNRKRLGTFHLKGGTTSAALPTETKDLCRRQITLSCPLKFRLSVFEVKKEIDNLRLEESLHLDRRRRRNRKDFFRRPRDVVPDLRYRRINRKEIKALGFLSRLKLEEFSFREYKLVRYIYLKILPRVDLVLYRYVHQLVTFGVLQLLETYLSNRELCSKLN | 163 |
| 44 | (+) | 1 | MKGNRKRLGTFHLKGGTTSAALPTETKDLCRRQITLSCPLKFRLSVFEVKKEIDNLRLEESLHLDRRRRRNRKDFFRRPRDVVPDLRYRRINRKEIKALGFLSRLKLEEFSFREYKLVRYIYLKILPRVDLVLYRYVHQLVTFGVLQLLETYLSNRELCSKLN | 163 |
| 43 | (+) | 1 | MKGNRKRLGTFHLKGGTTSAALPTETKDLCRRQITLSCPLKFRLSVFEVKKEIDNLRLEESLHLDRRRRRNRKDFFRRPRDVVPDLRYRRINRKEIKALGFLSRLKLEEFSFREYKLVRYIYLKILPRVDLVLYRYVHQLVTFGVLQLLETYLSNRELCSKLN | 163 |
| 42 | (+) | 1 | MKGNRKRLGTFHLKGGTTSAALPTETKDLCRRQITLSCPLKFRLSVFEVKKEIDNLRLEESLHLDRRRRRNRKDFFRRPRDVVPDLRYRRINRKEIKALGFLSRLKLEEFSFREYKLVRYIYLKILPRVDLVLYRYVHQLVTFGVLQLLETYLSNRELCSKLN | 163 |
| 41 | (+) | 1 | MKGNRKRLGTFHLKGGMTSAALPMETKDLCRRQITLSCPLKFRLSVFEVKKEIDNLKLEESLHLDRRRRRNRKDFFRRPRDVAPDLRYRRINRKEIKALGFLSRLKLEEFSFREYKLVRYIYLKILPRVDLVLYRYVHQLVTFGVLQLLETYLSNRELCSKLN | 163 |
| 40 | (+) | 1 | MKGNRKRLGTFHLKGGTTSAALPMETKDLCRRQITLSCPLKFRLSVFEVKKEIDNLKLEESLHLDRRRRRNRKDFFRRPRDVVPDLRYRRINRKEIKALGFLSRLKLEEFSFREYKLVRYIYLKILPRVDLVLYRYVHQLVTFGVLQLLETYLSNRELCSKLN | 163 |
| 4 | (+) | 1 | MKGNRKRLGTFHLKGGTTSAALPMETKDLCRRQITLSCPLKFRLSVFEVKKEIDNLKLEESLHLDRRRRRNRKDFFRRPRDVVPDLRYRRINRKEIKALGFLSRLKLEEFSFREYKLVRYIYLKILPRVDLVLYRYVHQLVTFGVLQLLETYLSNRELCSKLN | 163 |
| 39 | (+) | 1 | MKGNRKRLGTFHLKGGTTSAALPMETKDLCRRQITLSCPLKFRLSVFEVKKEIDNLKLEESLHLDRRRRRNRKDFFRRPRDVVPDLRYRRINRKEIKALGFLSRLKLEEFSFREYKLVRYIYLKILPRVDLVLYRYVHQLVTFGVLQLLETYLSNRELCSKLN | 163 |
| 38-1 | (+) | 1 | MKGNRKRLGTFHLKGGTTSAALPMETKDLCRRQITLSCPLKFRLSVFEVKKEIDNLKLEESLHLDRRRRRNRKDFFRRPRDVVPDLRYRRINRKEIKALGFLSRLKLEEFSFREYKLVRYIYLKILPRVDLVLYRYVHQLVTFGVLQLLETYLSNRELCSKLN | 163 |
| 38 | (+) | 1 | MKGNRKRLGTFHLKGGTTSAALPMETKDLCRRQITLSCPLKFRLSVFEVKKEIDNLKLEESLHLDRRRRRNRKDFFRRPRDVVPDLRYRRINRKEIKALGFLSRLKLEEFSFREYKLVRYIYLKILPRVDLVLYRYVHQLVTFSVLQLLETYLSNRELCSKLN | 163 |
| 37 | (+) | 1 | MKGNRKRLGTFHLKGGTTSAALPMETKDLCRRQITLSCPLKFRLSVFEVKKEIDNLKLEESLHLDRRRRRNRKDFFRRPRDVVPDLRYRRINRKEIKALGFLSRLKLEEFSFREYKLVRYIYLKILPRVDLVLYRYVHQLVTFGVLQLLETYLSNRELCSKLN | 163 |
| 36 | (+) | 1 | MKGNRKRLGIFHLKGGTTSAALPMETKDLCRRQITLSCPLKFRLSVFEVKKEIDNLKLEESLHLDRRRRRNRKDFFRRPRDVVPDLRYRRINRKEIKALGFLSRLKLEEFSFREYKLVRYIYLKILPRVDLVLYRYVHQLVTFSVLQLLETYLSNRELCSKLN | 163 |
| 35 | (+) | 1 | MKGNRKRLGTFHLKGGTTSAALPMETKDLCRRQITLSCPLKFRLSVFEVKKEIDNLKLEESLHLDRRRRRNRKDFFRRPRDVVPDLRYRRINRKEIKALGFLSRLKLEEFSFREYKLVRYIYLKILPRVDLVLYRYVHQLVTFSVLQLLETYLSNRELCSRLN | 163 |
| 34 | (+) | 1 | MKGNRKRLGIFHLKGGTTSAALPMETKDLCRRQITLSCPLKFRLSVFEVKKEIDNLKLEESLHLDRRRRRNRKDFFRRPRDVVPDLRYRRINRKEIKALGFLSRLKLEEFSFREYKLVRYIYLKILPRVDLVLYRYVHQLVTFSVLQLLETYLSNRELCSRLN | 163 |
| 33 | (+) | 1 | MKGNRKRLGIFHLKGGTTSAALPMETKDLCRRQITLSCPLKFRLSVFEVKKEIDNLKLEESLHLDRRRRRNRKDFFRRPRDVVPDLRYRRINRKEIKALGFLSRLKLEEFSFREYKLVRYIYLKILPRVDLVLYRYVHQLVTFSVLQLLETYLSNRELCSRLN | 163 |
| 32 | (+) | 1 | MKGNRKRLGTFHLKGGTTSAALPTETKDLCRRQITLSCPLKFRLSVFEVKKEIDNLKLEESLHLDRRRRRNRKDFFRRPRDVAPDLRYRRINRKEIKALGFLSRLKLEEFSFREYKLVRYIYLKILPRVDLVLYRYVHQLVTFGVLQLLETYLSNRELCSKLN | 163 |
| 31 | (+) | 1 | MKGNRKRLGTFHLKGGTTSAALPTETKDLCRRQITLSCPLKFRLSVFEVKKEIDNLKLEESLHLDRRRRRNRKDFFRRPRDVAPDLRYRRINRKEIKALGFLSRLKLEEFSFREYKLVRYIYLKILPRVDLVLYRYVHQLVTFGVLQLLETYLSNRELCSKLN | 163 |
| 30 | (+) | 1 | MKGNRKRLGTFHLKGGMTSAALPMETKDLCRRQITLSCPLKFRLSVFEVKKEIDNLKLEESLHLDRRRRRNRKDFFRRPRDVAPDLRYRRINRKEIKALGFLSRLKLEEFSFREYKLVRYIYLKILPRVDLVLYRYVHQLVTFGVLQLLETYLSNRELCSKLN | 163 |
| 3 | (+) | 1 | MKGNRKRLGTFHLKGGTTSAALPMETKDLCRRQITLSCPLKFRLSVFEVKKEIDNLKLEESLHLDRRRRRNRKNFFRRPRDVVPDLRYRRINRKEIKALGFLSRLKLEEFSFREYKLVRYIYLKILPRVDLVLYRYVHQLVTFGVLQLLETYLSNRELCSRLN | 163 |
| 29 | (+) | 1 | MKGNRKRLGTFHLKGGTTSAALPMETKDLCRRQITLSCPLKFRLSVFEVKKEIDNLKLEESLHLDRRRRRNRKDFFRRPRDVVPDLRYRRINRKEIKALGFLSRLKLEEFSFREYKLVRYIYLKILPRVDLVLYRYVHQLVTFSVLQLLETYLSNRELCSKLN | 163 |
| 28 | (+) | 1 | MKGNRKRLGTFHLKGGTTSAALPMETKDLCRRQITLSCPLKFRLSVFEVKKEIDNLKLEESLHLDRRRRRNRKDFFRRPRDVVPDLRYRRINRKEIKALGFLSRLKLEEFSFREYKLVRYIYLKILPRVDLVLYRYVHQLVTFSVLQLLETYLSNRELCSKLN | 163 |
| 27 | (+) | 1 | MKGNRKRLGTFHLKGGTTSAALPMETKDLCRRQITLSCPLKFRLSVFEVKKEIDNLKLEESLHLDRRRRRNRKNFFRRPRDVVPDLRYRRINRKEIKALGFLSRLKLEEFSFREYKLVRYIYLKILPRVDLVLYRYVHQLVTFSVLQLLETYLSNRELCSKLN | 163 |
| 26 | (+) | 1 | MKGNRKRLGTFHLKGGTTSAALPMETKDLCRRQITLSCPLKFRLSVFEVKKEIDNLKLEESLHLDRRRRRNRKDFFRRPRDVVPDLRYRRINRKEIKALGFLSRLKLEEFSFREYKLVRYIYLKILPRVDLVLYRYVHQLVTFSVLQLLETYLSNRELCSKLN | 163 |
| 25 | (+) | 1 | MKGNRKRLGTFHLKGGTTSAALPMETKDLCRRQITLSCPLKFRLSVFEVKKEIDNLKLEESLHLDRRRRRNRKDFFRRPRDVVPDLRYRRINRKEIKALGFLSRLKLEEFSFREYKLVRYIYLKILPRVDLVLYRYVHQLVTFSVLQLLETYLSNRELCSKLN | 163 |
| 24 | (+) | 1 | MKGNRKRLGTFHLKGGTTSAALPMETKDLCRRQITLSCPLKFRLSVFEVKKEIDNLKLEESLHLDRRRRRNRKDFFRRPRDVVPDLRYRRINRKEIKALGFLSRLKLEEFSFREYKLVRYIYLKILPRVDLVLYRYVHQLVTFSVLQLLETYLSNRELCSKLN | 163 |
| 23 | (+) | 1 | MKGNRKRLGTFHLKGGTTSAALPMETKDLCRRQITLSCPLKFRLSVFEVKKEIDNLKLEESLHLDRRRRRNRKDFFRRPRDVVPDLRYRRINRKEIKALGFLSRLKLEEFSFREYKLVRYIYLKILPRVDLVLYRYVHQLVTFSVLQLLETYLSNRELCSKLN | 163 |
| 22 | (+) | 1 | MKGNRKRLGTFHLKGGTTSAALPTETKDLCRRQITLSCPLKFRLSVFEVKKEIDNLKLEESLHLDRRRRRNRKDFFRRPRDVVPDLRYRRINRKEIKALGFLSRLKLEEFSFREYKLVRYIYLKILPRVDLVLYRYVHQLVTFSVLQLLETYLSNRELCSKLN | 163 |
| 21 | (+) | 1 | MKGNRKRLGTFHLKGGTTSAALPMETKDLCRRQITLSCPLKFRLSVFEVKKEIDNLKLEESLHLDRRRRRNRKDFFRRPRDVVPDLRYRRINRKEIKALGFLSRLKLEEFSFREYKLVRYIYLKILPRVDLVLYRYVHQLVTFSVLQLLETYLSNRELCSKLN | 163 |
| 20 | (+) | 1 | MKGNRKRLGTFHLKGGTTSAALPMETRDLCRRQITLSCPLKFRLSVFEVKKEIDNLRLEESLHLDRRRRRNRKDFFRRPRDVAPDLRYRRINRKEIKALGFLSRLKLEEFSFREYKLVRYIYLKILPRVDLVLYRYVHQLVTFSVLQLLETYLSNRELCSKLN | 163 |
| 2 | (+) | 1 | MKGNRKRLGTFHLKGGTTSAALPMETKDLCRRQITLSCPLKFRLSVFEVKKEIDNLKLEESLHLDRRRRRNRKDFFRRPRDVVPDLRYRRINRKEIKALGFLSRLKLEEFSFREYKLVRYIYLKILPRVDLVLYRYVHQLVTFGVLQLLETYLSNRELCSKLN | 163 |
| 19 | (+) | 1 | MKGNRKRLGTFHLKGGTTSAALPMETRDLCRRQITLSCPLKFRLSVFEVKKEIDNLKLEESLHLDRRRRRNRKDFFRRPRDVAPDLRYRRINRKEIKALGFLSRLKLEEFSFREYKLVRYIYLKILPRVDLVLYRYVHQLVTFSVLQLLETYLSNRELCSKLN | 163 |
| 18 | (+) | 1 | MKGNRKRLGTFHLKGGTTSAALPMETKDLCRRQITLSCPLKFRLSVFEVKKEIDNLKLEESLHLDRRRRRNRKDFFRRPRDVVPDLRYRRINRKEIKALGFLSRLKLEEFSFREYKLVRYIYLKILPRVDLVLYRYVHQLVTFGVLQLLETYLSNRELCSKLN | 163 |
| 17 | (+) | 1 | MKGNRKRLGTFHLKGGTTSAALPMETKDLCRRQITLSCPLKFRLSVFEVKKEIDNLKLEESLHLDRRRRRNRKDFFRRPRDVVPDLRYRRINRKEIKALGFLSRLKLEEFSFREYKLVRYIYLKILPRVDLVLYRYVHQLVTFGVLQLLETYLSNRELCSKLN | 163 |
| 16 | (+) | 1 | MKGNRKRLGTFHLKGGTTSAALPMETKDLCRRQITLSCPLKFRLSVFEVKKEIDNLKLEESLHLDRRRRRNRKDFFRRPRDVVPDLRYRRINRKEIKALGFLSRLKLEEFSFREYKLVRYIYLKILPRVDLVLYRYVHQLVTFGVLQLLETYLSNRELCSKLN | 163 |
| 15 | (+) | 1 | MKGNRKRLGTFHLKGGTTSAALPMETKDLCRRQITLSCPLKFRLSVFEVKKEIDNLKLEESLHLDRRRRRNRKDFFRRPRDVVPDLRYRRINRKEIKALGFLSRLKLEEFSFREYKLVRYIYLKILPRVDLVLYRYVHQLVTFGVLQLLETYLSNRELCSKLN | 163 |
| 14 | (+) | 1 | MKGNRKRLGTFHLKGGTTSAALPMETKDLCRRQITLSCPLKFRLSVFEVKKEIDNLKLEESLHLDRRRRRNRKDFFRRPRDVVPDLRYRRINRKEIKALGFLSRLKLEEFSFREYKLVRYIYLKILPRVDLVLYRYVHQLVTFGVLQLLETYLSNRELCSKLN | 163 |
| 13 | (+) | 1 | MKGNRKRLGTFHLKGGTTSAALPMETKDLCRRQITLSCPLKFRLSVFEVKKEIDNLKLEESLHLDRRRRRNRKDFFRRPRDVVPDLRYRRINRKEIKALGFLSRLKLEEFSFREYKLVRYIYLKILPRVDLVLYRYVHQLVTFSVLQLLETYLSNRELCSKLN | 163 |
| 120-1 | (+) | 1 | MKGNRKRLGTFHLKGGTTSAALPTETKDLCRRQITLSCPLKFRLSVFEVKKEIDNLRLEESLHLDRRRRRNRKDFFRRPRDVVPDLRYRRINRKEIKALGFLSRLKLEEFSFREYKLVRYIYLKILPRVDLVLYRYVHQLVTFGVLQLLETYLSNRELCSKLN | 163 |
| 120 | (+) | 1 | MKGNRKRLGTFHLKGGTTSAALPTETKDLCRRQITLSCPLKFRLSVFEVKKEIDNLRLEESLHLDRRRRRNRKDFFRRPRDVVPDLRYRRINRKEIKALGFLSRLKLEEFSFREYKLVRYIYLKILPRVDLVLYRYVHQLVTFGVLQLLETYLSNRELCSKLN | 163 |
| 12 | (+) | 1 | MKGNRKRLGTFHLKGGTTSAALPMETKDLCRRQITLSCPLKFRLSVFEVKKEIDNLKLEESLHLDRRRRRNRKNFFRRPRDVVPDLRYRRINRKEIKALGFLSRLKLEEFSFREYKLVRYIYLKILPRVDLVLYRYVHQLVTFSVLQLLETYLSNRELCSKLN | 163 |
| 119 | (+) | 1 | MKGNRKRLGTFHLKGGTTSAALPTETKDLCRRQITLSCPLKFRLSVFEVKKEIDNLKLEESLHLDRRRRRNRKDFFRRPRDVVPDLRYRRINRKEIKALGFLSRLKLEEFSFREYKLVRYIYLKILPRVDLVLYRYVHQLVTFSVLQLLETYLSNRELCSKLN | 163 |
| 118-1 | (+) | 1 | MKGNRKRLGTFHLKGGTTSAALPTETKDLCRRQITLSCPLKFRLSVFEVKKEIDNLKLEESLHLDRRRRRNRKDFFRRPRDVVPDLRYRRINRKEIKALGFLSRLKLEEFSFREYKLVRYIYLKILPRVDLVLYRYVHQLVTFSVLQLLETYLSNRELCSKLN | 163 |
| 118 | (+) | 1 | MKGNRKRLGTFHLKGGTTSAALPTETKDLCRRQITLSCPLKFRLSVFEVKKEIDNLKLEESLHLDRRRRRNRKDFFRRPRDVVPDLRYRRINRKEIKALGFLSRLKLEEFSFREYKLVRYIYLKILPRVDLVLYRYVHQLVTFSVLQLLETYLSNRELCSKLN | 163 |
| 117-1 | (+) | 1 | MKGNRKRLGTFHLKGGTTSAALPTETKDLCRRQITLSCPLKFRLSVFEVKKEIDNLRLEESLHLDRRRRRNRKDFFRRPRDVVPDLRYRRINRKEIKALGFLSRLKLEEFSFREYKLVRYIYLKILPRVDLVLYRYVHQLVTFGVLQLLETYLSNRELCSKLN | 163 |
| 117 | (+) | 1 | MKGNRKRLGTFHLKGGTTSAALPTETKDLCRRQITLSCPLKFRLSVFEVKKEIDNLRLEESLHLDRRRRRNRKDFFRRPRDVVPDLRYRRINRKEIKALGFLSRLKLEEFSFREYKLVRYIYLKILPRVDLVLYRYVHQLVTFGVLQLLETYLSNRELCSKLN | 163 |
| 116 | (+) | 1 | MKGNRKRLGTFHLKGGTTSAALPTETKDLCRRQITLSCPLKFRLSVFEVKKEIDNLKLEESLHLDRRRRRNRKDFFRRPRDVVPDLRYRRINRKEIKALGFLSRLKLEEFSFREYKLVRYIYLKILPRVDLVLYRYVHQLVTFGVLQLLETYLSNRELCSKLN | 163 |
| 113 | (+) | 1 | MKGNRKRLGTFHLKGGTTSAALPMETKDLCRRQITLSCPLKFRLSVFEVKKEIDNLKLEESLHLDRRRRRNRKDFFRRPRDVVPDLRYRRINRKEIKALGFLSRLKLEEFSFREYKLVRYIYLKILPRVDLVLYRYVHQLVTFGVLQLLETYLSNRELCSKLN | 163 |
| 111 | (+) | 1 | MKGNRKRLGTFHLKGGTTSAALPTETKDLCRRQITLSCPLKFRLSVFEVKKEIDNLRLEESLHLDRRRRRNRKDFFRRPRDVVPDLRYRRINRKEIKALGFLSRLKLEEFSFREYKLVRYIYLKILPRVDLVLYRYVHQLVTFGVLQLLETYLSNRELCSKLN | 163 |
| 110 | (+) | 1 | MKGNRKRLGTFHLKGGTTSAALPMETKDLCRRQITLSCPLKFRLSVFEVKKEIDNLKLEESLHLDRRRRRNRKDFFRRPRDVVPDLRYRRINRKEIKALGFLSRLKLEEFSFREYKLVRYIYLKILPRVDLVLYRYVHQLVTFGVLQLLETYLSNRELCSKLN | 163 |
| 11 | (+) | 1 | MKGNRKRLGTFHLKGGTTSAALPMETKDLCRRQITLSCPLKFRLSVFEVKKEIDNLKLEESLHLDRRRRRNRKNFFRRPRDVVPDLRYRRINRKEIKALGFLSRLKLEEFSFREYKLVRYIYLKILPRVDLVLYRYVHQLVTFSVLQLLETYLSNRELCSKLN | 163 |
| 108 | (+) | 1 | MKGNRKRLGTFHLKGGTTSAALPMETKDLCRRQITLSCPLKFRLSVFEVKKEIDNLKLEESLHLDRRRRRNRKDFFRRPRDVVPDLRYRRINRKEIKALGFLSRLKLEEFSFREYKLVRYIYLKILPRVDLVLYRYVHQLVTFGVLQLLETYLSNRELCSKLN | 163 |
| 105-1 | (+) | 1 | MKGNRKRLGTFHLKGGTTSAALPMETKDLCRRQITLSCPLKFRLSVFEVKKEIDNLKLEESLHLDRRRRRNRKDFFRRPRDVVPDLRYRRINRKEIKALGFLSRLKLEEFSFREYKLVRYIYLKILPRVDLVLYRYVHQLVTFSVLQLLETYLSNRELCSKLN | 163 |
| 105 | (+) | 1 | MKGNRKRLGTFHLKGGTTSAALPMETKDLCRRQITLSCPLKFRLSVFEVKKEIDNLKLEESLHLDRRRRRNRKDFFRRPRDVVPDLRYRRINRKEIKALGFLSRLKLEEFSFREYKLVRYIYLKILPRVDLVLYRYVHQLVTFSVLQLLETYLSNRELCSKLN | 163 |
| 100 | (+) | 1 | MKGNRKRLGTFHLKGGTTSAALPTETKDLCRRQITLSCPLKFRLSVFEVKKEIDNLRLEESLHLDRRRRRNRKDFFRRPRDVVPDLRYRRINRKEIKALGFLSRLKLEEFSFREYKLVRYIYLKILPRVDLVLYRYVHQLVTFGVLQLLETYLSNRELCSKLN | 163 |
| 10 | (+) | 1 | MKGNRKRLGTFHLKGGTTSAALPMETKDLCRRQITLSCPLKFRLSVFEVKKEIDNLKLEESLHLDRRRRRNRKNFFRRPRDVVPDLRYRRINRKEIKALGFLSRLKLEEFSFREYKLVRYIYLKILPRVDLVLYRYVHQLVTFSVLQLLETYLSNRELCSKLN | 163 |
| 1 | (+) | 1 | MKGNRKRLGTFHLKGGTTSAALPMETKDLCRRQITLSCPLKFRLSVFEVKKEIDNLKLEESLHLDRRRRRNRKDFFRRPRDVVPDLRYRRINRKEIKALGFLSRLKLEEFSFREYKLVRYIYLKILPRVDLVLYRYVHQLVTFSVLQLLETYLSNRELCSKLN | 163 |
