## Supplementary material for "MaRNAV-1, an intracellular virus of *Plasmodium vivax*, is associated with increased parasite transmission and altered host immune response": Supp figures and tables

**Supplementary information**

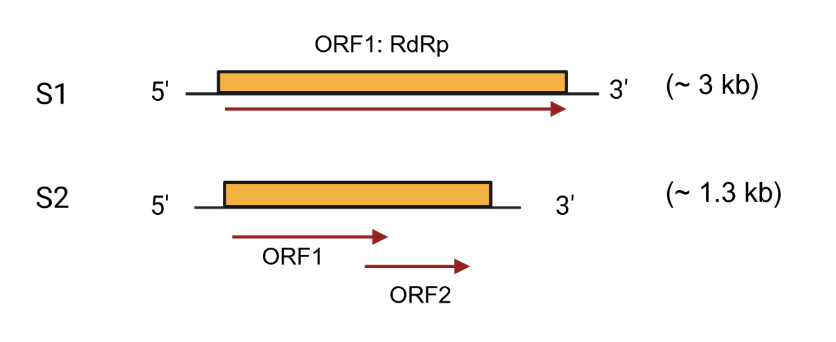

**Supp. Fig. 1.** **Genomic structure of MaRNAV-1.** MaRNAV-1 is bisegmented: the first segment (S1) encodes an RNA-dependent RNA polymerase (RdRp) of 966 aa (GenBank accession No. MN860568.1) that shares a conserved motif with the polymerase of fungal narnaviruses. The second segment (S2) encodes two overlapping open reading frames (ORF1 with 204 aa and ORF2 with 163 aa), possibly encoding proteins with no homology with annotated proteins (GenBank accession No. MN860569.1).

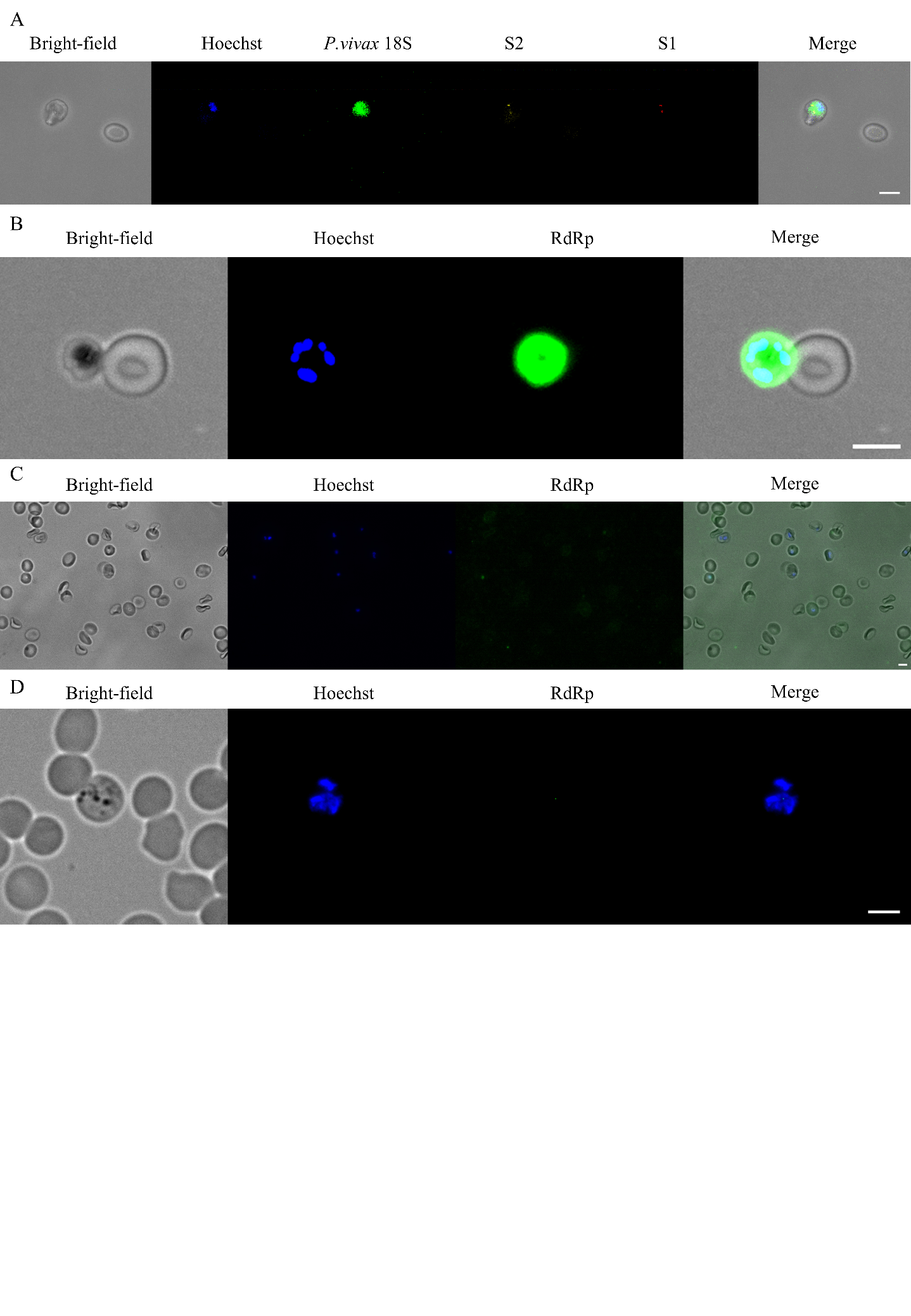

**Supp. Fig. 2**. **smiFISH and immunofluorescence assay (IFA) to detect MaRNAV-1 RNA and RdRp**. **A**. smiFISH performed on a MaRNAV-1-positive *P. vivax* infection. No viral signal was detected in the adjacent uninfected red blood cell. **B**. IFA performed *P. vivax* infection positive for MaRNAV-1. There is no signal of the RdRp protein in the adjacent uninfected red blood cell. **C**. IFA performed on *P. falciparum* 3D7-infected erythrocytes using anti-MaRNAV-1 RdRp antibodies. No RdRp signal was detected, confirming the antibody's specificity for *P. vivax* MaRNAV-1-infected parasites. **D**. IFA performed on *P. vivax-*infected reticulocytes using pre-immune rabbit IgG as a control. No fluorescence signal was detected. DNA was stained with Hoechst (blue). *P. vivax* 18S rRNA labeled with FLAP-Y-Alexa Fluor 488 (green); MaRNAV-1 S2 RNA labeled with FLAP-Y-Cy3 (yellow); MaRNAV-1 S1 RNA labeled with FLAP-Y-Cy5 (red). Goat anti-rabbit Alexa Fluor 488 secondary antibodies were used for the detection of the RdRp protein. Scale bar, 5 µm.

**Supp. Table 1.** **Clearance kinetics of *P. vivax* parasites and MaRNAV-1 following artesunate treatment**.

| Day follow-up | *P. vivax* | MaRNAV-1 |
| --- | --- | --- |
| D0 | 20/20 | 20/20 |
| D1 | 16/20 | 19/20 |
| D2 | 9/20 | 12/20 |
| D3 | 1/20 | 4/20 |
| D4 | 0/20 | 2/20 |
| D5 | 0/20 | 0/20 |
| D6 | 0/20 | 0/20 |
| D7 | 0/20 | 0/20 |

Counts represent the number of patients with detectable *P. vivax* or MaRNAV-1 RNA in peripheral blood at each follow-up day after artesunate treatment initiation (Day 0). *P. vivax* detection was performed by RT-qPCR targeting the 18S rRNA gene, while MaRNAV-1 detection was performed by RT-qPCR targeting viral RNA. Values are shown as number of positive samples over the total number of patients tested (n = 20). Parasite and viral RNA were monitored daily until Day 7 following treatment initiation.

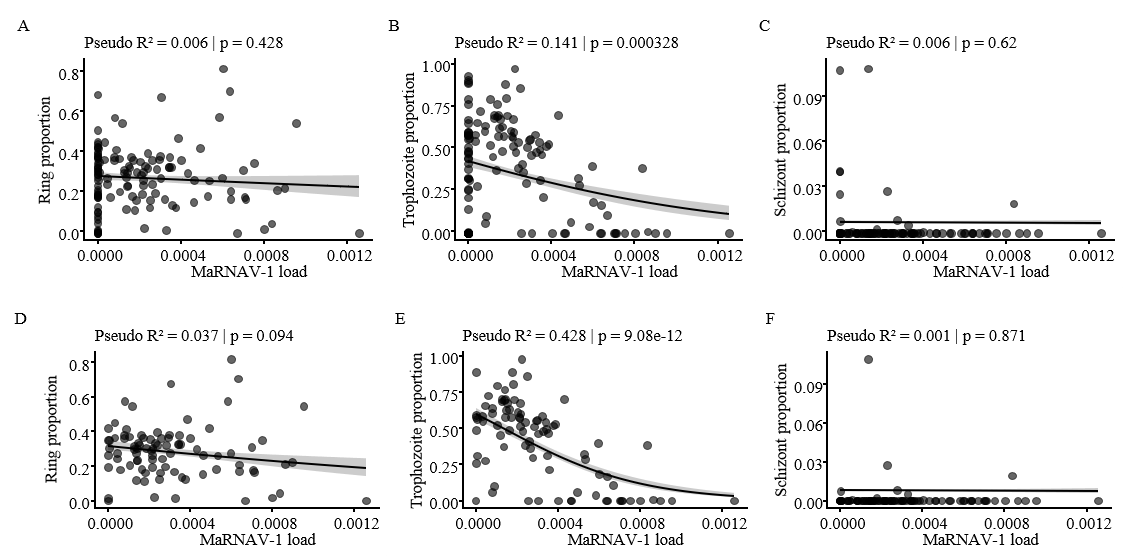

**Supp. Fig. 3.** **Associations between MaRNAV-1 load and blood stage parasite composition and gametocytemia by RNA-seq.** (**A-C)** Beta regression analyses assessing the association between MaRNAV-1 load and the proportion of blood stage parasite developmental stages (ring, trophozoite, and schizont), including both virus-negative and virus-positive samples (n = 126). Lines represent fitted values from beta regression models with shaded 95% confidence intervals (CI). Pseudo-R² and p-values are shown in each panel. **(D-F)** Same analyses restricted to MaRNAV-1-positive samples only (n = 88). In all panels, the x-axis represents MaRNAV-1 load, and the y-axis indicates the proportion of each parasite developmental stage (ring, trophozoite, schizont). Lines represent fitted values from beta regression models with shaded 95% CI. Pseudo-R² and p-values are shown in each panel.

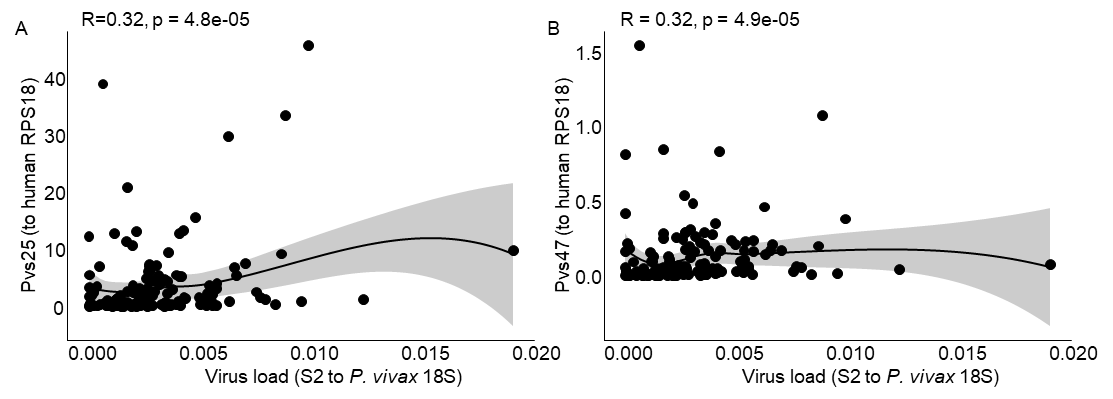

**Supp. Fig 4. Spearman correlation between MaRNAV-1 load and gametocytemia by RT-qPCR.** Virus load is measured by RT-qPCR as S2 expression normalized to *P. vivax* 18S rRNA, and gametocytemia is measured as **(A)** Pvs25 RNA relative abundance (normalized to human RPS18, n=152) or **(B)** Pvs47 (normalized to human RPS18, n=152). The black line represents a LOESS-smoothed fit, with the shaded area indicating 95% CI.

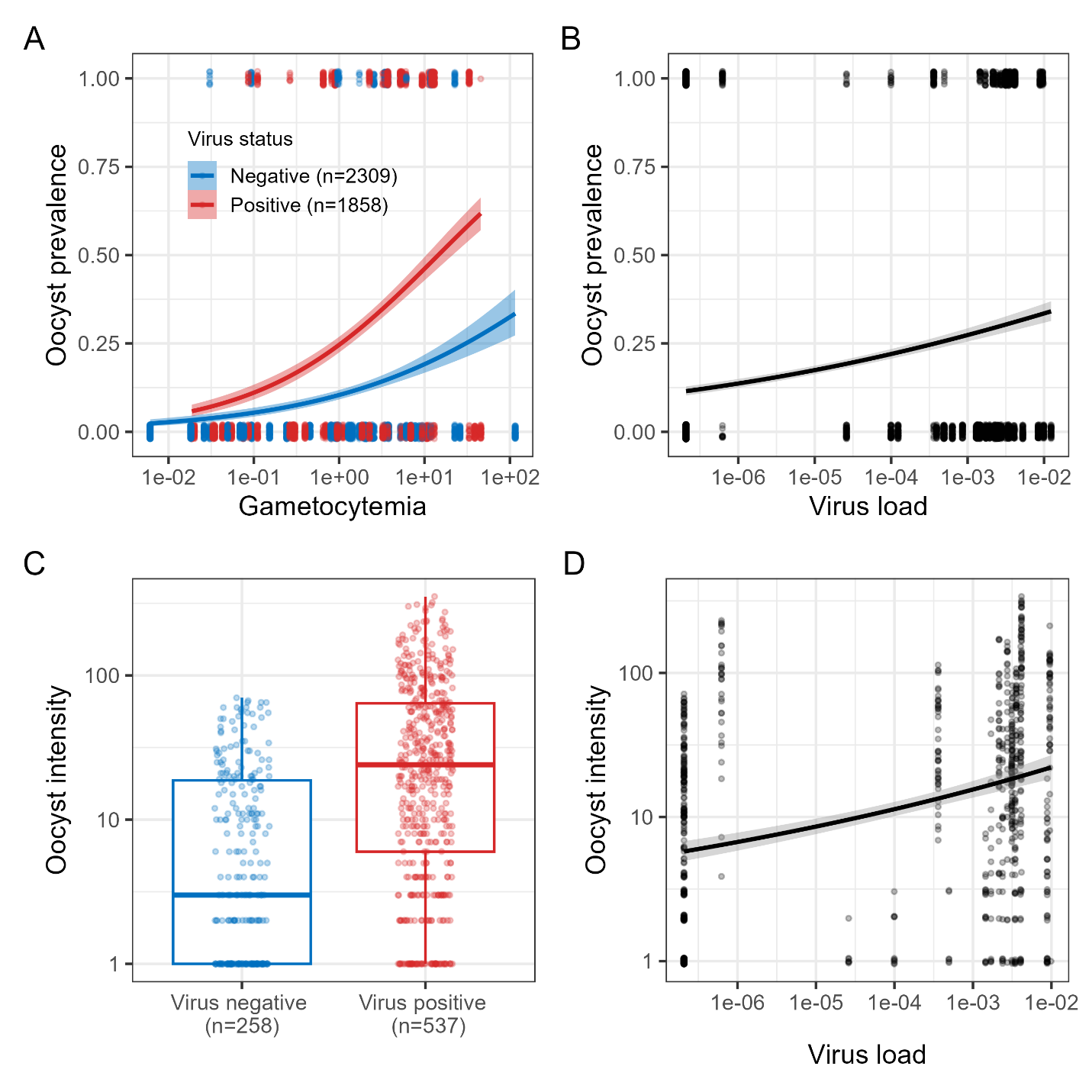

**Supp. Fig. 5**. **Association between MaRNAV-1 infection status or virus load and transmission of *P. vivax* to mosquito vectors.** Gametocytemia was quantified by RT-qPCR as the relative expression of Pvs25 normalized to the human RPS18 gene, and MaRNAV-1 load was quantified by RT-qPCR as the relative abundance of S2 normalized to *P. vivax* 18S rRNA. **A.** Association between gametocytemia and oocyst prevalence, stratified by MaRNAV-1 infection status (number of mosquitoes: n = 2309 MaRNAV-1-negative vs. n = 1858 MaRNAV-1-positive). Lines represent model-predicted probabilities with 95% confidence intervals (CI). **B.** Association between MaRNAV-1 load and oocyst prevalence. **C.** Oocyst intensity (number of oocysts per infected mosquito) according to MaRNAV-1 infection status (number of infected mosquitoes: n = 258 MaRNAV-1-negative vs. n = 537 MaRNAV-1). Boxplots show the median and interquartile range, with individual data points overlaid. **D.** Association between MaRNAV-1 load and oocyst intensity. Using MaRNAV-1 as a binary variable, the presence of MaRNAV-1 was positively associated with both oocyst prevalence (panel A) and oocyst intensity (panel C) ($\chi_{1}^{2}$= 5.2, p = 0.02; $\chi_{1}^{2}$= 8.7, p = 0.003, respectively). Gametocytemia was also positively associated with both oocyst prevalence and intensity ($\chi_{1}^{2}$= 10.5, p = 0.001, $\chi_{1}^{2}$= 5.4, p = 0.0001, respectively), whereas no significant interaction between MaRNAV-1 status and gametocytemia was detected ($\chi_{1}^{2}$= 2, p = 0.16; $\chi_{1}^{2}$= 0.7, p = 0.41, respectively). When the MaRNAV-1 load was analyzed as a continuous variable, similar associations were observed. Virus load was positively associated with oocyst prevalence (panel B) and intensity (panel D) ($\chi_{1}^{2}$= 4.1, p = 0.04$; \chi_{1}^{2}$= 8.5, p = 0.003, respectively), as was gametocytemia ($\chi_{1}^{2}$= 10, p = 0.001; $\chi_{1}^{2}$= 13.7, p = 0.0002, respectively), but with no evidence of interaction between virus load and gametocytemia ($\chi_{1}^{2}$= 1.5, p = 0.23; $\chi_{1}^{2}$= 3.7, p = 0.06, respectively).

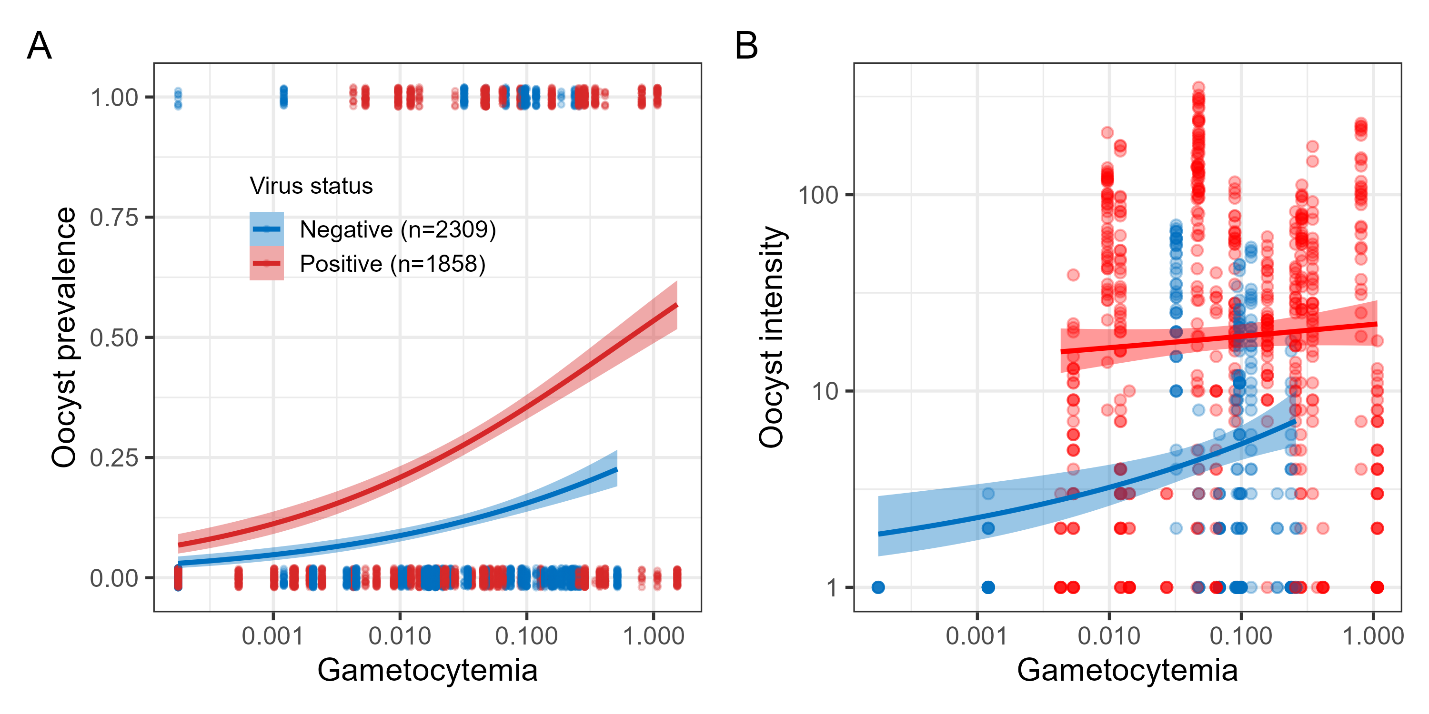

**Supp. Fig 6. Association between MaRNAV-1 infection status or virus load and transmission of *P. vivax* to mosquito vectors using Pvs47 as a gametocytemia measure.** Gametocytemia was quantified by qPCR as the relative expression of Pvs47 normalized to the human RPS18 gene, and MaRNAV-1 load was quantified by RT-qPCR as the relative abundance of S2 normalized to *P. vivax* 18S rRNA. **A.** Association between gametocytemia and oocyst prevalence, stratified by MaRNAV-1 infection status (number of mosquitoes: n = 2309 MaRNAV-1-negative vs n =1858 MaRNAV-1-positive). Lines represent model-predicted probabilities with 95% confidence intervals (CI). **B.** Association between gametocytemia and oocyst intensity, stratified by MaRNAV-1 infection status. Lines represent model fits with 95% CI. The presence of MaRNAV-1 was positively associated with both oocyst prevalence and oocyst intensity **(**$\chi_{1}^{2}$= 4.8, p = 0.03; $\chi_{1}^{2}$=6.8, p=0.009, respectively, panel A &B). Gametocytemia was positively associated with oocyst prevalence ($\chi_{1}^{2}$= 8.5, p = 0.003) and showed a marginal association with oocyst intensity ($\chi_{1}^{2}$ = 2.9, p = 0.09). No significant interaction between MaRNAV-1 status and gametocytemia was detected ($\chi_{1}^{2}$=1.6, p = 0.2, $\chi_{1}^{2}$ = 1.1, p = 0.3, respectively). MaRNAV-1 load was positively associated with oocyst prevalence and intensity ($\chi_{1}^{2}$= 4.1, p = 0.04, $\chi_{1}^{2}$ = 7.9, p = 0.004, respectively, Supp. Fig 5B & 5D). Gametocytemia remained positively associated with oocyst prevalence ($\chi_{1}^{2}$ = 8.7, p = 0.003) and showed a borderline association with oocyst intensity ($\chi_{1}^{2}$ = 3.5, p = 0.051). No significant interaction between virus load and gametocytemia was observed ($\chi_{1}^{2}$ = 0.93, p = 0.33, $\chi_{1}^{2}$= 3.17, p = 0.07, respectively).

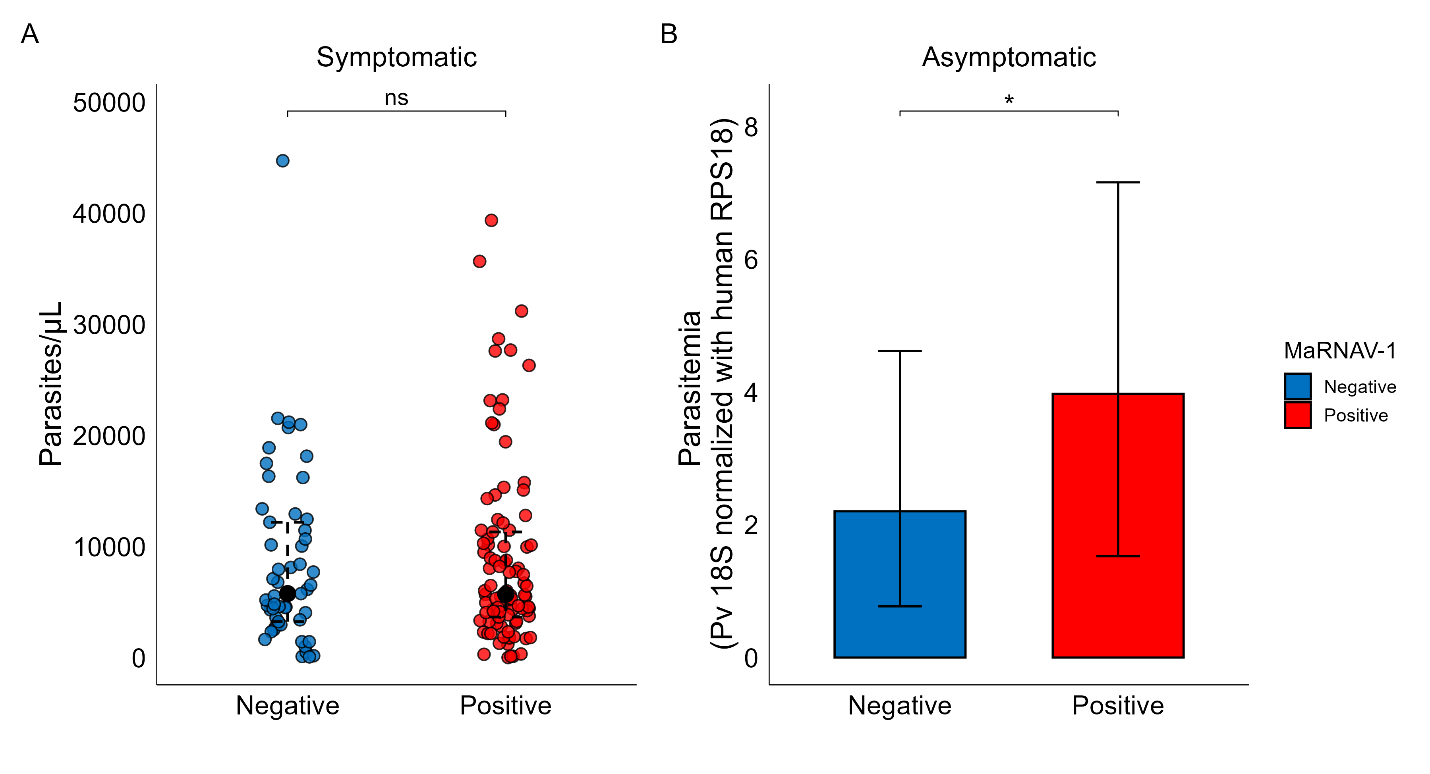

**Supp. Fig. 7**. **Comparison of parasitemia levels between MaRNAV-1-negative and positive samples in symptomatic and asymptomatic Cambodian *P. vivax*-infected patients**. **A**. No significant difference in parasitemia (determined by thick film light microscopy) among symptomatic *P. vivax*-infected individuals between MaRNAV-1-positive and negative infections (n _negative_ = 53, n _positive_ = 93, p = 0.8). **B**. Parasitemia (determined by RT-qPCR relative to human RPS18) in asymptomatic *P. vivax*-infected individuals is lower in MaRNAV-1 negative infections compared to positive ones (n _negative_ = 107, n _positive_ = 49, p = 0.02)

**Supp. Table 2.** **Association between MaRNAV-1 infection status and clinical correlates in *P. vivax*-infected patients.** Associations were assessed using linear regression models including parasitemia (log-transformed) and MaRNAV-1 infection status (presence vs absence) as covariates (n = 125; 87 virus-positive and 38 virus-negative samples). MaRNAV-1 infection status was not significantly associated with any clinical parameter after correction for multiple testing. In contrast, parasitemia was significantly associated with several clinical outcomes (FDR-adjusted p < 0.05).

| Outcome | Predictor | Estimate | CI_lower | CI_upper | Chisq | df | p_value | p_FDR |
| --- | --- | --- | --- | --- | --- | --- | --- | --- |
| Basophil counts | MaRNAV-1_status | 0.01 | -0.36 | 0.38 | 0.00 | 1 | 0.9576 | 0.9576 |
|  | log(Parasitemia) | -0.11 | -0.27 | 0.05 | 1.81 | 1 | 0.1619 | 0.3643 |
| CRP | MaRNAV-1_status | -0.03 | -0.39 | 0.32 | 0.03 | 1 | 0.8566 | 0.9576 |
|  | log(Parasitemia) | 0.07 | -0.08 | 0.22 | 0.76 | 1 | 0.3446 | 0.5048 |
| Eosinophil counts | MaRNAV-1_status | 0.60 | 0.12 | 1.08 | 9.42 | 1 | 0.0153 | 0.1373 |
|  | log(Parasitemia) | -0.11 | -0.32 | 0.09 | 1.84 | 1 | 0.2792 | 0.5026 |
| Hemoglobin | MaRNAV-1_status | 0.14 | -0.51 | 0.79 | 0.52 | 1 | 0.6697 | 0.9576 |
|  | log(Parasitemia) | 0.01 | -0.27 | 0.28 | 0.00 | 1 | 0.9691 | 0.9691 |
| Lymphocyte counts | MaRNAV-1_status | 0.11 | -0.05 | 0.26 | 0.29 | 1 | 0.1936 | 0.8509 |
|  | log(Parasitemia) | -0.03 | -0.10 | 0.04 | 0.13 | 1 | 0.3927 | 0.5048 |
| Monocyte counts | MaRNAV-1_status | 0.02 | -0.21 | 0.26 | 0.01 | 1 | 0.8492 | 0.9576 |
|  | log(Parasitemia) | -0.16 | -0.26 | -0.05 | 3.51 | 1 | 0.0028 | 0.0126 |
| Neutrophil counts | MaRNAV-1_status | -0.03 | -0.29 | 0.23 | 0.02 | 1 | 0.8279 | 0.9576 |
|  | log(Parasitemia) | -0.01 | -0.12 | 0.10 | 0.01 | 1 | 0.8812 | 0.9691 |
| Platelet counts | MaRNAV-1_status | 0.06 | -0.19 | 0.32 | 0.11 | 1 | 0.6197 | 0.9576 |
|  | log(Parasitemia) | -0.14 | -0.25 | -0.03 | 2.76 | 1 | 0.0126 | 0.0379 |
| Temperature | MaRNAV-1_status | 0.19 | -0.16 | 0.55 | 0.98 | 1 | 0.2836 | 0.8509 |
|  | log(Parasitemia) | 0.27 | 0.12 | 0.43 | 10.86 | 1 | 0.0005 | 0.0044 |

**Supp. Table. 3**. **Non-linear associations between MaRNAV-1 load and clinical correlates in MaRNAV-1-positive *P. vivax*-infected patients.** Associations were assessed using generalized additive models (GAMs) including smooth terms for log-transformed MaRNAV-1 load and parasitemia (n = 87). MaRNAV-1 load was significantly associated with temperature (FDR-adjusted p = 0.010), indicating a non-linear relationship between virus load and body temperature. In addition, significant interaction effects between MaRNAV-1 load and parasitemia were observed for neutrophil counts and temperature (FDR-adjusted p = 0.036), suggesting that the effect of virus load on these parameters depends on parasite density. Parasitemia remained significantly associated with several clinical parameters, including eosinophil, monocyte and platelet counts, and temperature (FDR-adjusted p < 0.05).

| Outcome | Predictor | edf | F | p_value | p_FDR |
| --- | --- | --- | --- | --- | --- |
| Basophil counts | s(log(Parasitemia)) | 1.45 | 1.17 | 0.221 | 0.344 |
|  | s(log(MaRNAV-1 load)) | 1.00 | 0.11 | 0.737 | 0.830 |
| CRP | s(log(Parasitemia)) | 4.43 | 1.36 | 0.230 | 0.344 |
|  | s(log(MaRNAV-1 load)) | 3.20 | 1.58 | 0.208 | 0.286 |
| Eosinophil counts | s(log(Parasitemia)) | 8.33 | 3.92 | 0.001 | 0.005 |
|  | s(log(MaRNAV-1 load)) | 1.00 | 2.26 | 0.137 | 0.286 |
| Hemoglobin | s(log(Parasitemia)) | 1.00 | 0.04 | 0.836 | 0.836 |
|  | s(log(MaRNAV-1 load)) | 1.00 | 1.51 | 0.222 | 0.286 |
| Lymphocyte counts | s(log(Parasitemia)) | 1.00 | 0.52 | 0.474 | 0.610 |
|  | s(log(MaRNAV-1 load)) | 1.62 | 1.56 | 0.219 | 0.286 |
| Monocyte counts | s(log(Parasitemia)) | 1.56 | 3.91 | 0.018 | 0.040 |
|  | s(log(MaRNAV-1 load)) | 1.00 | 0.00 | 0.994 | 0.994 |
| Neutrophil counts | s(log(Parasitemia)) | 3.90 | 0.61 | 0.675 | 0.760 |
|  | s(log(MaRNAV-1 load)) | 1.79 | 1.67 | 0.177 | 0.286 |
|  | ti(log(MaRNAV-1 load),log(Parasitemia)) | 5.46 | 2.60 | 0.018 | 0.036 |
| Platelet counts | s(log(Parasitemia)) | 7.25 | 3.14 | 0.005 | 0.024 |
|  | s(log(MaRNAV-1 load)) | 3.95 | 1.54 | 0.181 | 0.286 |
| Temperature | s(log(Parasitemia)) | 1.05 | 5.88 | 0.013 | 0.040 |
|  | s(log(MaRNAV-1 load)) | 2.40 | 5.98 | 0.001 | 0.010 |
|  | ti(log(MaRNAV-1_load),log(Parasitemia)) | 6.57 | 2.18 | 0.036 | 0.036 |

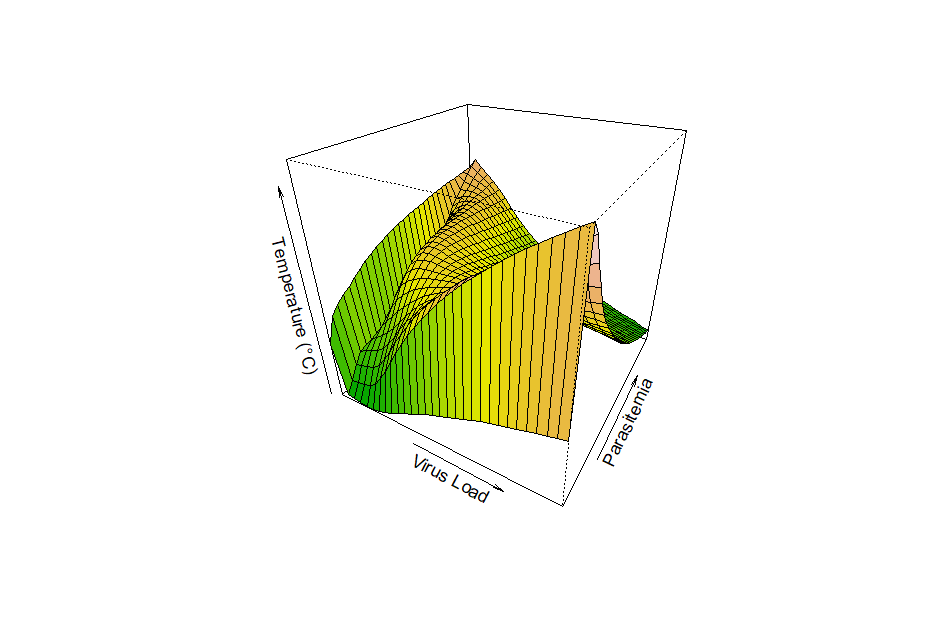

**Supp. Fig. 8.** **Interaction between MaRNAV-1 load and parasitemia on predicted body temperature.**
Three-dimensional model surface illustrating the joint effect of virus load and parasitemia on predicted body temperature. The surface represents fitted values from the generalized additive model, including an interaction term (see Supp. Table 3). Warmer colors indicate higher predicted temperatures. Axes correspond to MaRNAV-1 load, parasitemia, and temperature (°C), highlighting the non-linear and interactive effects of virus load and parasite density on host body temperature.

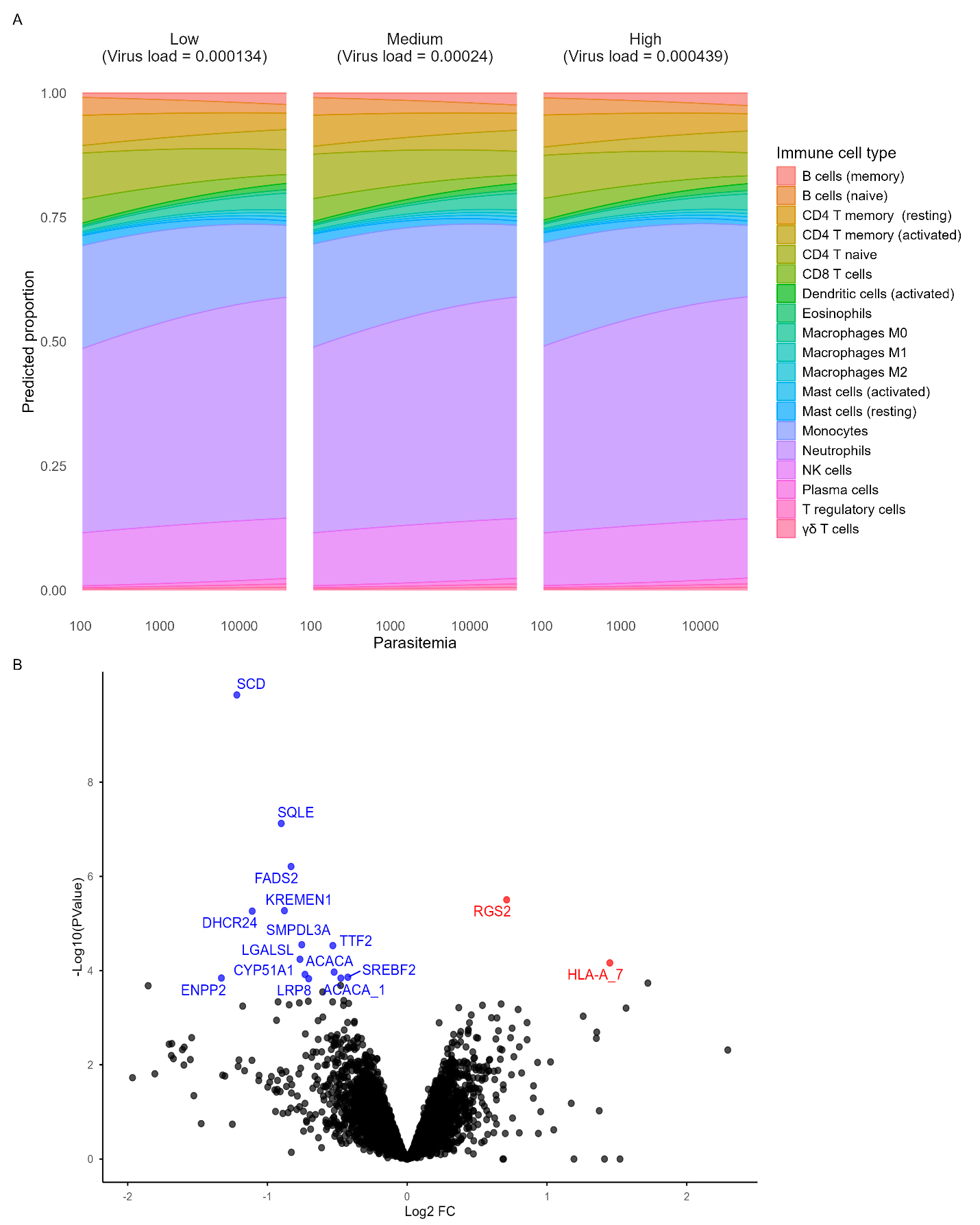

**Supp. Fig. 9. Host gene expression analysis between high and low MaRNAV-1 load in *P. vivax*-infected patients.** Volcano plot showing differentially expressed human genes between infections with low and high virus load (below the 25^th^ vs. above the 75^th^ percentile). Each point represents a gene, plotted by log₂ fold change and −log₁₀(p-value). Blue points indicate human genes that were downregulated in high-virus load, and red points indicate human genes that were upregulated in high virus load samples (FDR ≤ 0.1), with annotations for differentially expressed human genes. Several genes involved in lipid metabolism (*SQLE, SCD, FADS2*, and *ACACA*) and biosynthesis pathways (*DHCR24* and *SMPDL3A*) were down-regulated. Additionally, *LGALSL*, a gene structurally related to the galectin family and implicated in immune regulatory processes, and signaling factors (*KREMEN1* and *TTF2*) were significantly down-regulated in high MaRNAV-1 load samples. Conversely, *RGS2*, a regulator of G-protein receptor signaling, and *HLA-A*, a classical MHC class I molecule responsible for presenting intracellular antigens, were both up-regulated.

**Supp. Table 4. Global effects of MaRNAV-1 infection, parasitemia, and their interaction on immune cell composition.** Global associations with immune cell composition were assessed using Dirichlet regression models. Analyses were performed using MaRNAV-1 infection status as a binary variable (negative vs. positive, n = 125). Parasitemia was included as a log-transformed covariate. MaRNAV-1 status (χ² = 62.0, df = 38, p = 0.0082) and parasitemia (χ² = 137.9, df = 38, p < 0.0001) were significantly associated with global immune cell composition. A significant interaction between MaRNAV-1 status and parasitemia was also observed (χ² = 36.3, df = 19, p = 0.0096).

| Predictors | χ2 | df | p_value |
| --- | --- | --- | --- |
| MaRNAV-1 status (presence/absence) | 62.0 | 38 | 0.0082 |
| log(Parasitemia) | 137.9 | 38 | <0.0001 |
| MaRNAV-1 status: log(Parasitemia) | 36.3 | 19 | 0.0096 |

**Supp. Table 5. Monoclonal antibody panel for flow cytometry (innate and adaptive immune panels).**

| **Panel** | **Fluorochromes** | **Markers** | **Surface/Chemokine markers** | **Staining °C** | **Vol/test** | **Clones** | **Catalog N** | **company** |
| --- | --- | --- | --- | --- | --- | --- | --- | --- |
| Innate Panel | BUV395 | CD3 | Surface | 4 °C | 1 μL | SK7 | 564001 | BD |
|  | BUV737 | CCR7 | Chemokine | 37 °C | 1 μL | 2-L1-A | 749676 | BD |
|  | BV421 | CD86 | Surface | 4 °C | 1 μL | IT2.2 | 305426 | Biolegend |
|  | BV510 | Viability | Surface | 4 °C | 1 μL | - | 423101 | Biolegend |
|  | BV605 | CD38 | Surface | 4 °C | 1 μL | H/T2 | 303532 | Biolegend |
|  | BV650 | CD11c | Surface | 4 °C | 1 μL | Bu15 | 337238 | Biolegend |
|  | BV786 | CD69 | Surface | 4 °C | 1 μL | FN50 | 310932 | Biolegend |
|  | AF488 | HLA-DR | Surface | 4 °C | 1 μL | L243 | 307620 | Biolegend |
|  | PE | CD16 | Surface | 4 °C | 1 μL | 3G8 | 560995 | BD |
|  | PE-Dazzle | CD56 | Surface | 4 °C | 1 μL | QA17A16 | 392410 | Biolegend |
|  | PE Cy7 | CD123 | Surface | 4 °C | 1 μL | 7G3 | 560826 | BD |
|  | APC | CD14 | Surface | 4 °C | 1 μL | 63D3 | 367118 | Biolegend |
|  | APC-Fire750 | CD83 | Surface | 4 °C | 1 μL | HB15e | 305331 | Biolegend |
| Adaptive Panel | BUV395 | CD3 | Surface | 4 °C | 1 μL | SK7 | 564001 | BD |
|  | BUV496 | CD4 | Surface | 4 °C | 1 μL | *SK3* | 612936 | BD |
|  | BUV737 | CCR7 | Surface | 37 °C | 1 μL | 2-L1-A | 749676 | BD |
|  | BV421 | CXCR3 | Surface | 37 °C | 1 μL | G025H7 | 353716 | Biolegend |
|  | BV510 | Viability | Surface | 4 °C | 1 μL | - | 423101 | Biolegend |
|  | BV605 | TIM3 | Surface | 4 °C | 1 μL | F38-2E2 | 345018 | Biolegend |
|  | BV650 | CCR4 | Surface | 37 °C | 1 μL | 1G1 | 744140 | BD |
|  | BV711 | IgD | Surface | 4 °C | 1 μL | IA6-2 | 740794 | BD |
|  | BV786 | CD138 | Surface | 4 °C | 1 μL | MI 15 | 356538 | Biolegend |
|  | BB515 | CCR6 | Surface | 37 °C | 1 μL | 11A9 | 564479 | BD |
|  | PercP cy5.5 | CD27 | Surface | 4 °C | 1 μL | M-T271 | 356408 | Biolegend |
|  | PE-dazzle | PD-1 | Surface | 4 °C | 1 μL | EH12.2H7 | 329940 | Biolegend |
|  | PE Cy7 | CD19 | Surface | 4 °C | 1 μL | HIB19 | 302216 | Biolegend |
|  | APC | HLA-DR | Surface | 4 °C | 1 μL | LN3 | 327021 | Biolegend |
|  | AF700 | CD8 | Surface | 4 °C | 1 μL | SK1 | 344724 | Biolegend |
|  | APC-Cy7 | CD45RO | Surface | 4 °C | 1 μL | UCHL1 | 304228 | Biolegend |

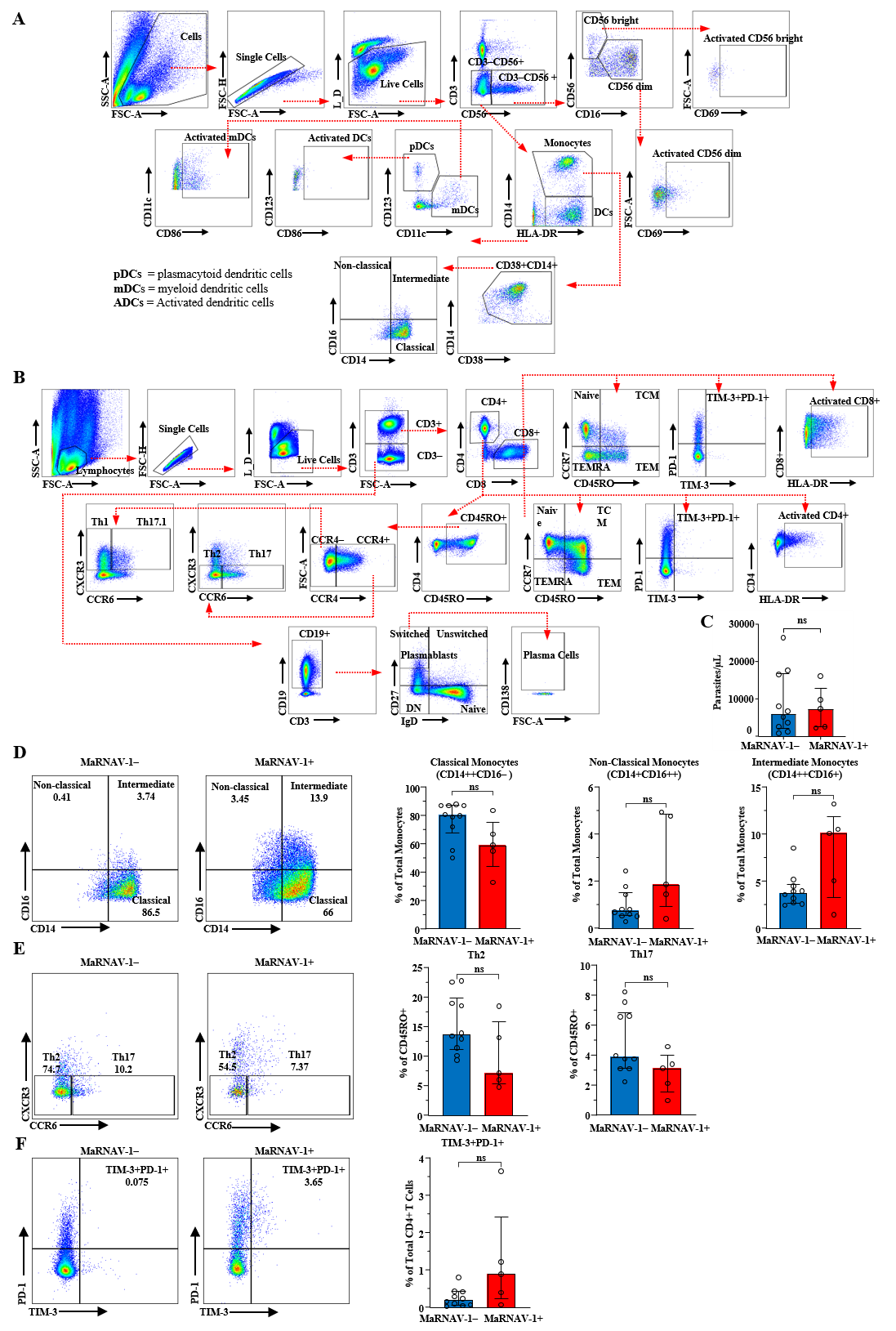

**Supp. Fig. 10**. **Flow cytometry gating strategy and immune cell subset proportions in *P. vivax*-infected patients stratified by MaRNAV-1 infection status (n=10 MaRNAV-1-negative and n=5 MaRNAV-1-positive)**. (**A**) Gating strategies to identify innate immune cell populations and (**B**) to identify adaptive immune cell populations in PBMCs. **C**. No significant difference in parasitemia between MaRNAV-1- negative and positive samples (p > 0.999). **D.** Proportion of monocytes in three subsets, including classical monocytes (CD14++CD16-), non-classical monocytes (CD14+CD16++), and intermediate monocytes (CD14++CD16+), in *P. vivax*-infected patients. **E**. Proportion of Th2 and Th17 out of CD45RO+ T cells. Th2 responses tend to be decreased in MaRNAV-1-positive samples, although the difference did not reach statistical significance (p=0.0753), while there is no significant difference in Th17 responses between the two groups (p=0.215). **F.** Proportion of TIM-3+PD-1+ CD4+ T cells out of total CD4+ T cells. TIM-3+PD-1+ CD4+ T cells tended to be higher in MaRNAV-1 positive samples (p= 0.0992).

**Supp. Table. 6**. **Association between MaRNAV-1 infection status and plasma cytokine levels in *P. vivax*-infected patients.** Associations were assessed using linear regression models that included log-transformed parasitemia and MaRNAV-1 infection status (present vs absent) as covariates (n = 77; 60 virus-positive and 17 virus-negative samples). MaRNAV-1 infection status was not significantly associated with cytokine levels after correction for multiple testing. In contrast, parasitemia was significantly associated with multiple cytokines (FDR-adjusted p < 0.05).

| Outcome | Predictor | Estimate | CI_lower | CI_upper | Chisq | df | p_value | p_FDR |
| --- | --- | --- | --- | --- | --- | --- | --- | --- |
| FGF_basic | log(Parasitemia) | 0.25 | 0.01 | 0.49 | 7.07 | 1 | 0.0389 | 0.0665 |
|  | MaRNAV-1_status | -0.07 | -0.76 | 0.63 | 0.06 | 1 | 0.8481 | 0.9704 |
| G.CSF | log(Parasitemia) | 0.26 | -0.01 | 0.53 | 7.62 | 1 | 0.0569 | 0.0754 |
|  | MaRNAV-1_status | -0.02 | -0.81 | 0.76 | 0.01 | 1 | 0.9530 | 0.9704 |
| IL-10 | log(Parasitemia) | 0.96 | 0.53 | 1.38 | 102.51 | 1 | 0.0000 | 0.0004 |
|  | MaRNAV-1_status | 0.24 | -1.00 | 1.48 | 0.75 | 1 | 0.7030 | 0.9704 |
| IL-13 | log(Parasitemia) | 0.36 | 0.01 | 0.70 | 14.14 | 1 | 0.0413 | 0.0665 |
|  | MaRNAV-1_status | 0.11 | -0.88 | 1.11 | 0.17 | 1 | 0.8220 | 0.9704 |
| IL-6 | log(Parasitemia) | 1.10 | 0.54 | 1.67 | 136.49 | 1 | 0.0002 | 0.0015 |
|  | MaRNAV-1_status | -0.33 | -1.98 | 1.33 | 1.42 | 1 | 0.6936 | 0.9704 |
| IL-12 | log(Parasitemia) | 0.26 | 0.04 | 0.48 | 7.77 | 1 | 0.0202 | 0.0437 |
|  | MaRNAV-1_status | 0.33 | -0.31 | 0.98 | 1.44 | 1 | 0.3090 | 0.9704 |
| Eotaxin | log(Parasitemia) | 0.40 | -0.01 | 0.81 | 17.90 | 1 | 0.0580 | 0.0754 |
|  | MaRNAV-1_status | -0.04 | -1.24 | 1.17 | 0.02 | 1 | 0.9527 | 0.9704 |
| MIP1a | log(Parasitemia) | 0.36 | 0.10 | 0.63 | 14.90 | 1 | 0.0078 | 0.0183 |
|  | MaRNAV-1_status | -0.05 | -0.83 | 0.72 | 0.04 | 1 | 0.8927 | 0.9704 |
| GMCSF | log(Parasitemia) | 0.47 | 0.01 | 0.93 | 24.76 | 1 | 0.0435 | 0.0665 |
|  | MaRNAV-1_status | -0.13 | -1.46 | 1.20 | 0.22 | 1 | 0.8470 | 0.9704 |
| MIP.1b | log(Parasitemia) | 0.45 | 0.20 | 0.70 | 22.64 | 1 | 0.0006 | 0.0033 |
|  | MaRNAV-1_status | -0.04 | -0.78 | 0.69 | 0.02 | 1 | 0.9061 | 0.9704 |
| MCP.1 | log(Parasitemia) | 0.72 | 0.37 | 1.08 | 58.59 | 1 | 0.0001 | 0.0011 |
|  | MaRNAV-1_status | 0.02 | -1.02 | 1.06 | 0.00 | 1 | 0.9704 | 0.9704 |
| IL-15 | log(Parasitemia) | 0.37 | -0.05 | 0.80 | 15.41 | 1 | 0.0865 | 0.1071 |
|  | MaRNAV-1_status | 0.24 | -1.00 | 1.48 | 0.73 | 1 | 0.7057 | 0.9704 |
| EGF | log(Parasitemia) | 0.19 | -0.16 | 0.55 | 4.18 | 1 | 0.2850 | 0.2850 |
|  | MaRNAV-1_status | 0.37 | -0.67 | 1.41 | 1.80 | 1 | 0.4816 | 0.9704 |
| HGF | log(Parasitemia) | 0.23 | 0.03 | 0.43 | 5.87 | 1 | 0.0270 | 0.0502 |
|  | MaRNAV-1_status | 0.19 | -0.40 | 0.78 | 0.47 | 1 | 0.5230 | 0.9704 |
| VEGF | log(Parasitemia) | 0.56 | 0.24 | 0.87 | 34.50 | 1 | 0.0008 | 0.0035 |
|  | MaRNAV-1_status | 0.59 | -0.33 | 1.51 | 4.58 | 1 | 0.2074 | 0.9704 |
| IFN.g | log(Parasitemia) | 0.30 | -0.11 | 0.71 | 9.95 | 1 | 0.1506 | 0.1780 |
|  | MaRNAV-1_status | -0.62 | -1.81 | 0.57 | 5.05 | 1 | 0.3039 | 0.9704 |
| IFN.a | log(Parasitemia) | 0.19 | -0.08 | 0.45 | 3.86 | 1 | 0.1621 | 0.1800 |
|  | MaRNAV-1_status | -0.13 | -0.89 | 0.63 | 0.23 | 1 | 0.7327 | 0.9704 |
| IL1-RA | log(Parasitemia) | 0.48 | 0.19 | 0.77 | 25.65 | 1 | 0.0015 | 0.0054 |
|  | MaRNAV-1_status | 0.14 | -0.70 | 0.98 | 0.26 | 1 | 0.7400 | 0.9704 |
| TNF.a | log(Parasitemia) | 0.56 | 0.19 | 0.92 | 34.82 | 1 | 0.0031 | 0.0102 |
|  | MaRNAV-1_status | -0.28 | -1.34 | 0.78 | 1.05 | 1 | 0.5982 | 0.9704 |
| IL-2 | log(Parasitemia) | 0.35 | -0.19 | 0.90 | 14.02 | 1 | 0.1984 | 0.2063 |
|  | MaRNAV-1_status | 0.12 | -1.46 | 1.71 | 0.21 | 1 | 0.8757 | 0.9704 |
| IL-7 | log(Parasitemia) | 0.23 | 0.00 | 0.46 | 5.89 | 1 | 0.0549 | 0.0754 |
|  | MaRNAV-1_status | -0.05 | -0.73 | 0.64 | 0.03 | 1 | 0.8919 | 0.9704 |
| CXCL10 | log(Parasitemia) | 0.35 | 0.11 | 0.59 | 13.63 | 1 | 0.0056 | 0.0162 |
|  | MaRNAV-1_status | 0.05 | -0.66 | 0.76 | 0.03 | 1 | 0.8918 | 0.9704 |
| IL-2R | log(Parasitemia) | -0.44 | -1.06 | 0.18 | 12.16 | 1 | 0.0072 | 0.0183 |
|  | MaRNAV-1_status | -8.04 | -13.99 | -2.09 | 0.03 | 1 | 0.8874 | 0.9704 |
|  | log(Parasitemia):MaRNAV-1_status | 0.90 | 0.24 | 1.57 | 11.56 | 1 | 0.0087 | 0.0087 |
| MIG | log(Parasitemia) | 0.23 | 0.03 | 0.43 | 5.90 | 1 | 0.0270 | 0.0502 |
|  | MaRNAV-1_status | 0.21 | -0.38 | 0.80 | 0.58 | 1 | 0.4826 | 0.9704 |
| IL-4 | log(Parasitemia) | 0.24 | -0.10 | 0.59 | 6.52 | 1 | 0.1662 | 0.1800 |
|  | MaRNAV-1_status | 0.04 | -0.97 | 1.04 | 0.02 | 1 | 0.9417 | 0.9704 |
| IL-8 | log(Parasitemia) | 0.78 | 0.44 | 1.12 | 67.93 | 1 | 0.0000 | 0.0004 |
|  | MaRNAV-1_status | -0.29 | -1.28 | 0.71 | 1.08 | 1 | 0.5685 | 0.9704 |

**Supp. Table 7. Non-linear associations between MaRNAV-1 load and cytokine levels in MaRNAV-1-positive *P. vivax*-infected patients.** Associations were assessed using generalized additive models (GAMs) including smooth terms for log-transformed MaRNAV-1 load and parasitemia (n = 60). Parasitemia was significantly associated with multiple cytokines (FDR-adjusted p < 0.05). MaRNAV-1 load was also significantly associated with several cytokines, including IFN-γ, IL-10, IL-1RA, IL-6, IP-10, and VEGF (FDR-adjusted p < 0.05), indicating non-linear relationships between viral load and host immune activation. A significant interaction between MaRNAV-1 load and parasitemia was observed for IL-6 (FDR-adjusted p = 0.011), suggesting that the effect of viral load on IL-6 levels depends on parasite density.

| Outcome | Predictor | edf | F | p_value | p_FDR |
| --- | --- | --- | --- | --- | --- |
| EGF | s(log(Parasitemia)) | 1.00 | 2.168 | 0.146 | 0.152 |
|  | s(log(MaRNAV-1 load)) | 1.00 | 0.307 | 0.582 | 0.630 |
| Eotaxin | s(log(Parasitemia)) | 1.00 | 4.097 | 0.048 | 0.062 |
|  | s(log(MaRNAV-1 load)) | 2.94 | 1.152 | 0.278 | 0.474 |
| FGF_basic | s(log(Parasitemia)) | 1.00 | 4.906 | 0.031 | 0.044 |
|  | s(log(MaRNAV-1 load)) | 1.00 | 1.310 | 0.257 | 0.474 |
| G.CSF | s(log(Parasitemia)) | 1.00 | 5.047 | 0.029 | 0.044 |
|  | s(log(MaRNAV-1 load)) | 1.11 | 1.000 | 0.284 | 0.474 |
| GMCSF | s(log(Parasitemia)) | 1.00 | 4.912 | 0.031 | 0.044 |
|  | s(log(MaRNAV-1 load)) | 1.00 | 0.170 | 0.681 | 0.709 |
| HGF | s(log(Parasitemia)) | 1.15 | 5.685 | 0.011 | 0.024 |
|  | s(log(MaRNAV-1 load)) | 1.26 | 0.705 | 0.357 | 0.492 |
| IFN.α | s(log(Parasitemia)) | 1.00 | 3.131 | 0.082 | 0.097 |
|  | s(log(MaRNAV-1 load)) | 1.00 | 1.051 | 0.310 | 0.474 |
| IFN.γ | s(log(Parasitemia)) | 1.00 | 2.877 | 0.095 | 0.104 |
|  | s(log(MaRNAV-1 load)) | 1.67 | 4.907 | 0.011 | 0.047 |
| IL-13 | s(log(Parasitemia)) | 1.00 | 5.176 | 0.027 | 0.044 |
|  | s(log(MaRNAV-1 load)) | 1.46 | 0.508 | 0.476 | 0.563 |
| IL-10 | s(log(Parasitemia)) | 1.00 | 22.145 | 0.000 | 0.000 |
|  | s(log(MaRNAV-1 load)) | 2.43 | 5.103 | 0.003 | 0.027 |
| IL-12 | s(log(Parasitemia)) | 1.00 | 6.696 | 0.012 | 0.025 |
|  | s(log(MaRNAV-1 load)) | 2.36 | 1.514 | 0.182 | 0.474 |
| IL-15 | s(log(Parasitemia)) | 1.41 | 2.059 | 0.096 | 0.104 |
|  | s(log(MaRNAV-1 load)) | 1.00 | 1.467 | 0.231 | 0.474 |
| IL-1RA | s(log(Parasitemia)) | 1.22 | 8.063 | 0.002 | 0.008 |
|  | s(log(MaRNAV-1 load)) | 5.69 | 3.501 | 0.004 | 0.027 |
| IL-2 | s(log(Parasitemia)) | 1.00 | 3.147 | 0.081 | 0.097 |
|  | s(log(MaRNAV-1 load)) | 1.00 | 1.489 | 0.227 | 0.474 |
| IL-2R | s(log(Parasitemia)) | 1.00 | 11.740 | 0.001 | 0.005 |
|  | s(log(MaRNAV-1 load)) | 1.50 | 0.913 | 0.310 | 0.474 |
| IL-4 | s(log(Parasitemia)) | 1.00 | 1.517 | 0.223 | 0.223 |
|  | s(log(MaRNAV-1 load)) | 1.00 | 0.330 | 0.568 | 0.630 |
| IL-6 | s(log(Parasitemia)) | 2.23 | 9.789 | 0.000 | 0.001 |
|  | s(log(MaRNAV-1 load)) | 2.42 | 8.110 | 0.000 | 0.007 |
|  | ti(log(Parasitemia),log(MaRNAV-1 load)) | 9.24 | 2.630 | 0.011 | 0.011 |
| IL-7 | s(log(Parasitemia)) | 1.00 | 5.800 | 0.019 | 0.036 |
|  | s(log(MaRNAV-1 load)) | 1.00 | 0.551 | 0.461 | 0.563 |
| IL-8 | s(log(Parasitemia)) | 2.20 | 9.807 | 0.000 | 0.001 |
|  | s(log(MaRNAV-1 load)) | 1.70 | 0.997 | 0.359 | 0.492 |
| CXCL10 | s(log(Parasitemia)) | 1.64 | 5.369 | 0.007 | 0.018 |
|  | s(log(MaRNAV-1 load)) | 2.14 | 4.434 | 0.009 | 0.045 |
| MCP.1 | s(log(Parasitemia)) | 2.19 | 8.130 | 0.000 | 0.002 |
|  | s(log(MaRNAV-1 load)) | 2.72 | 1.585 | 0.185 | 0.474 |
| MIG | s(log(Parasitemia)) | 1.00 | 4.830 | 0.032 | 0.044 |
|  | s(log(MaRNAV-1 load)) | 1.14 | 2.400 | 0.094 | 0.307 |
| MIP.1b | s(log(Parasitemia)) | 1.43 | 6.160 | 0.004 | 0.011 |
|  | s(log(MaRNAV-1 load)) | 2.72 | 0.833 | 0.475 | 0.563 |
| MIP1a | s(log(Parasitemia)) | 1.00 | 8.925 | 0.004 | 0.011 |
|  | s(log(MaRNAV-1 load)) | 1.00 | 0.106 | 0.747 | 0.747 |
| TNF.α | s(log(Parasitemia)) | 1.00 | 11.723 | 0.001 | 0.005 |
|  | s(log(MaRNAV-1 load)) | 3.52 | 2.102 | 0.080 | 0.298 |
| VEGF | s(log(Parasitemia)) | 1.00 | 9.929 | 0.003 | 0.008 |
|  | s(log(MaRNAV-1 load)) | 2.38 | 6.832 | 0.001 | 0.008 |

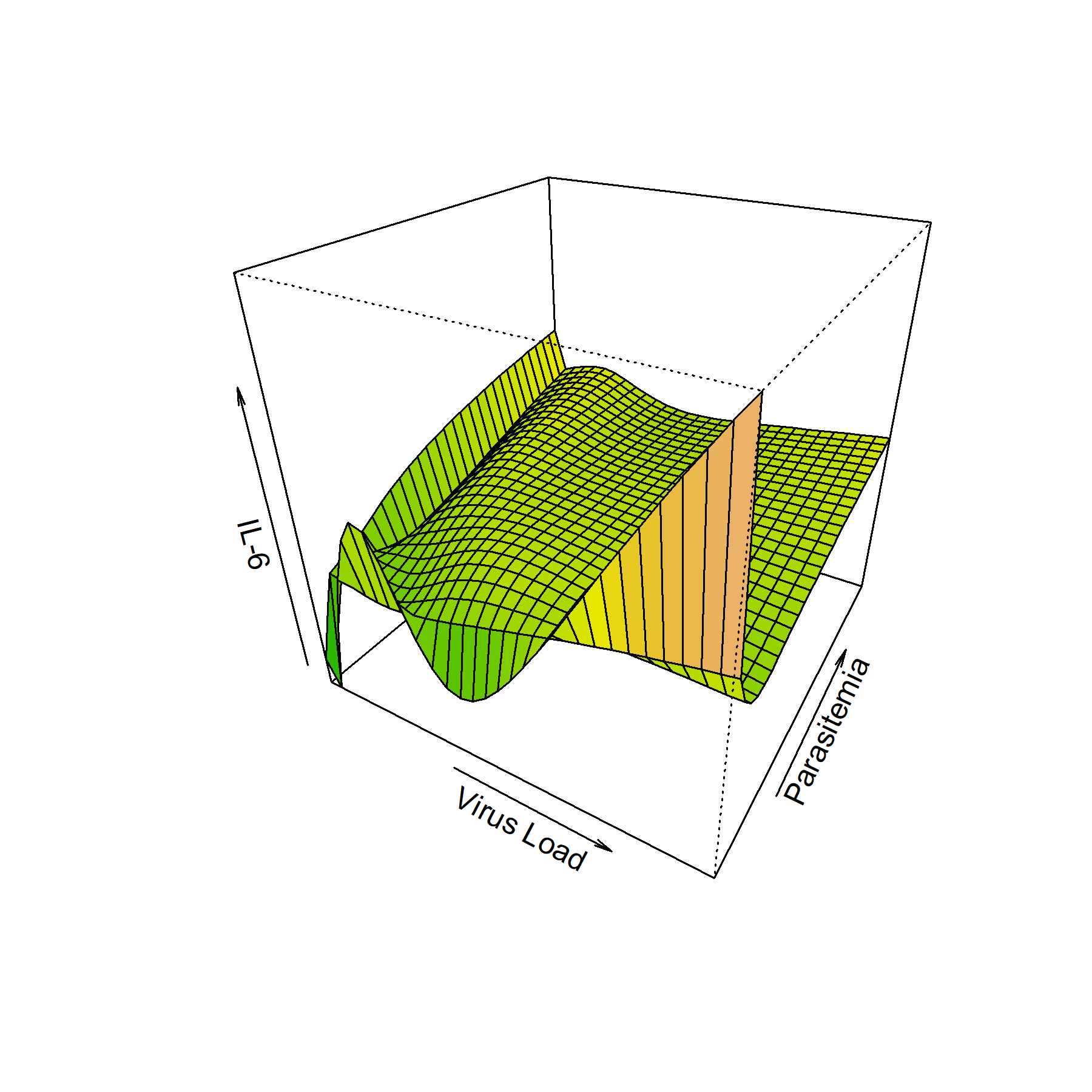

**Supp. Fig. 11. Interaction between MaRNAV-1 load and parasitemia on IL-6 levels**. The three-dimensional surface shows the GAM-fitted values, including an interaction term, illustrating how the combined effects of viral load and parasitemia influence IL-6 concentrations. Axes represent MaRNAV-1 load, parasitemia, and IL-6.

**Supp. Table 8. Primers used in this study**

| Primer Name | Sequence (5'- 3') | Application |
| --- | --- | --- |
| Pv_18S_F | ATGGCTATGACGGGTAACGG | *P. vivax* reference gene for RT-qPCR |
| Pv_18S_R | GCCTGCTGCCTTCCTTAGAT |  |
| PvnarS1_F | AAGGACGCGACGAATGCTC | S1 detection/quantification by RT-qPCR |
| PvnarS1_R | CGAAGTCAACGTAGGGATTTGG |  |
| PvnarS2_F | TCAATGTACCTTATGGCGTCCG | S2 detection/quantification by RT-qPCR |
| PvnarS2_R | GACAGGGCCTTAGGTTGGTC |  |
| Human_RPS18_Fw | ATGCAGAATCCACGCCAGTA | Human reference gene for RT-qPCR |
| Human_RPS18_Rev | CCAGACCATTGGCTAGGACC |  |
| Pvs25_F | GGCAAAGTCCCCAATCCAGA | Female gametocytes detection/quantification by RT-qPCR |
| Pvs25_R | GCCTTCCATACACTGGCACT |  |
| Pvs47_F | CGATACATATCGGCTCCAGCA | Male gametocytes detection/quantification by RT-qPCR |
| Pvs47_R | TCACCCCGTTGACGTCTTTT |  |
